# Mutagenesis study of a Bacteriophytochrome – insights for the development of labels for optical imaging

**DOI:** 10.64898/2026.03.03.708993

**Authors:** Azeem Mohammed, David Wiesner, Anna Dachs, Elias Wippermann, Simon Göllner, Stephanie Willeit, Uwe Klemm, Selina Schulz, Bjarne Perleberg, John J. Galligan, Elsa Rodrigues, Isabella Rembeck, Axel Duerkop, Vasilis Ntziachristos, Till Rudack, Andre C. Stiel

## Abstract

Bacteriophytochromes (BphPs) find increasing interest as near-infrared (NIR) labels for imaging. Applications range from whole animal imaging to cell-to super-resolution microscopy. Here we present a comprehensive study of a BphP from *Rhizobium etli* (ReBphP) allowing mutant-based insights into BphP photophysics. This is complemented by QM/MM-optimized deep-learning structure predictions of ReBphP and variants in their photoswitched states, rationalizing the effects of variants. Pertaining to imaging, based on our study we identify a bright and far red-shifted BphP for imaging in mammalian cells as well as a fluorescent photoswitching BphP. Utilizing the latter, we introduce photoswitching fluorescence background suppression for *in vivo* whole animal fluorescence imaging achieving higher contrast over background than possible utilizing non-switching labels.

## Introduction

Bacteriophytochromes (BphPs), a subclass of the larger family of phytochromes, are light-sensing proteins found in various bacteria. In their biological context, BphPs act as light sensors, triggering a variety of downstream functions^1^. In recent years, BphPs have gained prominence for their use in optogenetic applications, but also as labels in optical imaging^2^. The absorption and emission of BphPs in the near-infrared (NIR, >650 nm) extends the wavelength palette of genetically encodable labels for imaging beyond the family of green fluorescent protein (GFP)-like proteins (<650 nm). In contrast to the autocatalytically formed chromophore of GFP-like proteins, BphPs require covalent attachment of biliverdin (BV) to a reactive cysteine residue in the chromophore pocket. However, BV – a byproduct of the heme catabolism – is readily available in most mammalian cell types, hence BphPs can be considered fully genetically encoded labels. A key function of BphPs, owing to their biological function as light-sensors, is that they can be photoswitched by specific wavelengths between the conformational and photophysically distinct red (P_r_) and far-red (P_fr_) state (**Figure 1a and b**). The basis of this mechanism is a light-induced *cis*/*trans* (*ZZZssa*/*ZZEssa*) isomerization of the D-ring of the BV with a complex influence of the non-covalent interactions of the BV surrounding amino acids^3^.

**Figure 1.**
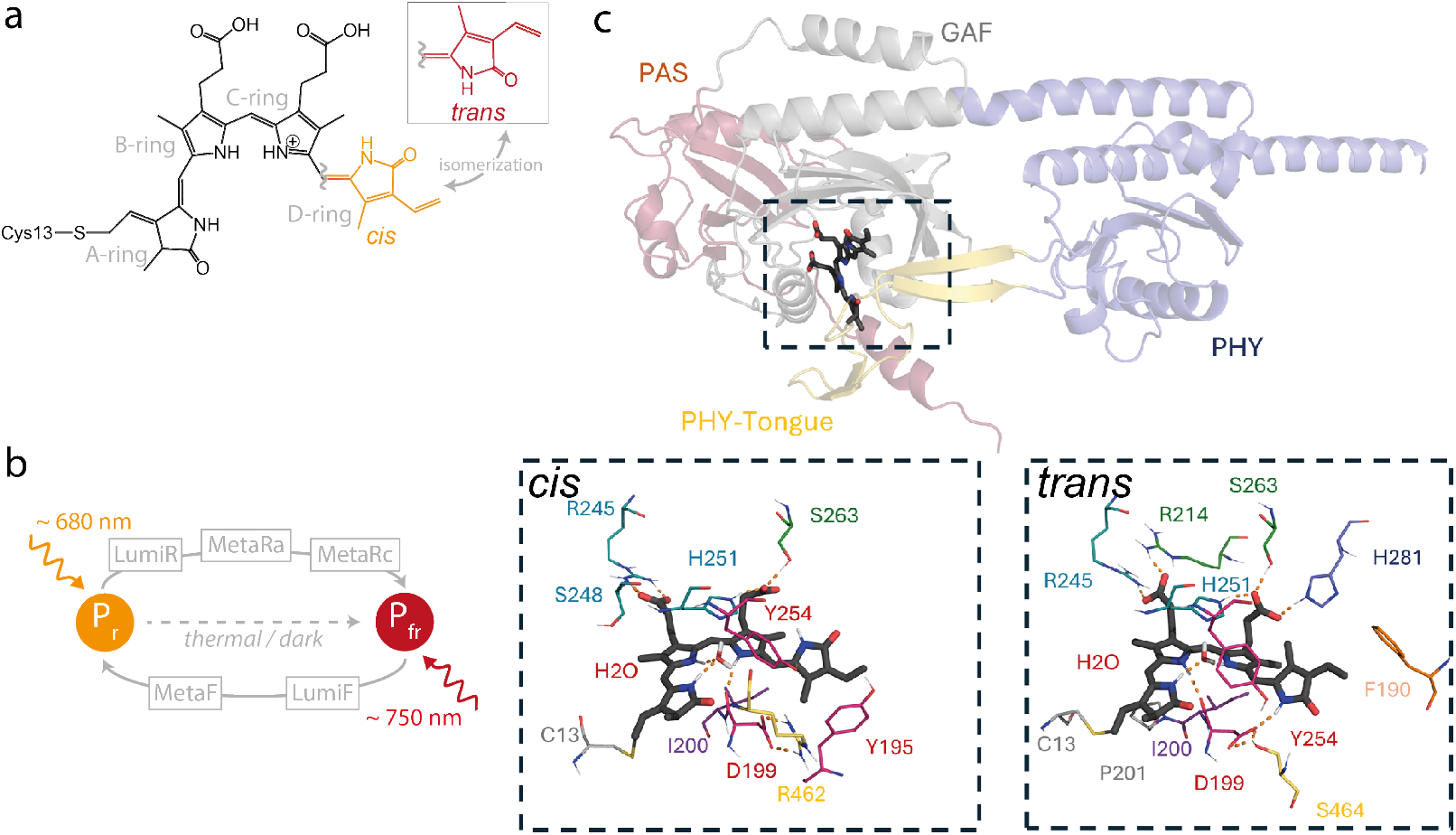
**a** Biliverdin (BV) chromophore in its *cis* and *trans* isomeric forms. **b** Photocycle of bathy BphPs with major intermediates indicated. **c** Model of ReBphP-PCM’s chromophore binding pocket (rectangle inset top left) in the P_r_ (BV *cis*, left) and P_fr_ (*trans*, right) state. Color coding of residues according to spatial or functional grouping (see **Figure 2a**).

Native BphPs are only poorly fluorescent largely due to the chromophore flexibility facilitating switching but disfavoring radiative deexcitation^4^, but also due to excited state proton transfer (ESPT)^5,6^ and ground state heterogeneities^5,7^. Hence, most engineering efforts so far have focused on increasing fluorescence by locking the chromophore in its *cis* isomeric state and adding fluorescence favoring mutations around the D-ring. This yielded relatively bright fluorescent variants now in use for microscopy and whole animal imaging (e.g.^8–14^). Conversely, BphPs have recently gained attention as photoswitching labels in optoacoustic (OA, also called photoacoustic) imaging, a technique that achieves high-resolution deep tissue imaging by specifically exciting the target with tissue penetrating NIR light and detecting an ultrasound readout instead of light^15,16^. In OA imaging the low fluorescence of native BphPs is beneficial since it increases the non-radiative relaxation yielding the OA signal. The photoswitching of BphPs on the other hand can be used to introduce a temporal modulation of the signal. This can help separating the signal of labeled cells from background by locked-in detection^17^, significantly improving the visualization of smaller cell populations in live animals^18–22^.

Protein engineering efforts tailoring BphPs as imaging labels rely on addressing the interplay of the BV chromophore and the protein matrix^4^. This engineering focuses on the GAF domain as hosting most BV protein interactions. In fact, the majority of BphPs used in imaging have been truncated to the PAS-GAF-PHY domains (PCM, photosensory core module) or even only PAS-GAF for fluorescent variants (CBD, chromophore binding domain). So far, extensive engineering has focused on *D. radiodurans* BphP (iFPs^23^, WiPHY^10^, SNIFP^11^), *R. pseudomonas* BphPP1 (iRFP670s, 703s, 709s, mIFP2^8^, mRhubarbs^13^), BphP2 (iRFP682s, 713s, 720s^8^) and *R. pseudomonas BphP6* (iRFP670s, 702s^8^). In this work, we present a systematic mutagenesis study of a BphP from *Rhizobium etli* (ReBphPs). So far, ReBphP has been studied only very little^24^ but shows excellent disposition for the engineering of imaging labels as we show in this work. We performed in depth analysis of about 90 variants together with structural models of selected variants in their respective assumed P_r_ and P_fr_ state providing a comprehensive view of ReBphP photophysics. This allowed us to identify variants beneficial for imaging: a bright far red-shifted BphP and a fluorescent photoswitching protein in the NIR. Using the latter, we demonstrate a photoswitching approach for autofluorescence background suppression in fluorescence whole animal imaging allowing a contrast-to-background increase of more than 20x compared to conventional genetically encoded labels.

## Results & Discussion

The positions for mutagenesis have been chosen based on intuition and prior BphP mutagenesis studies in a semi-rational scouting fashion. This led to overrepresentation of mutations at positions where initial variants showed effects, especially near the D-ring, which is primary involved in the isomerization of the BV chromophore. Nonetheless, the variant set yields an ample coverage of large parts of the BV pocket of *Re*BphP (variant overview **Suppl. Table 1**). Beyond single mutations, also combinatorial mutations have been introduced. Most mutations are introduced in ReBphP PCM with a subset exploring the effect of truncation to ReBphP CBD (PAS-GAF) and mutations therein. For all variants absorption spectra have been recorded after 780 and 660 nm illumination directly after purification (P_fr_ → P_r_: 780 ± 15 nm or P_r_ → P_fr_: 660 ± 10 nm, unless stated differently; absorption and fluorescence spectra for all variants **Suppl. Figure 1**). Additionally, for some variants, absorption spectra of the thermal equilibrium state, fluorescence spectra after 680 nm excitation and photoswitching kinetics for the P_fr_ → P_r_ transitions have been recorded. The interactions and residue arrangements within the binding pocket were analyzed using refined models of the ReBphP-PCM protein in its *cis* (Pr) and *trans* (Pfr) states. The models were created by a chimeric deep-learning structure prediction and homology modeling approach and further refined via hybrid quantum mechanics / molecular mechanics (see methods and **Suppl. Text 1** for details, **Figure 1c and Suppl. Figure 2**).

### Protein expression and chromophorylation

We report a protein expression score based on purified protein per culture volume (**Suppl. Table 1**). This expression score gives an indication which positions and substitutions affect the overall protein integrity more severely than others but does not attempt to exactly quantify expression levels and folding efficiency. While several variants showed reduced expression, suggesting poorly folded proteins, there was no variant that did not express or could not be purified at all (0/70 single mutants). This robustness aligns with earlier findings on DrBphP^25^. Using a double promoter vector, we co-expressed BphP together with heme oxygenase HO1 (*Nostoc sp*.) to facilitate chromophorylation in the bacteria. The effectivity of chromophorylation, i.e. the functional attachment of the BV chromophore to the protein, can be approximated by the ratio between the Soret and 280 nm peak with the latter, normalized by extinction coefficient of the protein, indicating protein concentration. Even under standardized conditions the chromophorylation in repeated expressions of ReBphP-PCM shows a large variation of 31% (N=13, 94 ± 30, **Suppl. Figure 3a**). Due to this variance in chromophorylation, we group the effect of mutations on chromophorylation only tentatively into a chromophorylation like ReBphP-PCM, slightly better or worse (**Suppl. Figure 3b**). Most of the single variants show a chromophorylation like ReBphP-PCM or worse; however, only 5 of 70 single variants showed a complete lack of measurable chromophorylation. For some variants reduced chromophorylation can be expected, for example, R254A likely affects the stabilization of the B-ring propionate side chain. More generally, variants like S263A can affect the electrostatic complementarity of the binding pocket, like such mutations in DrBphP^25^.

Other mutations like F190W might result in steric obstruction of access to the binding pocket or reduced shape complementarity. Variants G261R showed a slightly improved chromophorylation, an effect already described for DrBphP derived SNIFP^11^. Beyond that, most deletion variants of the PHY domain (Δ_PHY_) improved chromophorylation. Likely due to the improved accessibility of the chromophore binding pocket without the PHY tongue present. In general, we observe a slight correlation between folding (assumed via expression yield) and chromophorylation (**Suppl. Figure 4**). Along those lines, the mutations that do not show any measurable chromophorylation show only poor or mediocre expression, too (M166V, Y195T, R245L, R245I, R245E and I250N). Hence, structures with corrupt or slow folding might impede chromophorylation.

### Spectral characteristics of ReBphP-PCM variants

As other BphPs ReBphP-PCM shows two fundamental spectral states: P_r_ and P_fr_. The P_r_ state absorption spectrum is characterized by a dominant peak at ~700 nm with a blue shoulder at ~650 nm, while the P_fr_ state spectrum still features considerable contributions from the 700 nm peak, but also a prominent new peak at ~760 nm. We refer to these two peaks by the names 700- and 760-band, although the actual absorption is not always centered exactly at those wavelengths. P_r_ and P_fr_ are the photoproducts of illumination in the 760- and 700-band, respectively. The 700-band is attributed to the *cis* chromophore of BV, while the 760-band is associated with the *trans* chromophore. The substantial remaining 700-band in P_fr_ likely stems from mixed species in P_r_, whereof not all are photoswitchable resulting in the incomplete photochromism. The species mainly differ by their hydrogen bonding networks to the C-ring propionate and the D-ring carbonyl group^26^. The spectral characteristics are highly reproducible among independent ReBphP-PCM purifications (N=13, **Suppl. Figure 5 and 6**). The variants explored in this study introduce a range of spectral changes (**Figure 2**); to better capture the different characteristics we clustered the absorption spectra after illumination in the 760-band (P_fr_ → P_r,_ termed P_r_ clusters) and 700-band (P_r_ → P_fr,_ P_fr_ clusters) among themselves. The clustering only considers the spectral shape indifferent of the absolute absorption and peak wavelength position. Most of the variants show the behavior of ReBphP-PCM in all spectral aspects (P_r_-cluster 1 and P_fr_-cluster 1). Among those ReBphP-PCM-like variants, the only small differences are in the relation of 760-to 700-band in P_fr_. Variants with such largely unchanged spectra also include rather drastic substitutions, for example at position R245, which heavily affect chromophorylation but seemingly not the chromophores’ spectral properties.

**Figure 2.**
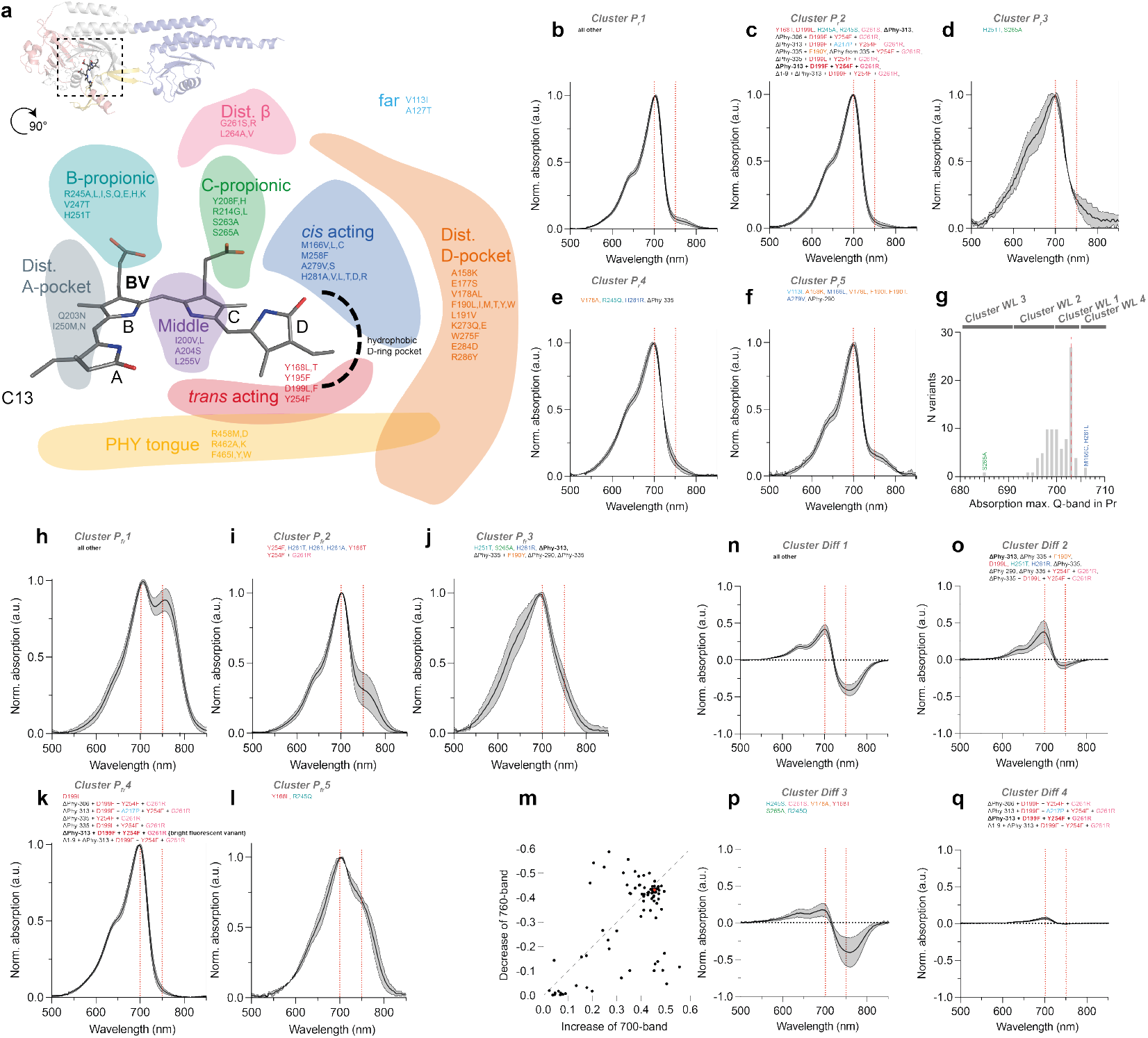
Spectral characterizations of ReBphP-PCM variants. **a** Outtake of the chromophore (BV, sticks) surrounding (boxed area inset). Residue positions that have been probed by mutagenesis are indicated and color-coded based on their position to the chromophore. **b-f** Absorption spectra after illumination in the 760-band (P_fr_ → P_r_) clustered by spectral shape indifferent of WL position or height. Mutants belonging to the respective cluster are indicated above in color coding as in a and combinatorial variants or truncations in black. Most variants show a ReBphP-PCM wildtype like behavior (cluster P_r_ 1) and are not specifically mentioned. **g** Frequency histogram of absorption peak wavelengths of variants in the P_r_ state. **h-l** Absorption spectra after illumination in the 700-band (P_r_ → P_fr_) clustered by spectral shape. Representation as in b-f. **m** Relation of the change in the 700- and 760-band upon illumination in the 760 nm band. ReBphP-PCM indicated in red. **n-q** Difference spectra (P_r_-P_fr_) clustered by shape. Representation as in b-f. For all three spectral analyses the appearance and hence variance of an “all-wildtype cluster” can be found in **Suppl. Figure 5**.

### Spectra after illumination in the 760-band (P_r_ spectra)

The clustering of spectra after illumination in the 760-band (P_fr_ → P_r_), i.e. the proteins in the P_r_ state (P_r_ clusters) show (*i*) different prominence of a blue shoulder (~650 nm). The exact determinants of the blue shoulder are not fully clarified, ranging from a general ground state heterogeneity of the BV^27,28^ or differences in protonation of the BV or nearby residues^29,30^. In our work, the blue shoulder is most pronounced in P_r_-cluster 3 and 4 with residues that can affect the charge distribution or the hydrogen bonding network around the chromophore (e.g. Y195L or R245Q). (*ii*) A second difference among spectra of the P_r_ states is the remaining contribution of the 760-band even after prolonged 760 nm illumination. Variants in cluster-P_r_ 5 show increased remaining 760-band contribution. Those variants seem to primarily loosen the packing in the hydrophobic pocket partially surrounding the D-ring. This potentially allows more P_fr_ state heterogeneity including a not switchable population. In contrast, the variants in cluster-P_r_ 2 with a truncated PHY-domain (Δ_PHY_), known to lock photoswitching^31^, are expectedly producing the cleanest P_r_ state spectra with no 760-band contribution whatsoever. *iii*) Regarding their peak wavelength (**Figure 2g**), most of the P_r_ state spectra show a maximum of the 700-band at 703 nm like ReBphP-PCM. However, several variants are blue-shifted compared to ReBphP-PCM with the most extreme shifts for variant S265A (>15 nm), likely due to a loss in polarity near the chromophore. Only variant M166C and H281L show a slight red shift (3 nm). The spectral shifts are in line with the general range of wildtype BphP showing absorption maxima around 700 nm and below, but not above, likely marking the maximum achievable red shift of absorption of the BV *cis*-isomer in a pocket of the GAF-fold^8^.

### Spectra after illumination in the 700-band (P_fr_ spectra)

(*i*) After illumination in the 700-band (P_r_ → P_fr_) native ReBphP-PCM and similar variants show spectra with a ratio between the 760- and 700 nm band of ~80% (cluster-P_fr_ 1, **Figure 2h-m**). However, several of the generated variants deviate from this ratio. The most drastic effect can be found for the well-known position D199^25,32,33^ which is a key player for the interaction network of the *trans* chromophore with Y254 and PHY-tongue S469 (cluster-P_fr_ 4). Variants D199L and D199F completely abolish the formation of a 760-band and hence the P_fr_ form similar to results in RpBphP ^34,35^, DrBphP^25^ and generally all BphP derivates engineered for high fluorescence brightness^31^. In DrBphP mutation Y176H also precluded *trans* formation^25^, we could not see a similarly strong effect for the corresponding position Y168 in ReBphP-PCM, since variants Y168L and T show incomplete, but existing 760 nm bands. Other variants which show only a reduction of the 760-band are for example Y254F (~40%, cluster-P_fr_ 2), which is known from engineered fluorescent BphP to destabilize the *trans* state^31^. (*ii*) A group of variants (cluster-P_fr_ 3) does not show a 760-band after illumination in the 700-band (P_r_ → P_fr_), but a clear broadening of the 700-band towards the red. Despite lacking a 760-band, variants like for example, Δ_PHY_313 can be readily switched back with 760 nm light (P_fr_ → P_r_). Spectral decomposition suggests that the photoswitched species is not a canonical P_fr_ (clear 760-band) but rather some intermediate with a more blue shifted absorption (**Suppl. Figure 7**). We observe similar broadening for variant H251T which aligns with similar findings for mutations in DrBphP (H260)^25^ and AtBphP1 (H250)^33^. Based on resonance Raman measurements those works suggest the species induced by illumination in the 700-band in those variants to be a Meta-R_c_ intermediate. A possible reason is a changed water molecule network due to the space freed up by the H251 mutations^36^. This aligns with what we find when comparing models of ReBphP-PCM and Δ_PHY_313 and in the *trans* state, the latter shows a distinctly different positioning of the C-ring propionate and the associated water molecule network (**Suppl. Figure 8**). Such water molecule network changes have been shown to influence state heterogeneity in DrBphP^26^. (*iii*) Lastly, a surprising finding is the low influence of H281 mutations. H281 is somewhat the opposing player to D199 since it interacts with the D-ring carbonyl stabilizing the *cis* form. For most mutations (H281AVLT and seemingly even H281D), we surprisingly see ReBphP-PCM-like behavior (Cluster-P_r_ and Cluster-P_fr_ 1) with only variants H281AVT showing a reduced 760-band after illumination in the 700-band (Cluster-P_fr_ 2) similar to H281N mutation spectra for DrBphP (H290N)^25^. The only influential mutation appears to be H281R with a broadening of the 700-band as described above (Cluster-P_fr_ 3).

### Photoswitching characteristics

The key feature of BphPs is the photoswitching of BV from the *trans* to *cis* isomer by illumination in the 760-band (P_fr_ → P_r_ transition) as well as the *cis* to *trans* isomerization by illumination in the 700-band (P_r_ → P_fr_). The effect of this photoswitching on the spectral characteristics of the variants differs resulting in altered difference spectra (P_r_ - P_fr_, **Figure 2m-q**). While ReBphP-PCM and most of the variants show a loss of the 700- and gain of the 760-band upon P_r_ → P_fr_ transition (cluster-Diff 1) a set of variants shows almost exclusively a change in the 700 nm absorption (cluster-Diff 2) or predominantly in the 760 nm absorption (cluster-Diff 3). Lastly, as described above, some variants show no photoswitching and photochromism and hence no spectral difference (cluster-Diff 4). The non-proportional change of variants (cluster-Diff 2 and 3) likely stems from multiple species in P_r_ not all equally photoswitchable and not all showing necessarily the same absorptivity or from different photostationary mixtures^37^ reachable depending on the switching propensity of P_r_ and P_fr_.

Additionally, we measured the bulk photoswitching time characteristics of the P_fr_ → P_r_ transition (**Figure 3a**). Most of the changes are minimal with only some variants showing clear slower bulk switching and even fever variants showing a minimal increase in photoswitching speed. The bulk switching speed is determined by the absorptivity of the band driving the transition in this case the 760 nm absorption of the P_fr_ state as well as the quantum yield for the transition. Several variants with slightly slower switching also show a reduced absorption of the 760 nm band which could be the primary reason for the slower switching (**Figure 3b**). However, many variants with a loss in 760 nm absorption show ReBphP-PCM switching kinetics or even faster switching, suggesting a more effective transition for those variants. The fact that the occurrence of faster variants is rare could be explained with the bulk switching of ReBphP-PCM being already fast – at least considerably faster than DrBphP or RpBphP1^22^.

**Figure 3.**
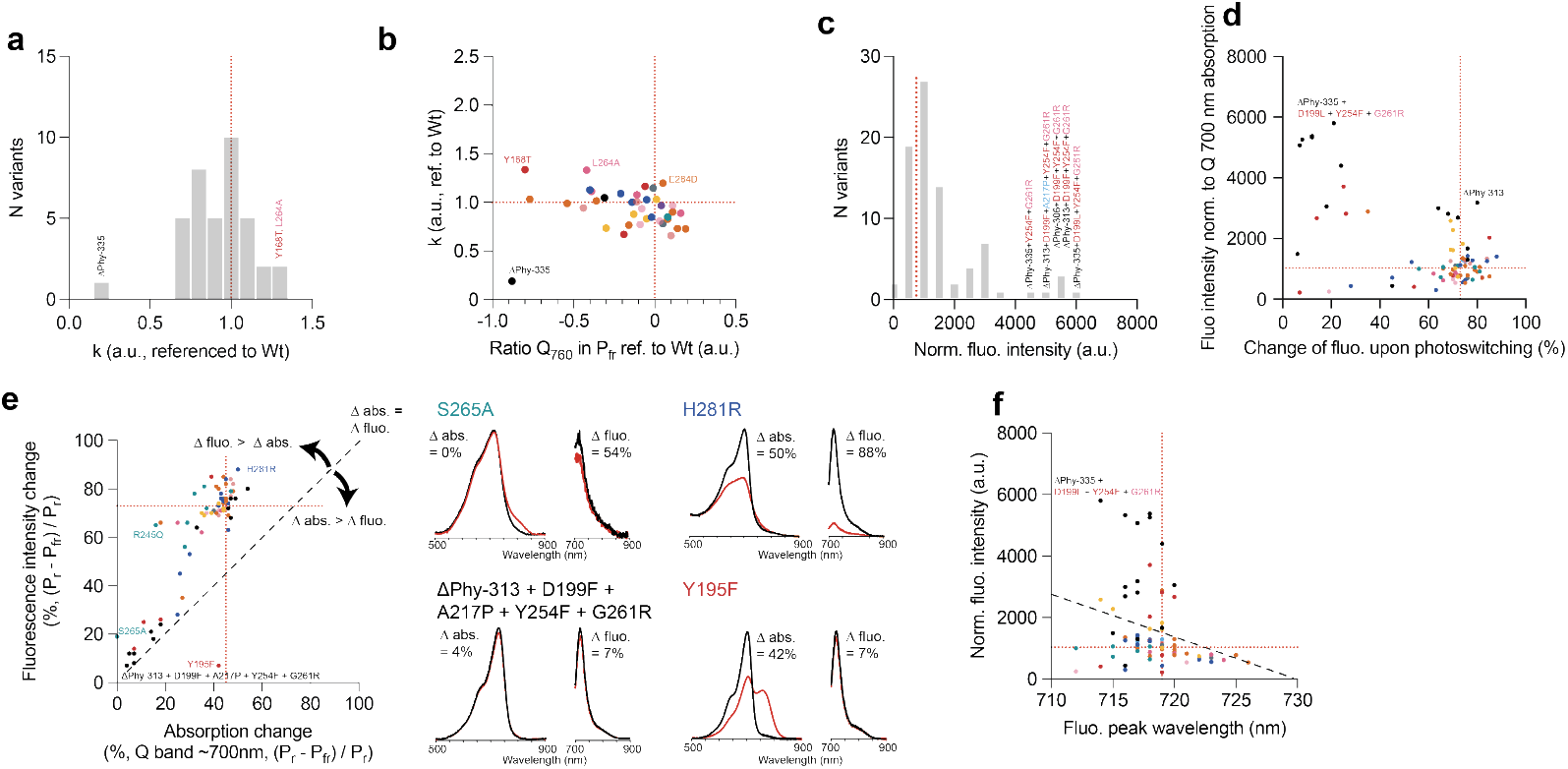
**a** Clustering of photoswitching kinetics for the P_fr_ → P_r_ transition with respect to ReBphP-PCM (=1). Variants that show the most extreme change are indicated colored as in 2a. The time constant is normalized to ReBphP-PCM. **b** Relation of photoswitching kinetic and absorption of the band driving the transition (P_fr_ state 760 nm band). Red dashed lines indicate ReBphP-PCM levels that have been used for normalization. Variants are color coded as in 2a with selected variants annotated. **c** Fluorescence intensity normalized to absorption of the 700 nm Q-band in the P_r_ state (excitation 680 nm). **d** Relation of fluorescence intensity and remaining photoswitchability. ReBphP-PCM fluorescence intensity indicated by red dashed line. **e** Relation of change of absorption of the 700 nm band upon photoswitching vs. change of fluorescence upon photoswitching. The dashed line indicates a 1:1 relation, i.e. the change of fluorescence is fully driven by the change in absorption of the 700 nm band. For selected variants the spectra are shown in the P_r_ (black) and P_fr_ (red) state for absorption and fluorescence (right) **f** Relation of fluorescence intensity and peak wavelength. Linear regression to illustrate dependence as black dashed line.

### Fluorescence of ReBphP-PCM variants

Like all known BphPs the fluorescence of ReBphP after excitation in the 700-band is comparably low (QY_fl_ 0.016). The same is true for most of the generated variants (**Figure 3c**). However, as described for several other BphPs the fluorescence can be drastically increased by destabilizing the *trans* form of the BV chromophore and favoring radiative deexcitation in the *cis* form^31^. In line with prior work^25^, we found this for positions directly interacting with the D-ring of the BV *trans*-isomer like D199 or Y254. However, the highest increase in fluorescence was achieved when also truncating the PHY domain (Δ_PHY_) and with this the PHY tongue. To some extent mutations to key PHY-tongue positions have similar increasing effects on fluorescence (R462 and F465). Beyond truncations, also a range of single positions like Y168T, V178L or L193H increase fluorescence. For our most fluorescent variant Δ_PHY_313+D199F+Y254F+G261R we studied the structural changes likely leading to the increased fluorescence compared to ReBphP-PCM. Similar to other fluorescent BphP variants^31^, an essential aspect is the loss of the ability to form a P_fr_ state due to the lost PHY tongue interactions as well as D199 and Y254 stabilizing the *trans* chromophore. Next to this, we find the *cis* BV of Δ_PHY_313+D199F+Y254F+G261R more stabilized than in ReBphP due to the contribution of several residues packing more tightly (Y195, P201, I250) as well as the mutation D199F resulting in an interaction of the PHE with the D-ring methyl group. Further, H251 makes a cleaner π-π interaction with the BV C-ring in the variant as compared to ReBphP-PCM. Overall, this results in a tighter chromophore pocket with a ~9 % smaller volume (**Suppl. Figure 9**). Along those lines the planarity of the BV in Δ_PHY_313+D199F+Y254F+G261R is increased compared to ReBphP-PCM with an overall planarity index increased by ~8 % (**Suppl. Figure 10**). Lastly, the ESPT route^5,6^ for Δ_PHY_313+D199F+Y254F+G261R via the “pyrrole water” seems less favorable when compared to ReBphP-PCM with the B-Ring pyrrole NH less optimally positioned (**Suppl. Figure 11**). All these aspects together likely contribute to the promotion of radiative deexcitation for Δ_PHY_313+D199F+Y254F+G261R^4,6,38^.

Despite the general trend clearly connecting high fluorescence with lack of photoswitching, several variants with relatively high fluorescence preserve photoswitching >60% (**Figure 3d**). For almost all variants the change in fluorescence upon photoswitching does not only stem from a change in absorptivity of the 700 nm band (**Figure 3e**, exemplary spectra). With some variants (e.g. H281R) exhibiting a change of fluorescence by far surpassing the change in absorption. This is likely due to heterogeneity of P_r_, where interconnected populations of P_r_ species show photoswitching or fluorescence^5^. Photoswitching variant Δ_PHY_313 shows comparably high fluorescence in *cis* despite still photoswitching. Some of the packing interactions, reduced EPST and heightened planarity described above can also be found for this variant, albeit not to a similar degree, explaining the lower fluorescence when compared to Δ_PHY_313+D199F+Y254F+G261R (**Suppl Figure 9-11**). The *trans* state in Δ_PHY_313 is less stabilized than in wildtype with H251 interaction changed and water molecules, as well as c-propionic and trans-acting amino acids forming a different interaction network (**Suppl. Figure 2**). This, and the different placing of the C-ring propionate described above might lead to the highly heterogenous nature of the *trans* state in Δ_PHY_313, possibly generally splitting into interconnected fluorescent and photoswitching populations as was suggested for recently discovered NIR photoswitching protein Penelope^39^. Lastly, high fluorescence is also associated with a blue shift of absorption and fluorescence spectra (**Figure 3f**), with the most fluorescent variant showing a peak at 715 nm or lower (Δ_PHY_335 + D199L + Y254F + G261R) and one of the least fluorescing a peak at 726 nm (K273E). This is in line with the energy gap law and observed by the general trend of engineered BphP being more fluorescent the further blue shifted the emission (**Suppl. Figure 12**).

### ReBphP variants as bright NIR fluorescence label

BphP optimized for fluorescence have extended the range of available fluorescent proteins considerably to the NIR^31,40,41^. Our variant Δ_PHY_313+D199F+Y254F+G261R shows an emission peak at 716 nm with a fluorescent quantum yield of 0.076 after 680 nm excitation. The variant is ~20% times brighter in solution and ~10% brighter when expressed in mammalian cell (**Figure 4a**) compared to the so far most red-shifted fluorescent protein iRFP720 (QY_fluo_ 0.06). The monomeric protein (**Suppl. Figure 13**) shows good tagging characteristics as demonstrated in a vimentin fusion expressed in Hela cells (**Figure 4b**). Photofatigue is ca. 5% after 1000 s of 12.3 mW/cm^2^ compared to 2% for iRFP720 (**Suppl. Figure 14**). Variant Δ_PHY_313+D199F+Y254F+G261R shows still minimal photoswitching, associated with a locked equilibrium state, i.e. a photophysical state that is inaccessible by photoswitching and can only be (re-) populated thermally a situation known from photoswitching GFP-like proteins like Dronpa^42^. Bright NIR proteins like SNIFP^11^ have been used for STED super-resolution microscopy. We determined the fluorescence lifetime to be 0.75 ± 0.02 ns which is in the same range as e.g. iRFP720^43^ (**Suppl. Figure 15**). Despite being slightly longer than SNIFP with 0.63 ns the lifetime is shorter than optimal for standard STED conditions (>1 ns)^44^. For example, the authors of SNIFP^11^ demonstrated a dedicated STED setup compensating for the shorter lifetimes with a shortened delay between excitation and STED pulse, in such a configuration Δ_PHY_313+D199F+Y254F+G261R could possibly show slightly improved results.

**Figure 4.**
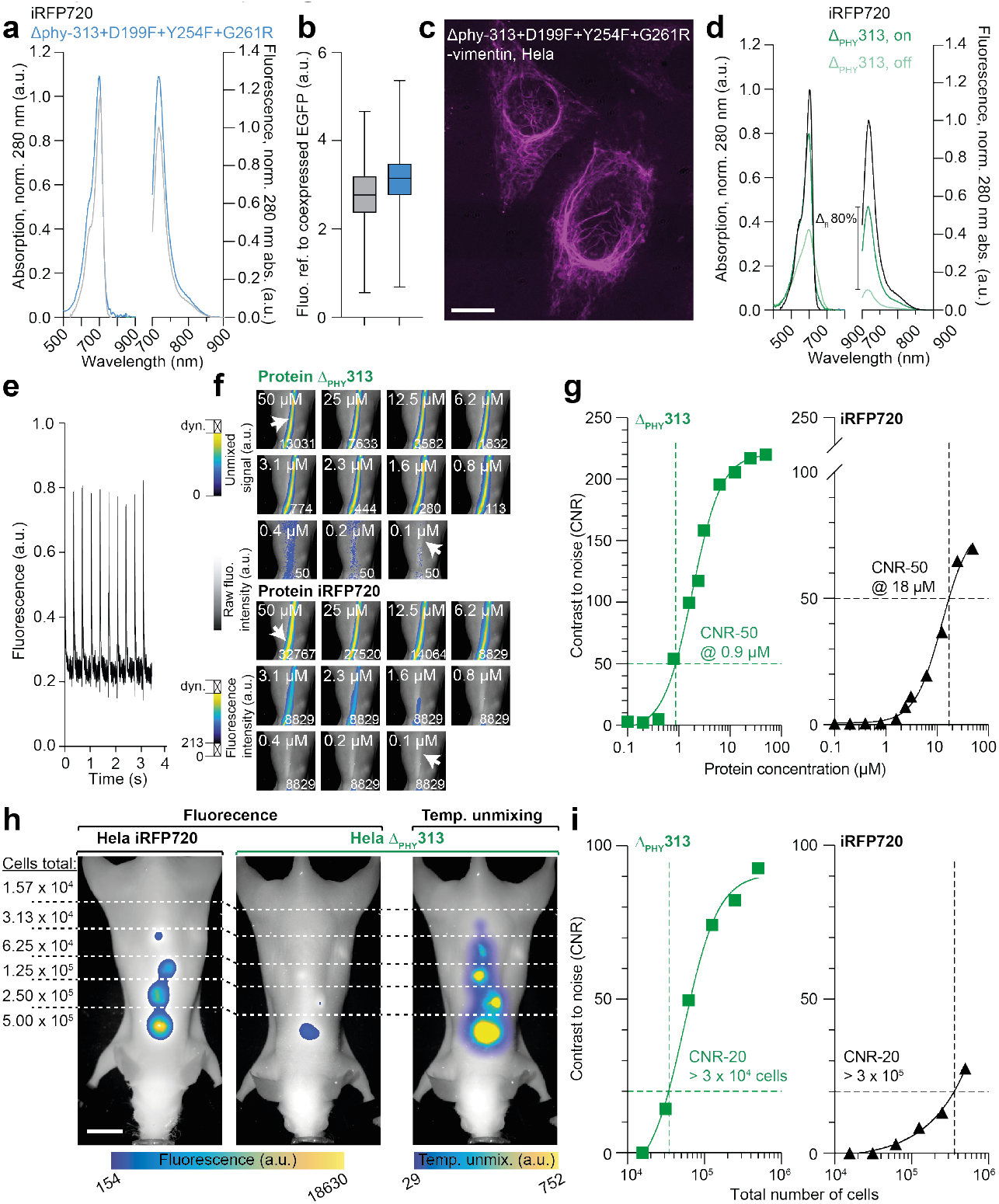
Characteristics and proof of concept applications for variants Δ_PHY_313+D199F+Y254F+G261R (**a** - **c**) and Δ_PHY_313 (**d – i**). Variant Δ_PHY_313+D199F+Y254F+G261R shows a ~20% higher fluorescence in solution compared to reference protein iRFP720 (**a**), this higher brightness translates into mammalian cells with ~10% higher brightness (**b**). Shown is the data for 300 individually segmented cells coexpressing an EGFP protein (G4S linker). **c** Δ_PHY_313+D199F+Y254F+G261R linked with vimentin expressed in Hela cells showing good tagging behavior. **d and e** The fluorescence of variant Δ_PHY_313 can be photoswitching by ~80% while the variant still retains a brightness of ca. 50% of <u>non-photoswitching</u> iRFP720. This photoswitching can be used to delineate the signal of Δ_PHY_313 from autofluorescence background. **f** Images of an *ex vivo* mouse model with a subcutaneous implant filled with varying concentrations of purified protein Δ_PHY_313 and iRFP720 (arrows in first and last image for orientation). Shown is the raw fluorescence in gray with a false color overlay of temporal unmixing for Δ_PHY_313 and fluorescence intensity for iRFP720. The false colors are scaled to a global minimum for each label but a dynamic maximum to prevent overglow (indicated in the bottom right corner of each image). **g** Contrast-to-noise ratios (CNR) for the two experiments. For comparison, the protein concentration corresponding to a CNR of 50 is indicated for both experiments. **h** *In vivo* mice with implants of different numbers of Hela cells expressing either Δ_PHY_313 or iRFP720. Shown are both mice fluorescence signal and the temporal unmixing of the mouse carrying the Δ_PHY_313 implants. Thresholding of false color to a level that leaves no background. **i** CNR rations of the *in vivo* experiments as described above.

### Fluorescent photoswitching ReBphP variants for whole animal NIR imaging

We identified several variants which show a considerable fluorescence while still retaining photoswitching capabilities. The brightest strongly photoswitching variant Δ_PHY_313 shows 80% of photoswitching while exhibiting a fluorescence intensity of ~54% (QY_fluo_ 0.038) in solution and ~20% in mammalian cells in comparison to iRFP720^8^. Interestingly, this combination of fluorescence and photoswitching is unique to this truncation position with truncations at 290 or 335 amino acids showing almost similar switching but less brightness (**Suppl. Table 1**). Δ_PHY_313 shows fast photoswitching between fluorescent and non-fluorescent state (P_r_ → P_fr_: t_1/2_~2 ms @80 mW/cm^2^), albeit ~5x slower than parent ReBphP-PCM (**Figure 4d**). In the tested energy range the switching accelerated exponentially saturating at ~2 ms. (**Suppl. Figure 16**). The fluorescence lifetime of Δ_PHY_313 is 0.52 ± 0.02 ns (**Suppl. Figure 15**) which is in-between the fluorescent non-switching variant Δ_PHY_313+D199F+Y254F+G261R (0.75 ± 0.02 ns) and the parent non-fluorescent and natively switching ReBphP-PCM (0.39 ± 0.02 ns). This aligns with the observation that engineering for a higher fluorescent quantum yield and reducing the switching goes along with a prolongation of the fluorescence lifetime^8^.

Next, we explored if Δ_PHY_313 can be used for lock-in augmented unmixing in NIR-whole animal imaging. Low concentrations of cells or agents are challenging to unmix for the tissue autofluorescence. Here the light-dependent fluorescence modulation of Δ_PHY_313 can help with unmixing. First, we compared a range of concentrations of purified Δ_PHY_313 and iRFP720 in a mouse *ex vivo* (**Figure 4e**). We chose iRFP720 as the furthest shifted NIR protein hence showing absorption and emission well in the NIR-I window. The mice were illuminated with repetitive patterns of 680 and 760 nm laser light (off/aq.: 110 ms x 65 and on: 6920 ms) and the fluorescence was detected with a camera behind a 700 or 800 nm longpass filter. For iRFP720 implants the data was considered and contrast-to-noise (CNR) determined solely on the fluorescence emission of the averaged images. For Δ_PHY_313 we employed a similar strategy as we use in OA photoswitching unmixing combining the differential of on/off signal, the exponential decay of the fluorescence and a frequency analysis of the known modulation^22,45^. Δ_PHY_313 shows a 20x higher CNR than iRFP720 based on its characteristics photoswitching fluorescence modulation. This allows excellent detection down to < 1 µM of protein. Next, we implanted similar cells in a mouse *in vivo* (**Figure 4f**). Also, here the photoswitching unmixing allows one order of magnitude better CNR compared iRFP720 despite its per cell signal in mammalian cells being 60 % lower than iRFP720 (brightness and expression). This effect is also preserved when looking only at the tail of the fluorescence using a longpass filter with an 800 nm cut off (**Suppl. Figure 17**). This suggests that such a temporal unmixing advantage might also be usable for SWIR infrared imaging.

Very recently, with Penelope^39^ another photoswitching NIR fluorescent BphP was presented. The proteins descents from SNIFP^11^ and hence DrBphP. It shows good brightness and photoswitching capabilities with a very fast dark relaxation in the milliseconds range making it usable for single wavelength RESOLFT. Our Δ_PHY_313 shows a dark relaxation of hours in solution, more comparable to ReBphP-PCM wildtype. This suggests that the intermediates potentially involved in the photoswitching process, although showing similar spectral characteristics (Cluster-P_fr_ 3), might be of different nature.

## Summary

We show a large-scale directed mutagenesis study on the so far only poorly researched BphP ReBphP from *Rhizobium etli*. Overall, the findings match with related studies on DrBphP^25^ or RpBphP^35^. Strikingly is the relative robustness of general chromophore attachment (5/70 single mutation variants chromophorylated) and of the general absorption characteristics of the P_r_ state (most variants ± 5 nm). This is true even for drastic changes in the pocket with mutations like R245A or H281D. However, many of the variants show effects on the photoswitching behavior. This includes the extent of P_fr_ state formed as well as the proportionality of the chromic effects of switching reaching from corresponding amplitudes of change to only changes in the 700-band or only changes in the 760-band. One of the reasons is the existence of substates for P_r_ and P ^26,46–49^, with different populations among variants. Most clearly this shows in a set of variants that, upon illumination in the 700-band, do not show the 760-band typically associated with P_fr_, but just a broadening of the 700-band. Likely this is associated with a switched non-P_fr_ state, possibly MetaR_c_^36^. Our structural analysis of the *trans* chromophores of ReBphP-PCM and Δ_PHY_313 variant showing this effect reveals a different interaction network around the chromophore mainly caused by a repositioned propionate side chain of the chromophore and associated waters molecules. Those different states are likely also associated with fluorescent variants of BphPs. The well-known approach of engineering fluorescent BphP by preventing photoswitching and stabilizing the *cis* isomer in a fluorescent compatible substate^31^ is also possible for ReBphP. Our variant Δ_PHY_313+D199F+Y254F+G261R brightness exceeding the well-known iRFP720. Next to preventing deexcitation through photoswitching our structural analysis reveals the fluorescence to stem from a more tightly packed pocket, higher chromophore planarity and closed ESPT pathways^4–6^. Next to this, we found a fluorescent variant that is still photoswitching. Apart from the very recently published photoswitching fluorescent BphP Penelope based on DrBphP^39^, this is the first demonstration that fluorescence and photoswitching can go together in BphP – opening the world of NIR reversibly switchable fluorescent proteins (rsNIRFPs). The substates of Δ_PHY_313 mentioned above also show in fluorescence excitation spectra indicating only a part of the Δ_PHY_313 population in a fluorescent compatible state – possibly distinct from the photoswitching population as suggested for Penelope^39^. It will be exciting to probe the impact on photoswitching with e.g. recording photoswitching action spectra. In contrast, the stability of the states between the two proteins are very different. Penelope shows very fast dark relaxation in milliseconds that allowed its use for single wavelength RESOLFT super resolution microscopy^39^. ReBphP based variant Δ_PHY_313 in contrast shows dark relaxations in the range of several hours despite equally fast photoswitching hinting a difference in the excited state vs. ground state isomerization processes. This is possibly due to different stabilizations of the photoswitched (illumination in the 700-band) state because only one canonical stabilizing residue in *trans* (D199) is preserved and Y254 mutated in Penelope^39^. Despite those differences the broadening of the 700-band described above is also found in Penelope^39^. It will be exciting to see, if such similarities and differences hint at general principles for rsNIRFPs. The slow switching makes Δ_PHY_313 usable for whole animal (or large organoid) locked-in unmixing from autofluorescence background. We show that Δ_PHY_313 can be used for temporal unmixing of labeled cells from background in whole animal imaging, increasing sensitivity. The increase in sensitivity can otherwise be only achieved with more complex methods like for example *in vivo* lifetime imaging^50^. Since the CNR enhancements also shows when detecting only the tail of the emission (> 800 nm) it can be presumed that the advantage propagates for SWIR imaging and might further boost its sensitivity^51^. The method bears resemblance to unmixing in microscopy using the optical lock in detection^52^ like eponymous OLID imaging^53^, SAFIRE^54^ or OPIOM^55^. Next to microscopy, surprisingly this has been demonstrated so far only in organisms like Xenopus larvae or Zebrafish^53^ despite available approaches with for example NIR absorbing Cy5 dye^56^. Despite the presented Δ_PHY_313 not being an optimized variant, it already shows 20x increase in CNR over bright iRFP720. This suggests that such proteins when improved can achieve even higher increase in CNR and hence sensitivity for detection of small cell populations in whole animal tissue or organoids even surpassing sensitivity advanced of e.g. lifetime imaging^50^ towards bioluminescence but with greater freedom of using fluorescent proteins.

## Material and Methods

### Cloning and Mutagenesis of constructs

Coding sequences of ReBphP-PCM, were PCR‐amplified as NdeI/XhoI fragments and inserted into the second multiple cloning site (MCS) of pET‐Duet1 (Novagen, Merck Millipore). For biliverdin synthesis, the heme oxygenase (HO) gene from *Nostoc sp*. was cloned into the first MCS of pET‐Duet1 using NcoI/HindIII restriction sites. Site-specific mutations were introduced to generate desired amino acid substitutions variants using Gibson Assembly (New England Biolabs). Mutation-containing primers were designed with overlapping regions (20–25 bp) complementary to the vector sequence flanking the insertion site. The target plasmid pET‐Duet1 backbone was digested with NdeI and XhoI to open the plasmid. The linearized PCR product was purified via gel extraction, and Gibson Assembly Master Mix was used to perform seamless ligation at 50 °C for 60 min, joining the overlapping ends to regenerate a circular plasmid carrying the desired mutation.

Mammalian expression constructs were based on an *AgeI* and *NotI* linearized pN1 vector (Addgene plasmid #45461; Vladislav Verkhusha). The coding sequences of BphP fragments were PCR-amplified from the corresponding BphP expression plasmids, while the sfGFP fragment was amplified from pET21a(+)-sfGFP. BphPs and sfGFP sequences were fused in frame through a flexible (G4S) linker using Gibson Assembly (New England Biolabs) to enable equimolar co-expression and fluorescence normalization in mammalian cells.

For cytoskeletal targeting experiments, The BphPs fragments were fused in frame G4S linker with the vimentin fragment, amplified from the mCherry-Vimentin-C-18 plasmid (Addgene #55157) via Gibson Assembly into the same pN1 backbone between *AgeI* and *NotI* sites. Additionally, the SNIFP coding sequence was PCR-amplified from pAAV_CAG_SNIFP_P2A_GFP (Addgene #197157) using the same cloning strategy for comparative analysis of expression and fluorescence intensity.

### Protein purification

For protein production, proteins were expressed in *E. coli* BL21 (DE3) cells (New England Biolabs, #C2527). With the secondary culture in TB medium supplemented with ampicillin (100 µg mL^−1^) grown at 37 °C until the optical density at 600 nm (OD_600_) reached 1.5-1.8. Protein expression was induced with IPTG (isopropyl-β-D-thiogalactopyranoside) at a final concentration of 0.5 mM and further growth for 16–18 h at 22 °C. The protein was purified using HiTrap chelating HP column (GE Healthcare, Illinois, USA) and following size-exclusion chromatography on a HiLoad 16/600 superdex 200pg column (GE Healthcare Life Sciences, Freiburg, Germany). The basic buffer in all steps being 50 mM Tris–HCl, 300 mM NaCl, pH8.

### Standard measurement of absorption, fluorescence and excitation spectra

All measurements, if not stated otherwise were performed immediately after purification of the protein. Absorption measurements were recorded using a Shimadzu UV-1800 spectrophotometer (Shimadzu Inc., Kyoto, Japan) or Cary 60 UV-Vis spectrophotometer (Agilent, Santa Clara, USA) with a 1 cm quartz cuvette. Fluorescence emission and excitation spectra were recorded on a Cary Eclipse fluorescence spectrophotometer (Varian Inc., Australia). The fluorescence emission spectra were recorded from 700-900 nm at an excitation wavelength of 680 nm using 10 nm excitation and emission slits. For the measurements the absorbance at 700 nm was adjusted to 0.1 to minimize inner filter effects. Photoswitching was performed by illuminating the sample in the cuvette from the top with a 660/20 nm (P_r_ → P_fr_) or 780/30 nm (P_fr_ → P_r_) light-emitting diode (Thorlabs) through a liquid lightguide (Thorlabs). The samples were illuminated and spectra measured immediately thereafter. This was repeated until no spectral change was visible, assuming a fully switched state.

### Time resolved spectral measurements and switching kinetics

For selected variants the measurements of fully switched states as described above were were complemented by a dynamic measurement of the switching process. Switching kinetics based on absorption spectra were recorded using an Ocean Optics HR-series high-resolution spectrometer coupled to a DH-2000-BAL broadband light source (Ocean Optics, USA). The photoswitching was achieved using 680 nm (P_r_ → P_fr_) and 750 nm (P_fr_ → P_r_) using laser sources (B&W TEK Inc). Laser intensity at the cuvette as stated in the figure legends. Rate constants of switching were determined by extracting the spectral development at a given wavelength and fitting with a single exponential.

### Equilibrium and thermal dark relaxation measurements

For dark relaxation measurements the purification described above was caried out in the dark or at 515 nm light. The dark relaxation was carried out at 22°C and followed based on absorption spectra recorded using a Cary 60 UV-Vis spectrophotometer. As a first step, the equilibrium was measured without any illumination. Thereafter, the protein was switched (see above) full to the state most distant from the equilibrium and relaxation followed by measuring absorption spectra in 30 min intervals for a time which allows to fit the relaxation with sufficient accuracy. At the end of the measurements, the proteins were photoswitched again to assess functional integrity and the P_r_ / P_fr_ absorption reachable. This final switching allows to assess how much functional protein was lost during the measurements which often took > 12 h.

### Measurement of fluorescent quantum yield

Fluorescence quantum yield (QY_fluo_) was determined in reference to iRFP720 (QY_fluo_ 0.06^8^). Fluorescence emission was measured for purified iRFP720, ReBphP-PCM, ReBphP-ΔPhy+D199L+R254F+G261R and ReBphP-ΔPhy at five concentrations with absorptions between 0.04 and 0.01. All measurements showed linearity. QY_fluo_ were calculated in respect to iRFP720 considering absorption emission and refractive index of the buffer^57^.

### Lifetime determination

The fluorescent lifetime (τ) was determined using a FS5 spectrofluorometer (Edinburgh Instruments), an SC-20 for liquid samples, and a time-correlated single photon-counting (TCPSC) module with an EPL 375 pulsed diode laser excitation source of 375 nm. τ was evaluated with Origin 2024 (OriginLab Corporation, Northampton, MA, USA) using the ExpDec1 function 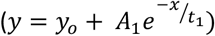.

### Mammalian cell culture

HeLa cells were cultured in Dulbecco’s Modified Eagle’s Medium (DMEM). The medium was supplemented with 10% fetal bovine serum (FBS; Invitrogen) and antibiotics (penicillin (100 U mL^−1^) and streptomycin (100 µg mL^−1^)). Cells were maintained at 37 °C in a humidified incubator with 5% CO_2_. Cultures were split every 2–3 days to sustain exponential growth and prevent overconfluence. Cells were transfected using Lipofectamine 2000 (Thermo Fisher Scientific) according to the manufacturer’s instructions. After transfection cells were cultured in antibiotic-free Opti-MEM reduced-serum medium. The medium was replaced with fresh DMEM after 4 h to minimize cytotoxicity and apply section pressure. Cells were further incubated for 48 h at 37 °C and 5% CO_2_ before imaging or further analyses.

### Microscopy

Imaging was performed using a Leica epi-fluorescence microscope (Leica DMI3000B) equipped with standard GFP and Cy5.5 filter sets. All images were acquired using identical exposure times and gain settings across samples to ensure comparability. Fluorescence of BphPs was normalized by dividing the mean Cy5.5 fluorescence intensity for each segmented cell with its corresponding GFP intensity to account for differences in expression level. Image analysis was carried out using FIJI/ImageJ. Vimentin-tagged constructs were imaged on a Zeiss Axio Imager.M2 upright fluorescence microscope using identical filter configurations and objectives for consistent comparison.

### Whole animal imaging system

Whole animal imaging was conducted using a custom-built non-magnified, objective-free whole-animal optical imaging system. The sample plane was illuminated using three continuous-wave laser diodes at 488 nm (Thorlabs), 671 nm or 786 nm (Lumics, Berlin). The latter two allowed photoswitching and excitation of fluorescence while the 488 nm light was used to excite coexpressed EGFP for normalization purposes. Light was homogenized using an engineered diffuser and shuttered with a 10 ms response time shutter to prevent ramping times of the laser interfering with the photoswitching kinetics. Fluorescence emission was recorded using a PCO Edge scientific CMOS monochrome camera, equipped with appropriate 500nm, 700nm and 800 long-pass filters to isolate fluorescence signals from excitation light.

### Whole animal Imaging

All animal experiments were conducted in compliance with institutional and governmental regulations and were approved by the Government of Upper Bavaria, Germany. For the ***ex vivo* tubing experiments**, *Re*Bphp-CBD-D_PHY_313 and *iRFP720* proteins were prepared in Tris-HCl buffer at concentrations ranging from 50 µM to 0.0488 µM. Protein samples were loaded into transparent polymer tubing (inner diameter = 0.58 mm; RCT Reichelt Chemietechnic GmbH), which was subcutaneously positioned in sacrificed FoxN1 nude mice. Each tube was filled sequentially with a single protein sample at a defined concentration, followed by imaging and buffer washing steps between measurements. For ***in vivo* imaging**, HeLa cells expressing *Re*Bphp-CBD-D_PHY_313 and *iRFP720* were prepared at various densities ranging from 500 × 10^3^ to 15.6 × 10^3^ cells. Cells were mixed with Matrigel (Corning) in equal parts (1:1, v/v) to improve cell survival and keep them in place after injection. Cell suspensions were implanted subcutaneously in the back of two FoxN1 nude mice (Charles River Laboratories, Boston, USA). Each mouse received cells expressing only one type of fluorescent protein. To minimize background body autofluorescence, all mice were transitioned to a western diet two weeks before imaging. All live animal imaging was performed using the custom imaging platform described above. Mice were anesthetized with 2% isoflurane delivered in medical oxygen and positioned securely within the imaging chamber. Fluorescence signals were recorded under identical imaging parameters for direct comparison of both constructs.

### Data analysis and statistical methods

The mouse imaging time-series were analysed in MATLAB 2024b. For *in vivo* imaging data, a motion correction step was applied before the image analysis to minimize the impact of respiratory and cardiac motion. The motion correction algorithm uses a non-rigid approach^58^, estimating a 2D displacement field for each frame in the image time-series. To reduce the magnitude of the displacements and lower the amount of required interpolation, the reference frame was selected algorithmically according to its similarity to the other frames in the time-series based on the highest mean structural similarity index (SSIM).

The *in vivo* and *ex vivo* imaging time-series were processed in an identical manner. To reduce noise, each frame was smoothed with a square kernel of 11 × 11 pixels (box filter) and down-sampled to half-size. For iRFP720, all frames in the time-series were averaged, resulting in a single image which was subsequently used for the fluorescence signal analysis. In case of Δ_PHY_313, an approach based on well-established temporal unmixing algorithm used in OA imaging^22,45,59^ was used to separate the photoswitching signal from the background. In brief: the image time-series recorded for Δ_PHY_313 captures cycles of P_r_ → P_fr_ transitions. The temporal unmixing algorithm separates the modulation of the photoswitching signal from the non-modulating background by analysing three aspects of the time-series: The differential of the on/off signal (switching delta), exponential decay of the signal (switching kinetics), and frequency of the signal modulation. These aspects were analysed per-pixel with the frequency analysis considering the full time-series, while the differential and signal decay are computed on a mean cycle obtained by averaging all recorded cycles of P_fr_ → P_r_ transitions. More specifically, the differential of the on/off signal can be expressed in the form *d* = *f*_0*N*_ − *f*_0*FF*_, where *f*_0*N*_ and *f*_0*FF*_ are pixel values representing the on and off signal, respectively. The *f*_0*N*_ and *f*_0*FF*_ were obtained by averaging the first two and last two values in a mean cycle to reduce noise. The kinetic of the switching signal was estimated by fitting a negative exponential of the form *f*_*t*_ = *a* · *e*^−*kt*^, where *f*_*t*_ is the value in the mean cycle at time *t, a* is the initial value, and *k* is the decay rate. A coefficient of determination *R*^2^ was computed to measure how well the negative exponential fits the signal. Next, a discrete Fourier transform of the full recorded time-series was computed to determine the coefficient of the modulation frequency *F*_*mod*_ (where the frequency is known a priori from the illumination and recording schedule) and sum of the coefficients of its harmonic frequencies *F*_*harm*_. Finally, temporal unmixing was computed as a weighted sum *u* = ∑ *w*_1_ · *d* + *w*_2_ · *k* + *w*_3_ · *R*^2^ + *w*_4_ · *F*_*mod*_ + *w*_5_ · *F*_*harm*_, where *w*_1_ to *w*_5_ are weights of the computed signal features. This step concluded the signal processing and was subsequently followed by the evaluation.

To evaluate the contrast over background of iRFP720 and Δ_PHY_313 labels, we computed contrast-to-noise ratios given by *CNR* = (*S*_*label*_ − *S*_*background*_) / *σ*_*background*_. Here, *S*_*label*_ is the signal in the region of interest (ROI) representing the label, and *S*_*background*_ and *σ*_*background*_ are the signal and standard deviation in the ROI representing the background, respectively.

### Computational methods

In short, structures were generated through a chimeric AI-structure prediction approach (combining Chai-1^60^and Boltz-2^61^ predictions) and further structural refinement of the chromophore active site via hybrid QM/MM geometry optimizations (PBE-D4/def2-SVP:AMBER) conducted in ORCA 6.0^62^. To ensure biological accuracy, homology information from high-resolution templates was integrated: the BV binding pocket was modeled based on crystal structures (PDB-IDs: 6G1Y^63^ and 5LLW^63^), which also served as templates for the characteristic secondary structure of the PHY-tongue. This workflow ensures that the models represent well-defined energetic minima on the potential energy surface.

To solvate the structural models internal water molecule positions and the ones of the first and second solvation sphere were predicted using an adaptation of the Vedani algorithm^64^ implemented in MAXIMOBY (CHEOPS, Germany, version 2026). The position of the identified two key water molecules within the active site are consistent with established mechanisms. The models incorporate the highly conserved ‘pyrrole water’^63^ and— specifically for the *cis*-isomer—an additional water molecule in the QM region. The latter is positioned as an ESPT-mediator between the D-ring and the propionate side chain, following the structural motif suggested by Grigorenko et al. for IFP1.4^65^. While these configurations represent stationary states rather than dynamic ensembles, they provide a robust and high-resolution framework for discussing the hydrogen-bonding networks and the structural basis of the biliverdin photoisomerization. (Detailed computational methods see **Suppl. Text 1**).

## Supporting information

Suppl. Mat.

## Author contributions

AM cloned the constructs, measured data, performed mammalian imaging and contributed to the analysis; DW analyzed the imaging data; AD contributed to the spectroscopy measurements protein purification and analysis; EW performed and analyzed computational structure modeling; SG set up whole animal imaging experimentation; SW performed and analyzed lifetime data; UK performed animal experimentation; SeS contributed to cloning and protein purification; BP contributed to the photoswitching imaging; JG, IR and ADu contributed to lifetime data; ER helped purification and contributed to the manuscript; VN contributed to the manuscript; TR supervised computational structure modeling and contributed to its analysis; ACS conceived study, contributed to the analysis and wrote manuscript with contributions from all authors.

## Acknowledgments

The authors thank Stefan Schlenz and Udo Höweler for assisting with the computational structure optimization, Yona Perstat for help with the analysis of the mouse imaging data and Javier Garcia Lopez for discussions on the early manuscript.

## Bibliography

1 Auldridge, M. E. & Forest, K. T. Bacterial phytochromes: more than meets the light. Crit Rev Biochem Mol Biol 46, 67–88 (2011). 10.3109/10409238.2010.546389

2 Shcherbakova, D. M., Shemetov, A. a., Kaberniuk, A.a. & Verkhusha, V. V. Natural Photoreceptors as a Source of Fluorescent Proteins, Biosensors, and Optogenetic Tools. Vol. 84 (2014).

3 Takala, H., Edlund, P., Ihalainen, J. A. & Westenhogff, S. Tips and turns of bacteriophytochrome photoactivation. Photochem Photobiol Sci 19, 1488–1510 (2020). 10.1039/d0pp00117a

4 Bhattacharya, S., Auldridge, M. E., Lehtivuori, H., Ihalainen, J. A. & Forest, K. T. Origins of fluorescence in evolved bacteriophytochromes. J Biol Chem 289, 32144–32152 (2014). 10.1074/jbc.M114.589739

5 Toh, K. C., Stojkovic, E. A., van Stokkum, I. H., Moffat, K. & Kennis, J. T. Proton-transfer and hydrogen-bond interactions determine fluorescence quantum yield and photochemical egiciency of bacteriophytochrome. Proc Natl Acad Sci U S A 107, 9170–9175 (2010). 10.1073/pnas.0911535107

6 Toh, K. C., Stojkovic, E. A., van Stokkum, I. H., Moffat, K. & Kennis, J. T. Fluorescence quantum yield and photochemistry of bacteriophytochrome constructs. Phys Chem Chem Phys 13, 11985–11997 (2011). 10.1039/c1cp00050k

7 Mathes, T. et al. Femto-to Microsecond Photodynamics of an Unusual Bacteriophytochrome. J Phys Chem Lett 6, 239–243 (2015). 10.1021/jz502408n

8 Shcherbakova, D. M. & Verkhusha, V. V. Near-infrared fluorescent proteins for multicolor in vivo imaging. Nat Methods 10, 751–754 (2013). 10.1038/nmeth.2521

9 Shcherbakova, D. M. et al. Bright monomeric near-infrared fluorescent proteins as tags and biosensors for multiscale imaging. Nat Commun 7, 12405 (2016). 10.1038/ncomms12405

10 Auldridge, M. E., Satyshur, K. A., Anstrom, D. M. & Forest, K. T. Structure-guided engineering enhances a phytochrome-based infrared fluorescent protein. J Biol Chem 287, 7000–7009 (2012). 10.1074/jbc.M111.295121

11 Kamper, M., Ta, H., Jensen, N. A., Hell, S. W. & Jakobs, S. Near-infrared STED nanoscopy with an engineered bacterial phytochrome. Nat Commun 9, 4762 (2018). 10.1038/s41467-018-07246-2

12 Filonov, G. S. et al. Bright and stable near-infrared fluorescent protein for in vivo imaging. Nat Biotechnol 29, 757–761 (2011). 10.1038/nbt.1918

13 Rogers, O. C., Johnson, D. M. & Firnberg, E. mRhubarb: Engineering of monomeric, red-shifted, and brighter variants of iRFP using structure-guided multi-site mutagenesis. Sci Rep 9, 15653 (2019). 10.1038/s41598-019-52123-7

14 Matlashov, M. E. et al. A set of monomeric near-infrared fluorescent proteins for multicolor imaging across scales. Nat Commun 11, 239 (2020). 10.1038/s41467-019-13897-6

15 Wang, L. V. & Hu, S. Photoacoustic tomography: in vivo imaging from organelles to organs. Science 335, 1458–1462 (2012). 10.1126/science.1216210

16 Ntziachristos, V. Going deeper than microscopy: the optical imaging frontier in biology. Nat Methods 7, 603–614 (2010). 10.1038/nmeth.1483

17 Stiel, A. C. & Ntziachristos, V. Controlling the sound of light: photoswitching optoacoustic imaging. Nat Methods 21, 1996–2007 (2024). 10.1038/s41592-024-02396-2

18 Li, L. et al. Small near-infrared photochromic protein for photoacoustic multicontrast imaging and detection of protein interactions in vivo. Nat Commun 9, 2734 (2018). 10.1038/s41467-018-05231-3

19 Yao, J. et al. Multiscale photoacoustic tomography using reversibly switchable bacterial phytochrome as a near-infrared photochromic probe. Nat Methods 13, 67–73 (2016). 10.1038/nmeth.3656

20 Märk, J. et al. Dual-wavelength 3D photoacoustic imaging of mammalian cells using a photoswitchable phytochrome reporter protein. Communications Physics 1, 3–3 (2018). 10.1038/s42005-017-0003-2

21 Chee, R. K. W., Li, Y., Zhang, W., Campbell, R. E. & Zemp, R. J. In vivo photoacoustic difference-spectra imaging of bacteria using photoswitchable chromoproteins. J Biomed Opt 23, 1–11 (2018). 10.1117/1.JBO.23.10.106006

22 Mishra, K. et al. Multiplexed whole-animal imaging with reversibly switchable optoacoustic proteins. Sci Adv 6, eaaz6293 (2020). 10.1126/sciadv.aaz6293

23 Shu, X. et al. Mammalian expression of infrared fluorescent proteins engineered from a bacterial phytochrome. Science 324, 804–807 (2009). 10.1126/science.1168683

24 Rottwinkel, G., Oberpichler, I. & Lamparter, T. Bathy phytochromes in rhizobial soil bacteria. J Bacteriol 192, 5124–5133 (2010). 10.1128/JB.00672-10

25 Wagner, J. R. et al. Mutational analysis of Deinococcus radiodurans bacteriophytochrome reveals key amino acids necessary for the photochromicity and proton exchange cycle of phytochromes. J Biol Chem 283, 12212–12226 (2008). 10.1074/jbc.M709355200

26 Chenchiliyan, M. et al. Ground-state heterogeneity and vibrational energy redistribution in bacterial phytochrome observed with femtosecond 2D IR spectroscopy. J Chem Phys 158, 085103 (2023). 10.1063/5.0135268

27 von Stetten, D. et al. Chromophore heterogeneity and photoconversion in phytochrome crystals and solution studied by resonance Raman spectroscopy. Angew Chem Int Ed Engl 47, 4753–4755 (2008). 10.1002/anie.200705716

28 Nieder, J. B., Brecht, M. & Bittl, R. Dynamic intracomplex heterogeneity of phytochrome. J Am Chem Soc 131, 69–71 (2009). 10.1021/ja8058292

29 Kirpich, J. S. et al. Protonation Heterogeneity Modulates the Ultrafast Photocycle Initiation Dynamics of Phytochrome Cph1. J Phys Chem Lett 9, 3454–3462 (2018). 10.1021/acs.jpclett.8b01133

30 Velazquez Escobar, F. et al. Protonation-Dependent Structural Heterogeneity in the Chromophore Binding Site of Cyanobacterial Phytochrome Cph1. J Phys Chem B 121, 47–57 (2017). 10.1021/acs.jpcb.6b09600

31 Shcherbakova, D. M., Baloban, M. & Verkhusha, V. V. Near-infrared fluorescent proteins engineered from bacterial phytochromes. Curr Opin Chem Biol 27, 52–63 (2015). 10.1016/j.cbpa.2015.06.005

32 Hahn, J. et al. Probing protein-chromophore interactions in Cph1 phytochrome by mutagenesis. FEBS J 273, 1415–1429 (2006). 10.1111/j.1742-4658.2006.05164.x

33 von Stetten, D. et al. Highly conserved residues Asp-197 and His-250 in Agp1 phytochrome control the proton affinity of the chromophore and Pfr formation. J Biol Chem 282, 2116–2123 (2007). 10.1074/jbc.M608878200

34 Stojkovic, E. A. et al. FTIR Spectroscopy Revealing Light-Dependent Refolding of the Conserved Tongue Region of Bacteriophytochrome. J Phys Chem Lett 5, 2512–2515 (2014). 10.1021/jz501189t

35 Yang, X., Stojkovic, E. A., Kuk, J. & Moffat, K. Crystal structure of the chromophore binding domain of an unusual bacteriophytochrome, RpBphP3, reveals residues that modulate photoconversion. Proc Natl Acad Sci U S A 104, 12571–12576 (2007). 10.1073/pnas.0701737104

36 Lehtivuori, H., Rumfeldt, J., Mustalahti, S., Kurkinen, S. & Takala, H. Conserved histidine and tyrosine determine spectral responses through the water network in Deinococcus radiodurans phytochrome. Photochem Photobiol Sci 21, 1975–1989 (2022). 10.1007/s43630-022-00272-6

37 Yi, C., Meier, S. S. M., Kehr, M. & Moglich, A. Spectroscopic Analyses of Photoconversion in Phytochromes. Methods Mol Biol 2970, 177–191 (2026). 10.1007/978-1-0716-4791-2_11

38 Samma, A. a., Johnson, C. K., Song, S., Alvarez, S. & Zimmer, M. On the origin of flurescence in bacteriophytochrome infrared fluorescent proteins. J. Phys. Chem. B 114, 15362–15369 (2010). 10.1021/jp107119q.On

39 Stumpf, D. et al. The near-infrared bacteriophytochrome-derived fluorescent protein PENELOPE enables RESOLFT superresolution microscopy. Proc Natl Acad Sci U S A 122, e2504748122 (2025). 10.1073/pnas.2504748122

40 Shcherbakova, D. M., Stepanenko, O. V., Turoverov, K. K. & Verkhusha, V. V. Near-Infrared Fluorescent Proteins: Multiplexing and Optogenetics across Scales. Trends Biotechnol 36, 1230–1243 (2018). 10.1016/j.tibtech.2018.06.011

41 Piatkevich, K. D. et al. Near-Infrared Fluorescent Proteins Engineered from Bacterial Phytochromes in Neuroimaging. Biophys J 113, 2299–2309 (2017). 10.1016/j.bpj.2017.09.007

42 Habuchi, S. et al. Photo-induced protonation/deprotonation in the GFP-like fluorescent protein Dronpa: mechanism responsible for the reversible photoswitching. Photochem Photobiol Sci 5, 567–576 (2006). 10.1039/b516339k

43 Zhu, J., Shcherbakova, D. M., Hontani, Y., Verkhusha, V. V. & Kennis, J. T. Ultrafast excited-state dynamics and fluorescence deactivation of near-infrared fluorescent proteins engineered from bacteriophytochromes. Sci Rep 5, 12840 (2015). 10.1038/srep12840

44 Jahr, W., Velicky, P. & Danzl, J. G. Strategies to maximize performance in STimulated Emission Depletion (STED) nanoscopy of biological specimens. Methods 174, 27–41 (2020). 10.1016/j.ymeth.2019.07.019

45 Stankevych, M., Mishra, K., Ntziachristos, V. & Stiel, A. C. in Methods in Enzymology Vol. 657 365–383 (Elsevier Inc., 2021).

46 Wang, C. et al. Bacteriophytochrome Photoisomerization Proceeds Homogeneously Despite Heterogeneity in Ground State. Biophys J 111, 2125–2134 (2016). 10.1016/j.bpj.2016.10.017

47 Velazquez Escobar, F. et al. Conformational heterogeneity of the Pfr chromophore in plant and cyanobacterial phytochromes. Front Mol Biosci 2, 37 (2015). 10.3389/fmolb.2015.00037

48 Song, C. et al. Two ground state isoforms and a chromophore D-ring photoflip triggering extensive intramolecular changes in a canonical phytochrome. Proc Natl Acad Sci U S A 108, 3842–3847 (2011). 10.1073/pnas.1013377108

49 Rao, A. G., Wiebeler, C., Sen, S., Cerutti, D. S. & Schapiro, I. Histidine protonation controls structural heterogeneity in the cyanobacteriochrome AnPixJg2. Phys Chem Chem Phys 23, 7359–7367 (2021). 10.1039/d0cp05314g

50 Rice, W. L., Shcherbakova, D. M., Verkhusha, V. V. & Kumar, A. T. In vivo tomographic imaging of deep-seated cancer using fluorescence lifetime contrast. Cancer Res 75, 1236–1243 (2015). 10.1158/0008-5472.CAN-14-3001

51 Wang, F., Zhong, Y., Bruns, O., Liang, Y. & Dai, H. In vivo NIR-II fluorescence imaging for biology and medicine. Nature Photonics 18, 535–547 (2024). 10.1038/s41566-024-01391-5

52 Yan, Y., Marriott, M. E., Petchprayoon, C. & Marriott, G. Optical switch probes and optical lock-in detection (OLID) imaging microscopy: high-contrast fluorescence imaging within living systems. Biochem J 433, 411–422 (2011). 10.1042/BJ20100992

53 Marriott, G. et al. Optical lock-in detection imaging microscopy for contrast-enhanced imaging in living cells. Proc Natl Acad Sci U S A 105, 17789–17794 (2008). 10.1073/pnas.0808882105

54 Richards, C. I. et al. Optically modulated fluorophores for selective fluorescence signal recovery. J Am Chem Soc 131, 4619–4621 (2009). 10.1021/ja809785s

55 Querard, J. et al. Resonant out-of-phase fluorescence microscopy and remote imaging overcome spectral limitations. Nat Commun 8, 969 (2017). 10.1038/s41467-017-00847-3

56 Fan, C., Hsiang, J. C. & Dickson, R. M. Optical modulation and selective recovery of Cy5 fluorescence. Chemphyschem 13, 1023–1029 (2012). 10.1002/cphc.201100671

57 Rurack, K. In Standardization and Quality Assurance in Fluorescence Measurements Vol. 1 (ed Ute Resch-Genger) (Springer, 2008).

58 Vishnevskiy, V., Gass, T., Szekely, G., Tanner, C. & Goksel, O. Isotropic Total Variation Regularization of Displacements in Parametric Image Registration. IEEE Trans Med Imaging 36, 385–395 (2017). 10.1109/TMI.2016.2610583

59 Huang, Y. et al. Photoswitching protein-XTEN fusions as injectable optoacoustic probes. Acta Biomater 195, 536–546 (2025). 10.1016/j.actbio.2025.02.002

60 Boitreaud, J. et al. (2024). 10.1101/2024.10.10.615955

61 Passaro, S. et al. Boltz-2: Towards Accurate and Efficient Binding Affinity Prediction. bioRxiv (2025). 10.1101/2025.06.14.659707

62 Neese, F. Software update: The ORCA program system—Version 5.0. WIREs Computational Molecular Science 12 (2022). 10.1002/wcms.1606

63 Schmidt, A. et al. Structural snapshot of a bacterial phytochrome in its functional intermediate state. Nat Commun 9, 4912 (2018). 10.1038/s41467-018-07392-7

64 Vedani, A. & Huhta, D. W. Algorithm for the systematic solvation of proteins based on the directionality of hydrogen bonds. Journal of the American Chemical Society 113, 5860–5862 (2002). 10.1021/ja00015a049

65 Grigorenko, B. L., Polyakov, I. V. & Nemukhin, A. V. Modeling photophysical properties of the bacteriophytochrome-based fluorescent protein IFP1.4. J Chem Phys 154, 065101 (2021). 10.1063/5.0026475

