## Supplementary material for "Mutagenesis study of a Bacteriophytochrome – insights for the development of labels for optical imaging": Suppl. Mat.

**Supplementary Text 1: Structure computational methods**

*Structure Prediction and Modeling*

Initial structural models of ReBPHP were generated using a hybrid procedure combining AI-based structure prediction with template-based homology modeling. Five $P_{fr}$-state models of the PCM wild-type were predicted using Chai-1[1] and Boltz-2[2], respectively. To ensure an accurate representation of the biliverdin (BV) binding pocket, the crystal structure of a homologous phytochrome (PDB-ID: 6G1Y[3]) served as structural template. Similarly, five $P_r$-state models were generated using both tools, utilizing the crystal structure with PDB-ID 5LLW[3] as template. In all cases, the BV chromophore was included as a covalently bound ligand during the initial prediction.

For the $P_{fr}$-state, Boltz-2 showed superior performance in predicting the structural features of the PAS-GAF domain, whereas Chai-1 was more accurate for the $P_r$-state PAS-GAF domain when compared to the homologous structures 6G1Y, 5LLW, and 6ETZ[3]. We focused on the residues in the immediate vicinity of the chromophore. Notably, the PHY-Domain required a chimeric approach: Boltz-1 correctly predicted the characteristic sheet secondary structure of the $P_r$-state, while Chai-1 successfully reproduced the helical fold typical of the $P_{fr}$-state PHY-tongue. Consequently, the structure prediction with the highest confidence and with the closest structural alignment with the homologous templates for the respective states were combined to obtain refined starting models for ReBPHP. Conserved side chains known to form the chromophore binding pocket were manually inspected and if needed manually adjusted to ensure consistency with well-established interaction mechanisms. Structural models for the truncated $\Delta_{PHY}$313 variant were derived by using the corresponding shortened sequence used in the experiments. Finally, the $\Delta_{PHY}$313+D199F+Y254F+G261R variant was generated using the PyMOL mutagenesis wizard, where sidechain rotamers were selected in such way to optimize local interactions and avoid steric clashes.

*System Preparation and Molecular Mechanics*

The six structural models comprising the $P_r$ and $P_{fr}$ states of each variant were protonated, solvated, and subjected to energy minimization using the MAXIMOBY software package (CHEOPS, Germany, version 2026). To ensure compatibility with the AMBER force field[4] implemented in MAXIMOBY, partial charges for the BV chromophore were determined via Restricted Hartree-Fock (RHF/STO-3G) calculations. Electrostatic Potential (ESP) fit charges were derived as implemented in Gaussian 09. Following established models for bacteriophytochrome-bound chromophores[5] the BV protonation state was set with all four pyrrole rings protonated and both propionate side chains deprotonated, resulting in a net charge of -1. Amino acid protonation states were assigned based on local $pK_a$ calculations according to the method of Nielsen and Vriend[6], as implemented in MAXIMOBY.

All six systems were then solvated by predicting the positioning of all internal water molecules as well as the water molecules of first and second solvation sphere: the first sphere was placed to satisfy all potential hydrogen-bonding sites of the protein and chromophore, while a second sphere was added to coordinate the primary hydration layer. The water molecule position prediction and placement was performed by the MAXIMOBY solvation algorithm that is based on the algorithm of Vedani and Huth[7]. This procedure included the prediction of the highly conserved "pyrrole water" (PW)[3], which is coordinated by the A, B, and C-rings of the chromophore. Additionally, for the *cis* conformation, an ESPT-facilitating water (EW) was positioned between the D-ring pyrrole and the C-propionate group as reported by Grigorenko et al. for IFP1.4[60]. Molecular mechanics (MM) optimizations for energetic relaxation and to resolve local clashes were performed using the BFGS algorithm and the AMBER force field within the MAXIMOBY environment.

*QM/MM calculations*

The as aforementioned prepared structures of ReBPHP-PCM, $D_{PHY}$313 and $D_{PHY}$313+D199F+Y254F+G261R in the respective $P_r$ and $P_{fr}$ state were optimized using the hybrid quantum mechanics/molecular mechanics (QM/MM) algorithm in ORCA6.0[8]. Within the ORCA environment, the QM region's electronic density was initialized with a Mulliken charge calculation. The QM region was defined to include the BV chromophore, the conserved 'pyrrole water' (PW), and—specifically for the *cis*-isomers—the 'ESPT-water' (EW). Additionally, the sidechain of Cys13 was included in the QM region and the QM/MM boundary was placed between the $C_\alpha$ and $C_\beta$ atom, ensuring that the cut does not truncate formal charge groups. A hydrogen link atom was employed to saturate the dangling bond of the QM region in each system, as implemented in the ORCA QM/MM module. To allow for sufficient structural relaxation, an active MM region was defined, encompassing all residues within a 5.0 Å radius of the chromophore. To maintain structural integrity and avoid artificial strain at the QM/MM boundary, only complete amino acid residues were included in the active set. All remaining protein and solvent atoms were kept frozen at their MM-optimized positions to maintain the overall scaffold integrity.

The QM region, comprising 85 (*trans*) and 88 (*cis*) atoms, was treated at the PBE/def2-SVP level of theory[9,10]. To accurately capture the non-bonded interactions within the binding pocket, the D4 dispersion correction was applied[11]. The Resolution of Identity (RI) approximation with AutoAux auxiliary basis sets was utilized to accelerate the electronic structure calculations[12]. Convergence was ensured by employing a TightSCF criterion ($10^{-8}$Eh). The geometry was considered optimized when the standard ORCA convergence criteria for gradients and displacements were met. The MM-Region was handled with the AMBER forcefield.

*Structural Analysis and Interaction Mapping*

To characterize the protein-chromophore interface, inter-domain interactions, and the coordination of the internal water network, non-covalent interactions were identified using PyContact[13] and the contact-matrix algorithm implemented in MAXIMOBY. For volume calculation of the BV binding pocket the PyMOL Plugin PyVOL [10.1101/816702] was used employing a minimum probe radius of 1.2 Å and a maximum probe radius of 3.4 Å. The planarity of the biliverdin chromophore was evaluated by measuring the dihedral angles of the methine bridges in PyMOL. Additionally, interplanar angles (Θ) were determined by calculating the normal vectors of each pyrrole ring's least-squares plane. To assess the overall distortion of the chromophore, the root-mean-square deviation (RMSD) from the least-squares plane of the core heavy atoms was calculated

**Supplementary Table 1:** Overview of variants generated in this study. Colors indicate residue or cluster groups or min/max.

| AA Position | Group | Mutation | Cluster $P_r$ | Cluster $P_{fr}$ | Cluster Diff. | Weighted Soret to 280 (a.u.) | Q peak in Pr (nm) | Q to Soret (a.u.) | Fluo. peak (nm) | Corr. fluo. signal (a.u.) | Brigthness (a.u.) | Δ fluo. (%) | Δ abs. ~700 nm (%) |
|---|---|---|---|---|---|---|---|---|---|---|---|---|---|
| ***ReBphP-PCM wildtype*** | | | | | | | | | | | | | |
| | | ReBphP-PCM (N = 11) | 1 | 1 | 1 | 94 ± 30 | 703 ± 0.5 | 2.38 ± 0.17 | 719 ± 3 | 1031 ± 272 | 1104 | 73 ± 1 | 45 ± 1 |
| ***Single mutants*** | | | | | | | | | | | | | |
| 113 | far | V113I | 5 | 1 | 1 | 44 | 702 | 1,91 | 719 | 1291 | 515 | 74 | 42 |
| 127 | far | A127T | 1 | 1 | 1 | 102 | 703 | 2,49 | 717 | 1224 | 1492 | 75 | 44 |
| 158 | Dist. D-pocket | A158K | 5 | 1 | 1 | 51 | 702 | 2,04 | 716 | 1350 | 681 | 74 | 44 |
| 166 | cis acting | M166L | 5 | 1 | 1 | 70 | 704 | 2,69 | 718 | 1280 | 1160 | 78 | 43 |
| 166 | cis acting | M166V | *no measurable chromophorylation* | | | | | | | | | | |
| 166 | cis acting | M166C | 1 | 1 | 1 | 110 | 706 | 2,60 | 723 | 553 | 764 | 74 | 46 |
| 168 | trans acting | Y168L | 1 | 5 | 1 | 67 | 702 | 3,11 | 719 | 777 | 748 | 70 | 41 |
| 168 | trans acting | Y168T | 2 | 2 | 3 | 119 | 700 | 1,59 | 720 | 2667 | 2335 | 14 | 7 |
| 177 | Dist. D-pocket | E177S | 1 | 1 | 1 | 72 | 703 | 2,36 | 720 | 1297 | 1058 | 75 | 45 |
| 178 | Dist. D-pocket | V178L | 5 | 1 | 1 | 12 | 701 | 0,69 | 719 | 2883 | 114 | 35 | 27 |
| 178 | Dist. D-pocket | V178A | 4 | 1 | 3 | 79 | 698 | 0,85 | 717 | 1005 | 327 | 66 | 18 |
| 190 | Dist. D-pocket | F190W | 1 | 1 | 1 | 30 | 703 | 0,64 | 719 | 965 | 99 | 70 | 41 |
| 190 | Dist. D-pocket | F190Y | 1 | 1 | 1 | 98 | 703 | 2,63 | 720 | 902 | 1152 | 80 | 42 |
| 190 | Dist. D-pocket | F190I | 5 | 1 | 1 | 70 | 700 | 1,30 | 720 | 1058 | 462 | 81 | 41 |
| 190 | Dist. D-pocket | F190L | 1 | 1 | 1 | 94 | 704 | 2,41 | 720 | 958 | 1039 | 79 | 48 |
| 190 | Dist. D-pocket | F190T | 5 | 1 | 1 | 90 | 703 | 1,69 | 718 | 749 | 547 | 85 | 44 |
| 190 | Dist. D-pocket | F190M | 1 | 1 | 1 | 52 | 704 | 2,15 | 721 | 826 | 444 | 69 | 46 |
| 191 | Dist. D-pocket | L191V | 1 | 1 | 1 | 118 | 703 | 2,49 | 722 | 702 | 987 | 82 | 45 |
| 195 | trans acting | Y195F | 1 | 1 | 1 | 120 | 703 | 2,35 | 719 | 217 | 285 | 7 | 42 |
| 199 | trans acting | D199L | 2 | 4 | 2 | 71 | 699 | 1,45 | 718 | 3708 | 1830 | 25 | 11 |
| 200 | middle | I200L | 1 | 1 | 1 | 34 | 700 | 1,40 | 717 | 1336 | 301 | 84 | 47 |
| 200 | middle | I200V | 1 | 1 | 1 | 111 | 701 | 2,37 | 719 | 1175 | 1477 | 77 | 47 |
| 203 | Dist. A-pocket | Q203N | 1 | 1 | 1 | 111 | 703 | 2,45 | 719 | 1105 | 1446 | 71 | 40 |
| 204 | middle | A204S | 1 | 1 | 1 | 114 | 701 | 2,48 | 718 | 1254 | 1697 | 69 | 42 |
| 208 | C-Prop | Y208F | 1 | 1 | 1 | 83 | 702 | 1,56 | 722 | 655 | 397 | 70 | 37 |
| 208 | C-Prop | Y208H | 1 | 1 | 1 | 98 | 703 | 2,69 | 724 | 612 | 747 | 75 | 45 |
| 214 | C-Prop | R214G | 1 | 1 | 1 | 81 | 703 | 1,99 | 723 | 773 | 598 | 71 | 42 |
| 214 | C-Prop | R214L | 1 | 1 | 1 | 82 | 700 | 2,51 | 721 | 537 | 530 | 71 | 44 |
| 245 | B-Prop | R245A | 2 | 1 | 1 | 16 | 698 | 1,46 | 712 | 997 | 113 | 56 | 28 |
| 245 | B-Prop | R245K | 1 | 1 | 1 | 128 | 699 | 2,01 | 717 | 908 | 1122 | 81 | 36 |
| 245 | B-Prop | R245S | 2 | 1 | 3 | 76 | 696 | 0,70 | 715 | 1058 | 268 | 66 | 29 |
| 245 | B-Prop | R245L | *no measurable chromophorylation* | | | | | | | | | | |
| 245 | B-Prop | R245I | *no measurable chromophorylation* | | | | | | | | | | |
| 245 | B-Prop | R245H | 1 | 1 | 1 | 48 | 700 | 1,73 | 717 | 1103 | 443 | 72 | 37 |
| 245 | B-Prop | R245E | *no measurable chromophorylation* | | | | | | | | | | |
| 245 | B-Prop | R245Q | 4 | 5 | 3 | 19 | 694 | 0,67 | 715 | 724 | 44 | 65 | 16 |
| 247 | B-Prop | V247T | 1 | 1 | 1 | 86 | 702 | 2,26 | 718 | 1055 | 989 | 79 | 47 |
| 250 | Dist. A-pocket | I250M | 1 | 1 | 1 | 87 | 702 | 2,46 | 723 | 690 | 711 | 76 | 44 |
| 250 | Dist. A-pocket | I250N | *no measurable chromophorylation* | | | | | | | | | | |
| 251 | B-Prop | H251T | 3 | 3 | 2 | 28 | 696 | 0,41 | 718 | 640 | 35 | 78 | 32 |
| 254 | trans acting | Y254F | 1 | 2 | 1 | 76 | 703 | 1,92 | 719 | 2816 | 1913 | 26 | 18 |
| 255 | middle | L255V | 1 | 1 | 1 | 84 | 702 | 2,54 | 722 | 719 | 741 | 82 | 48 |
| 258 | cis acting | M258F | 1 | 1 | 1 | 74 | 704 | 2,26 | 722 | 629 | 506 | 75 | 44 |

**Supplementary Table 1 Continuation**

| AA Position | Group | Mutation | Cluster $P_r$ | Cluster $P_{fr}$ | Cluster Diff. | Weighted Soret to 280 (a.u.) | Q peak in Pr (nm) | Q to Soret (a.u.) | Fluo. peak (nm) | Corr. fluo. signal (a.u.) | Brigthness (a.u.) | Δ fluo. (%) | Δ abs. ~700 nm (%) |
|---|---|---|---|---|---|---|---|---|---|---|---|---|---|
| 261 | Dist beta | G261R | 1 | 1 | 1 | 137 | 703 | 2,78 | 723 | 796 | 1456 | 72 | 45 |
| 261 | Dist beta | G261S | 2 | 1 | 3 | 110 | 700 | 1,46 | 724 | 617 | 478 | 66 | 25 |
| 263 | C-Prop | S263A | 1 | 1 | 1 | 53 | 703 | 2,54 | 718 | 1236 | 795 | 75 | 45 |
| 264 | Dist beta | L264A | 1 | 1 | 1 | 43 | 703 | 1,46 | 718 | 878 | 264 | 71 | 43 |
| 264 | Dist beta | L264V | 1 | 1 | 1 | 73 | 702 | 1,51 | 720 | 839 | 445 | 62 | 35 |
| 265 | C-Prop | S265A | 3 | 3 | 3 | 58 | 685 | 0,32 | 712 | 247 | 22 | 19 | 0 |
| 273 | Dist. D-pocket | K273Q | 1 | 1 | 1 | 83 | 703 | 2,49 | 720 | 785 | 777 | 76 | 45 |
| 273 | Dist. D-pocket | K273E | 1 | 1 | 1 | 105 | 703 | 2,64 | 726 | 536 | 718 | 75 | 45 |
| 275 | Dist. D-pocket | W275F | 1 | 1 | 1 | 65 | 703 | 1,97 | 725 | 777 | 423 | 73 | 42 |
| 279 | cis acting | A279S | 1 | 1 | 1 | 75 | 701 | 2,00 | 717 | 1122 | 810 | 73 | 42 |
| 279 | cis acting | A279V | 5 | 1 | 1 | 41 | 696 | 0,87 | 716 | 1254 | 215 | 84 | 45 |
| 281 | cis acting | H281T | 1 | 2 | 1 | 67 | 700 | 2,12 | 718 | 1223 | 836 | 53 | 30 |
| 281 | cis acting | H281R | 4 | 3 | 2 | 88 | 701 | 0,65 | 717 | 1402 | 386 | 88 | 50 |
| 281 | cis acting | H281V | 1 | 2 | 1 | 68 | 703 | 2,34 | 719 | 432 | 330 | 28 | 25 |
| 281 | cis acting | H281D | 1 | 1 | 1 | 92 | 701 | 2,01 | 717 | 1421 | 1259 | 76 | 43 |
| 281 | cis acting | H281L | 1 | 1 | 1 | 94 | 706 | 2,43 | 716 | 297 | 325 | 63 | 46 |
| 281 | cis acting | H281A | 1 | 2 | 1 | 63 | 703 | 2,62 | 722 | 707 | 560 | 45 | 26 |
| 284 | Dist. D-pocket | E284D | 1 | 1 | 1 | 88 | 703 | 2,45 | 720 | 777 | 806 | 76 | 45 |
| 286 | Dist. D-pocket | R286Y | 1 | 1 | 1 | 44 | 703 | 1,29 | 720 | 1095 | 306 | 75 | 45 |
| 458 | Phy tongue | R458D | 1 | 1 | 1 | 66 | 703 | 2,24 | 719 | 1023 | 728 | 71 | 44 |
| 458 | Phy tongue | R458M | 1 | 1 | 1 | 80 | 703 | 2,39 | 722 | 740 | 678 | 72 | 44 |
| 462 | Phy tongue | R462A | 1 | 1 | 1 | 92 | 698 | 1,63 | 719 | 1611 | 1157 | 70 | 35 |
| 462 | Phy tongue | R462K | 1 | 1 | 1 | 70 | 699 | 1,87 | 718 | 1628 | 1022 | 69 | 36 |
| 465 | Phy tongue | F465W | 1 | 1 | 1 | 119 | 698 | 2,28 | 714 | 2583 | 3754 | 69 | 36 |
| 465 | Phy tongue | F465Y | 1 | 1 | 1 | 66 | 700 | 1,66 | 715 | 2277 | 1238 | 70 | 39 |
| 465 | Phy tongue | F465I | 1 | 1 | 1 | 90 | 698 | 1,85 | 719 | 1824 | 1457 | 74 | 42 |
| | | ***Double mutants*** | | | | | | | | | | | |
| | | Y254F + G261R | 1 | 2 | 1 | 160 | 703 | 2,44 | 720 | 3054 | 5539 | 18 | 15 |
| | | G261R + F465W | 1 | 1 | 1 | 72 | 698 | 1,72 | 716 | 2996 | 1981 | 64 | 33 |
| | | ***Truncations of wildtype ReBphP-PCM*** | | | | | | | | | | | |
| | | ΔPhy-290 | 2 | 3 | 2 | 61 | 697 | 0,68 | 716 | 2689 | 207 | 72 | 46 |
| | | ΔPhy-313 (**photoswitching fluorescent variant**) | 2 | 3 | 2 | 470 | 698 | 1,39 | 717 | 3177 | 3865 | 80 | 54 |
| | | ΔPhy-335 | 4 | 3 | 2 | 295 | 698 | 0,91 | 719 | 1667 | 893 | 76 | 49 |
| | | ***Truncated variants*** | | | | | | | | | | | |
| | | ΔPhy-306 + D199F + Y254F + G261R | 2 | 4 | 4 | 569 | 699 | 1,57 | 718 | 5257 | 8013 | 8 | 7 |
| | | ΔPhy-313 + D199F + A217P + Y254F + G261R | 2 | 4 | 4 | 493 | 699 | 1,48 | 717 | 5069 | 6848 | 7 | 4 |
| | | ΔPhy-335 + F190Y | 2 | 3 | 2 | 164 | 697 | 1,08 | 717 | 2814 | 1068 | 68 | 47 |
| | | ΔPhy-335 + Y254F + G261R | 2 | 4 | 2 | 321 | 699 | 1,46 | 719 | 4397 | 3831 | 24 | 18 |
| | | ΔPhy-335 + D199L + Y254F + G261R | 2 | 4 | 2 | 297 | 695 | 1,72 | 714 | 5797 | 5497 | 21 | 14 |
| | | ΔPhy-313 + D199F + Y254F + G261R (**bright fluorescent variant**) | 2 | 4 | 4 | 567 | 699 | 1,54 | 716 | 5329 | 8656 | 12 | 5 |
| | | Δ1-9 + ΔPhy-313 + D199F + Y254F + G261R | 2 | 4 | 3 | 730 | 698 | 1,61 | 718 | 5373 | 11713 | 12 | 7 |

**Supplementary Table 2:** Analysis of chromophore planarity. According to the out-of-plane RMSD values, all *trans*-chromophores exhibit lower planarity compared to their *cis*-counterparts. Notably, the ΔPHY313 *trans* variant is the least planar overall. This distortion correlates with the observed torsion at the B–C methine bridge, which is likely driven by the altered orientation of the C-propionate group in this truncated variant. Interestingly, the ΔPHY313 *cis* variant appears to be the most planar chromophore based on the global RMSD. However, analysis of the individual methine bridge torsions reveals that the ΔPHY313+D199F+R254F+G261R *cis* variant shows bridge geometries closer to ideal planarity.

| Variant/Conformation | Planarity (RMSD) | Torsions A, B | Torsions B, C | Torsions C, D |
|---|---|---|---|---|
| ***cis*** | | | | |
| ReBphP-PCM | **0.3414 Å** | $\Theta_{A,B}$ = 20.22°<br>$\Phi_{A,B}$ =-162.2°<br>$\Psi_{A,B}$ = 177.3° | $\Theta_{B,C}$ = 12.11°<br>$\Phi_{B,C}$ =-171.2°<br>$\Psi_{B,C}$ = 178.9° | $\Theta_{C,D}$ = 41.41°<br>$\Phi_{C,D}$ = 28.5°<br>$\Psi_{C,D}$ =-160.1° |
| ΔPhy-313+D199F<br>+R254F+G261R | **0.3170 Å** | $\Theta_{A,B}$ = 17.42°<br>$\Phi_{A,B}$ =-173.6°<br>$\Psi_{A,B}$ =-172.0° | $\Theta_{B,C}$ = 10.21°<br>$\Phi_{B,C}$ =-176.3°<br>$\Psi_{B,C}$ = 178.1° | $\Theta_{C,D}$ = 33.06°<br>$\Phi_{C,D}$ = 22.7°<br>$\Psi_{C,D}$ =-160.2° |
| ΔPhy-313 | **0.2973 Å** | $\Theta_{A,B}$ = 16.92°<br>$\Phi_{A,B}$ =-171.9°<br>$\Psi_{A,B}$ = 158.7° | $\Theta_{B,C}$ = 12.74°<br>$\Phi_{B,C}$ =-171.2°<br>$\Psi_{B,C}$ = 177.8° | $\Theta_{C,D}$ = 43.50°<br>$\Phi_{C,D}$ = 33.8°<br>$\Psi_{C,D}$ =-160.3 |
| ***trans*** | | | | |
| ReBphP-PCM | **0.3734 Å** | $\Theta_{A,B}$ = 32.49°<br>$\Phi_{A,B}$ =-152.5°<br>$\Psi_{A,B}$ = 179.4° | $\Theta_{B,C}$ = 10.79°<br>$\Phi_{B,C}$ = 172.2°<br>$\Psi_{B,C}$ = 172.7° | $\Theta_{C,D}$ = 54.4°<br>$\Phi_{C,D}$ = 23.1°<br>$\Psi_{C,D}$ = 41.7° |
| ΔPhy-313+D199F<br>+R254F+G261R | **0.3771 Å** | $\Theta_{A,B}$ = 33.40°<br>$\Phi_{A,B}$ =-159.7°<br>$\Psi_{A,B}$ =-173.8° | $\Theta_{B,C}$ = 9.24°<br>$\Phi_{B,C}$ = 174.8°<br>$\Psi_{B,C}$ = 171.3° | $\Theta_{C,D}$ = 49.05°<br>$\Phi_{C,D}$ = 34.4°<br>$\Psi_{C,D}$ = 26.3° |
| ΔPhy-313 | **0.3843 Å** | $\Theta_{A,B}$ = 32.84°<br>$\Phi_{A,B}$ =-151.7°<br>$\Psi_{A,B}$ = 179.6° | **$\Theta_{B,C}$ = 21.85°**<br>$\Phi_{B,C}$ = 176.1°<br>$\Psi_{B,C}$ = **158.5°** | $\Theta_{C,D}$ = 50.58°<br>$\Phi_{C,D}$ = 29.7°<br>$\Psi_{C,D}$ = 34.1° |

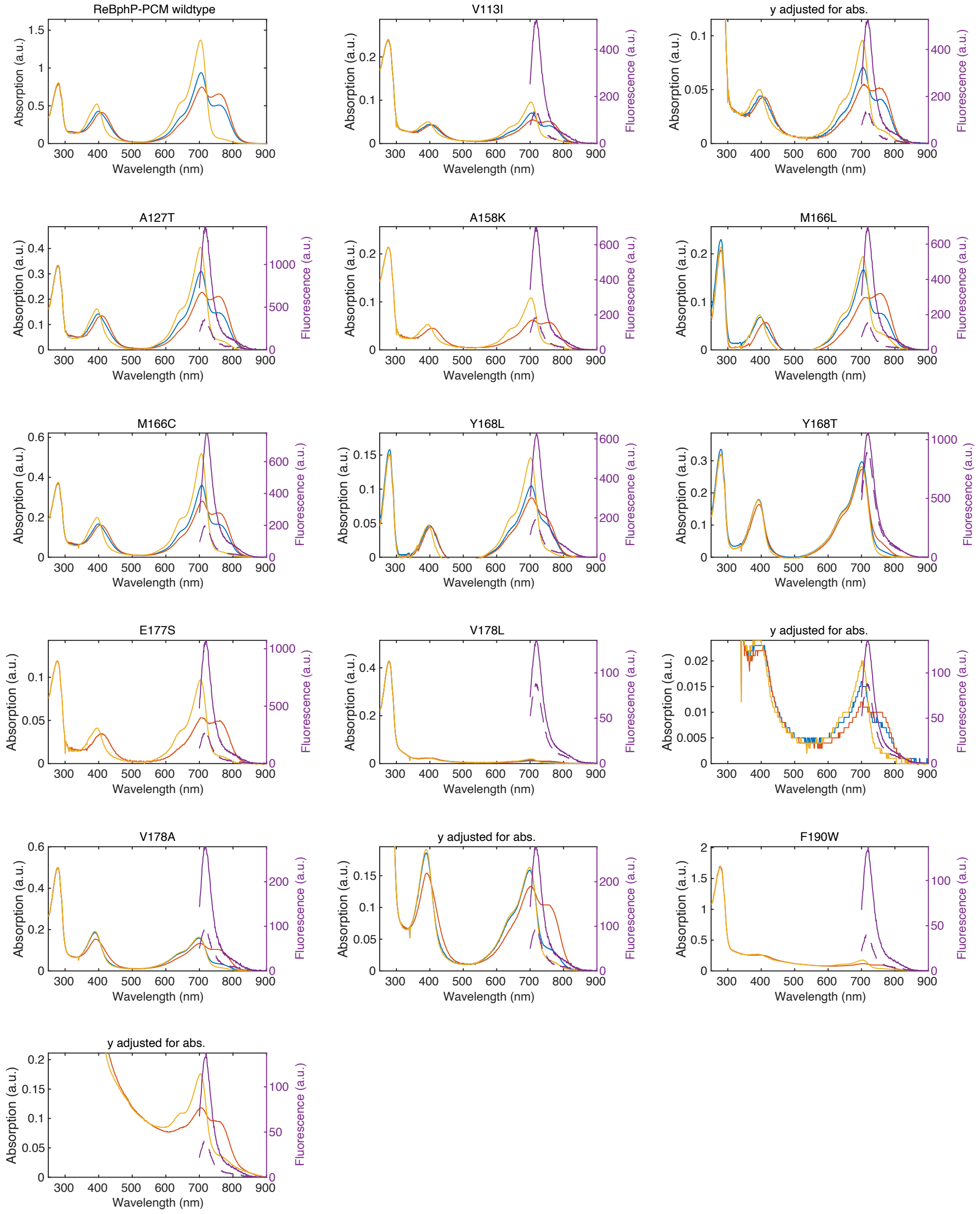


**Supplementary Figure 1:** Raw spectral data of variants measured in this study. Spectra are ordered as in **Suppl. Table 1**. Shown are the raw absorption spectra of the purified protein in its equilibrium state without illumination (blue) in its $P_r$ state after 780 nm illumination (orange) and its $P_{fr}$ state after 660 nm illumination (red). Spectra are normalized for the absorption at 280 nm. For selected variants, the fluorescence is shown (right y axis) in the $P_r$ (solid line) and $P_{fr}$ (dashed line). The fluorescence is normalized to the peak absorption of the 700 nm Q band in the $P_r$ state of the respective variant. Hence, it reflects the fluorescence of a given variant but not the absolute brightness. Generally, for all variants the absorption spectra are shown such, that all peaks of the spectrum are visible. For variants with a poor absorption of the chromophore peaks (e.g. due to poor chromophorylation) the following panel shows the absorption spectra with the y axis adjusted such as to show the chromophore peaks clearly. The fluorescence spectra remain unaltered. For ReBphP-PCM wildtype an exemplary spectrum is shown, see **Suppl. Figure 4 and 5** for wild-type spectra of several purifications to assess variance.

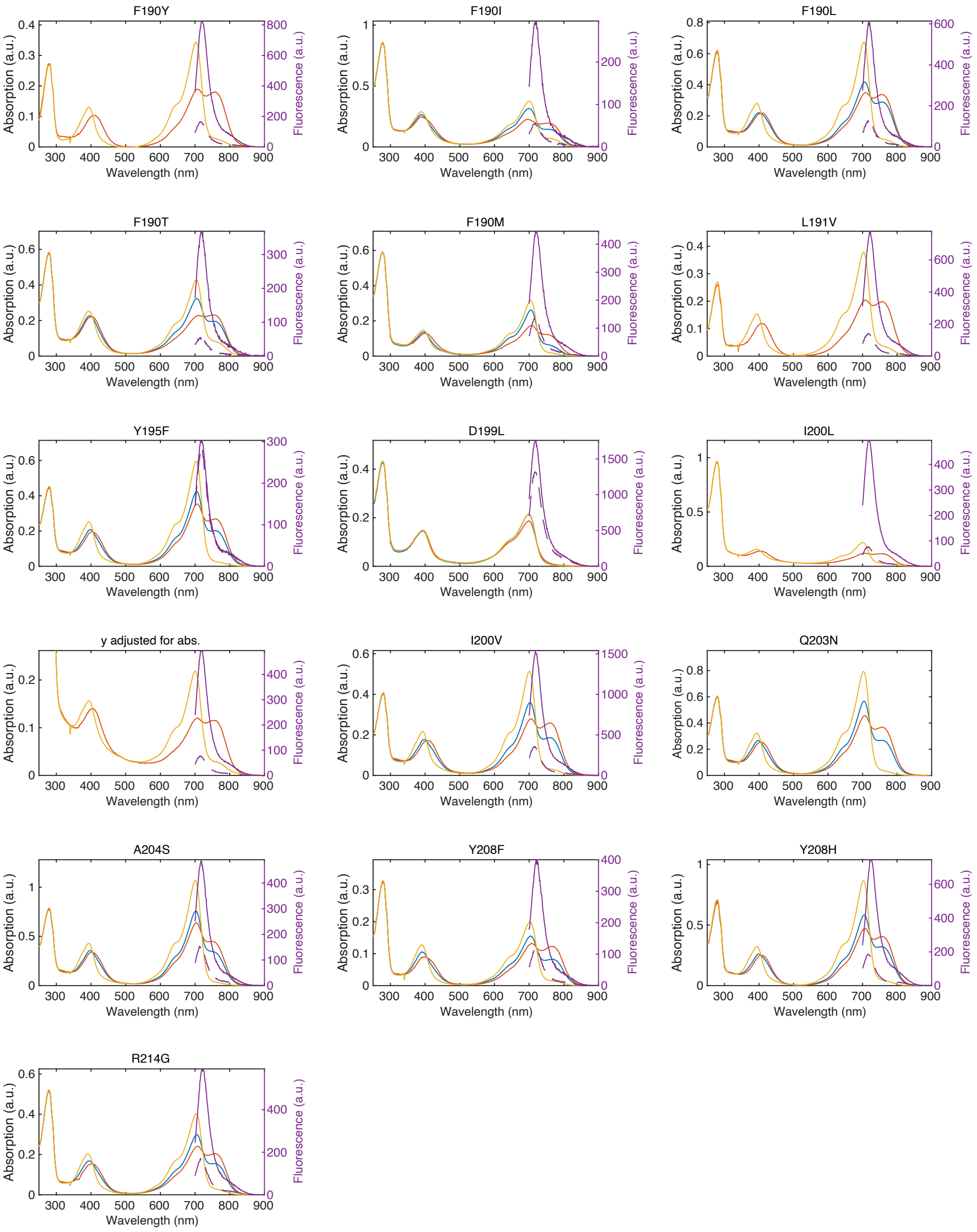


**Continuation of Supplementary Figure 1**

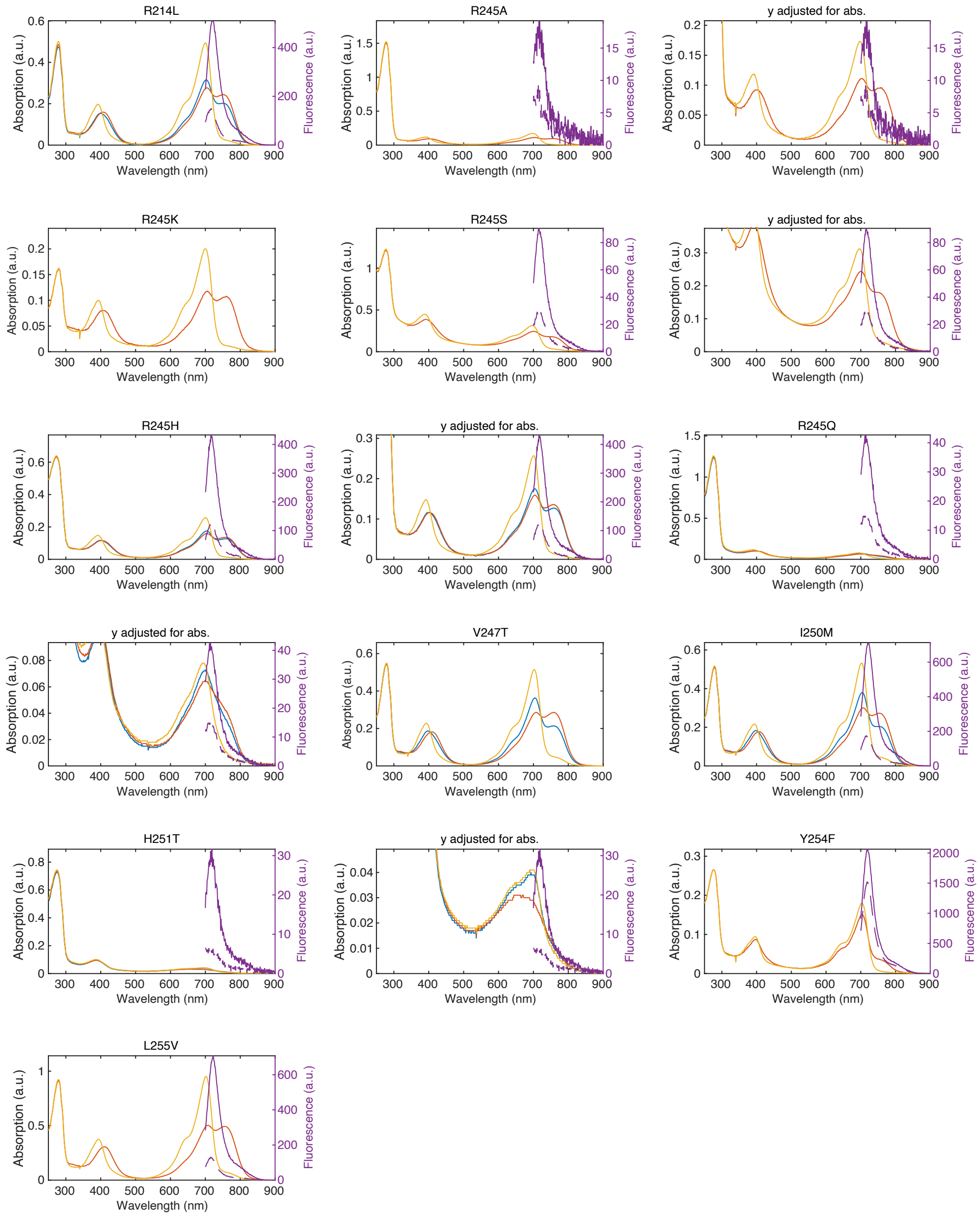


**Continuation of Supplementary Figure 1**

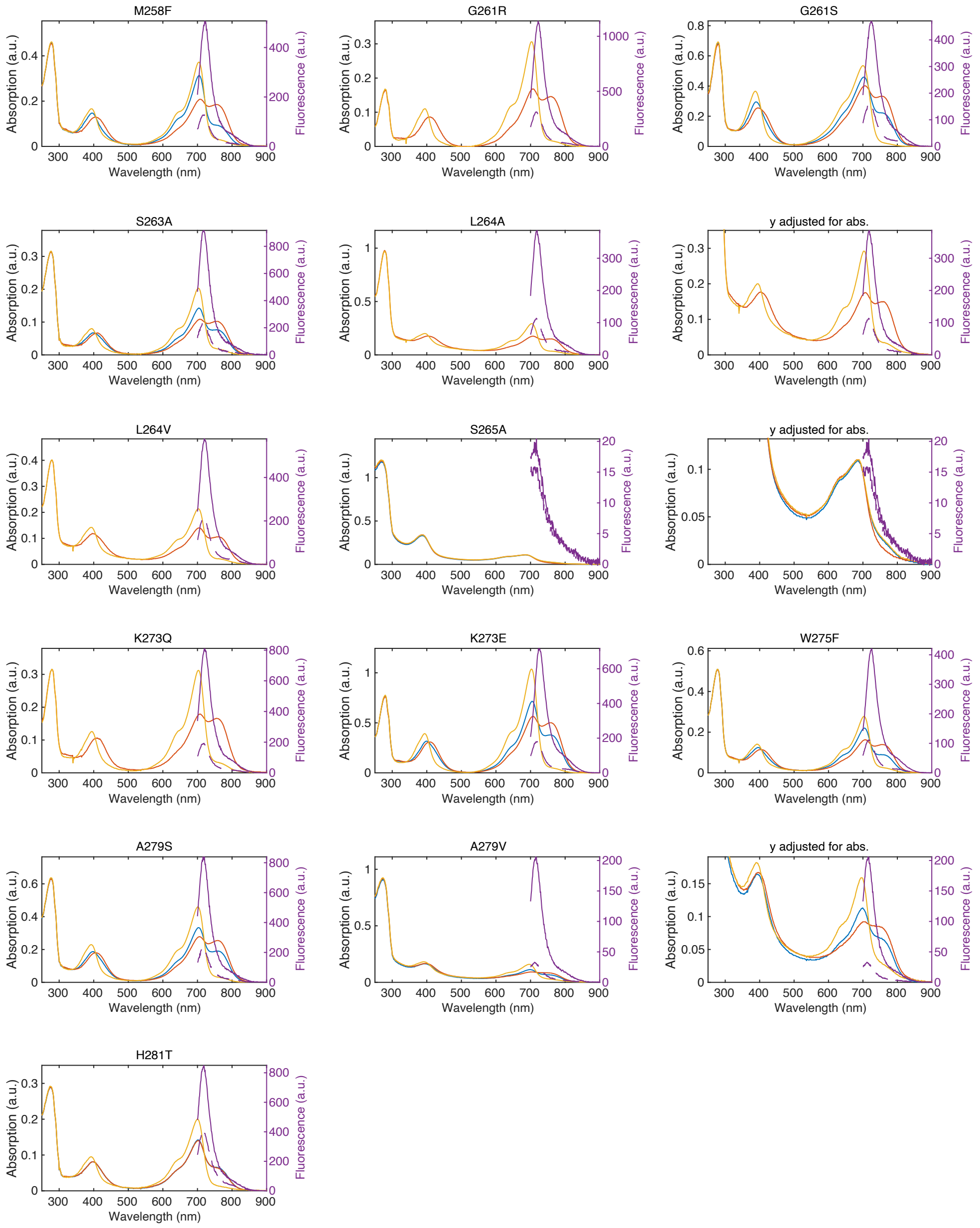


**Continuation of Supplementary Figure 1**

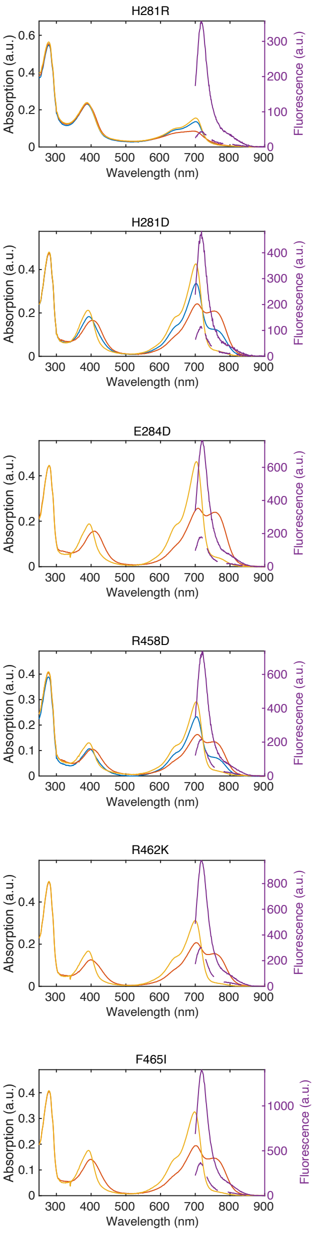


**Continuation of Supplementary Figure 1**

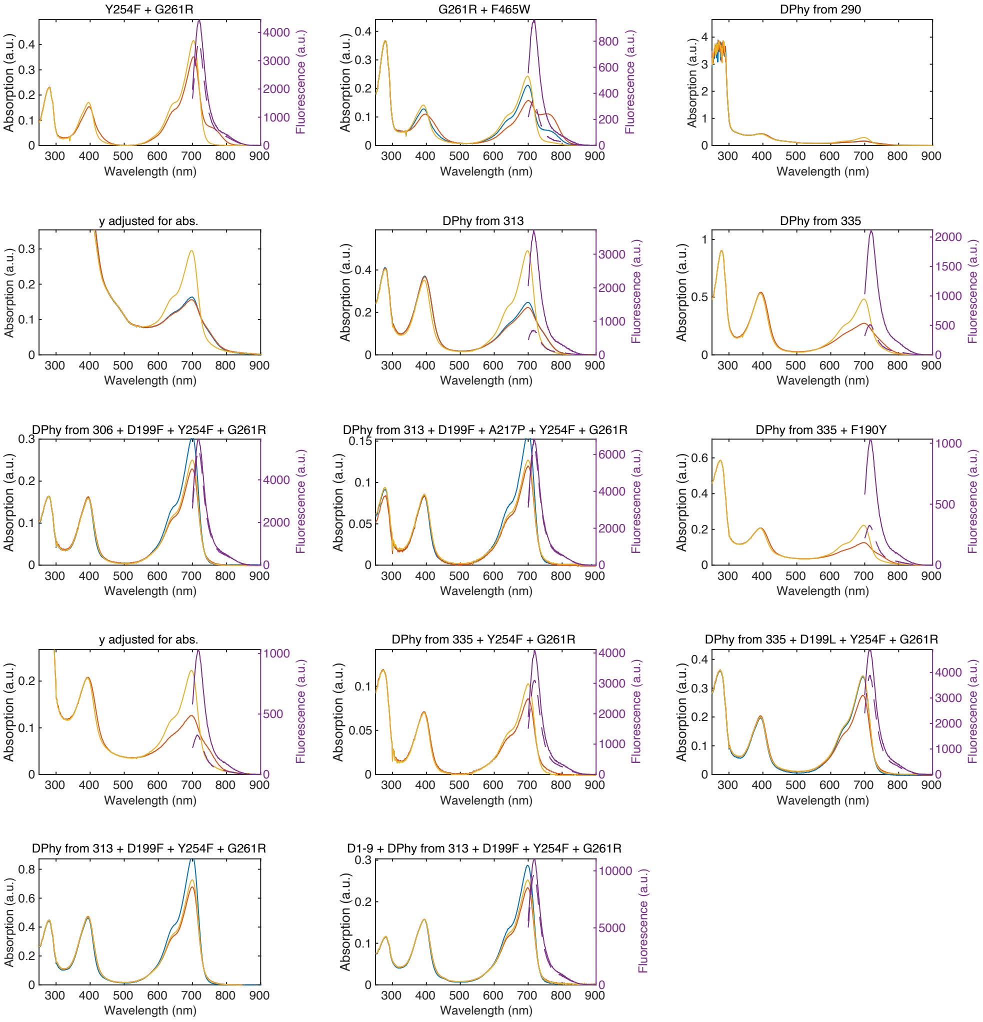


**Continuation of Supplementary Figure 1**

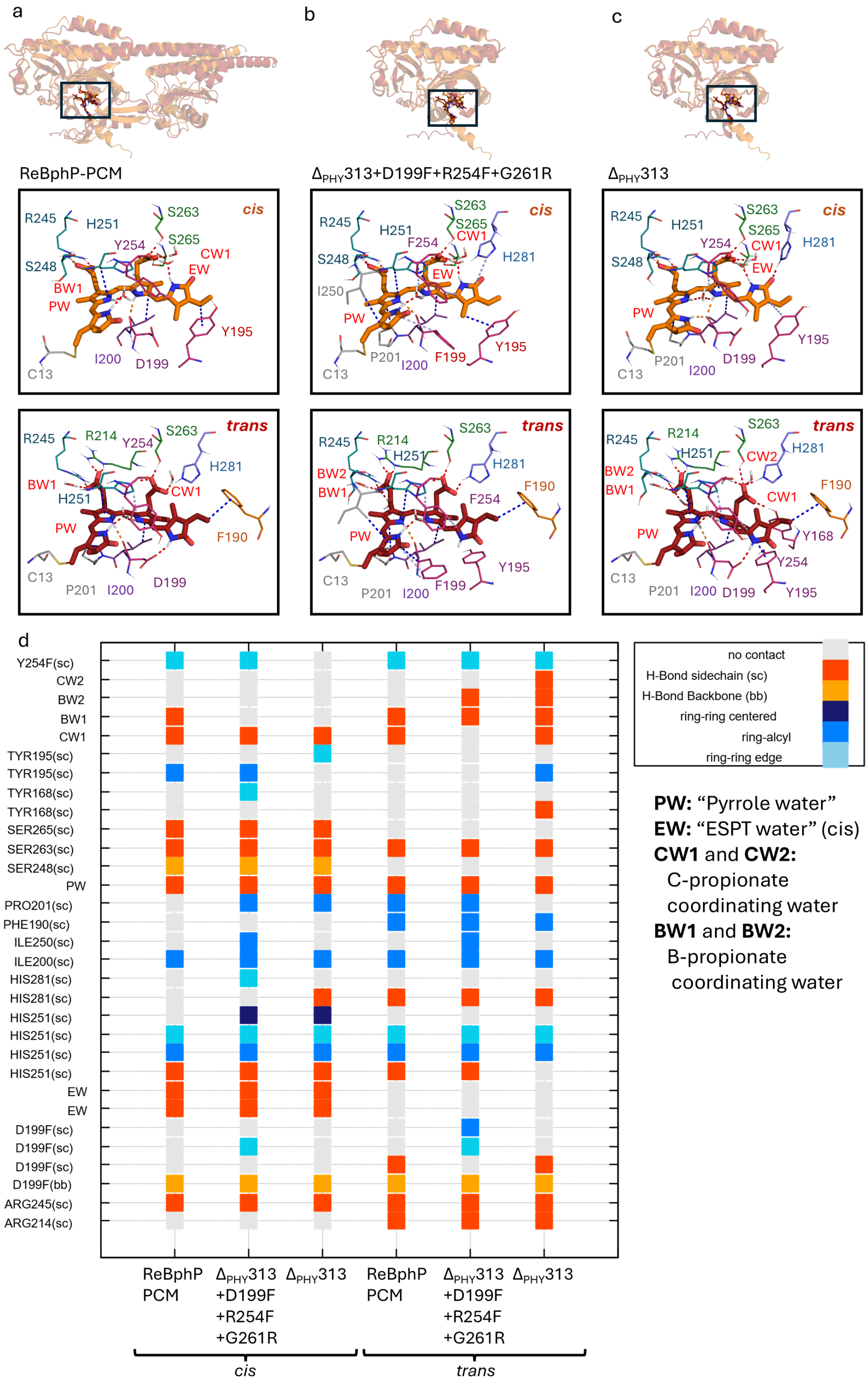


**Supplementary Figure 2:** Structural models and interaction networks of ReBphP variants. For clarity, not all van der Waals interactions and solvent contacts are displayed. **(a–c)** Structural overview and detailed view of interactions (dashed lines) of the chromophore in **(a)** ReBphP-PCM, **(b)** the $\Delta_{PHY}$313+D199F+R254F+G261R variant, and **(c)** $\Delta_{PHY}$313, featuring the biliverdin chromophore in assumed *cis* and *trans* conformations. **(d)** Contact map summarizing the non-covalent environment. Interactions are categorized into hydrogen bonds and van der Waals forces, including π-stacking (ring–ring) and hydrophobic (ring–alkyl) contacts. The "pyrrole water" (PW), a conserved structural element coordinated by the pyrrole nitrogen atoms, is shown in accordance with[3]. The term "ESPT water" (EW) refers to a specific water molecule positioned to facilitate the excited-state proton transfer pathway, observed exclusively in the *cis* conformation[14].

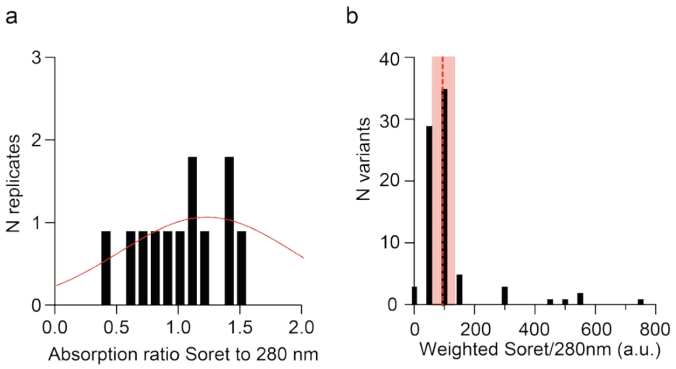


**Supplementary Figure 3:** Chromophorylation as estimated by the ratio between Soret band and 280 nm peak. **a** The variance of chromophorylation is comparably huge as visible by a comparison of Soret to 280 nm absorption for all ReBphP-PCM preparations (N=13). Red line indicates a gaussian fit to the data normalized by the mean. **b** Overview of all variants, most of the variants lie within the standard deviation (shaded red) of ReBphP-PCM (dashed red lines). Variants having either better or worse chromophorylation are indicated.

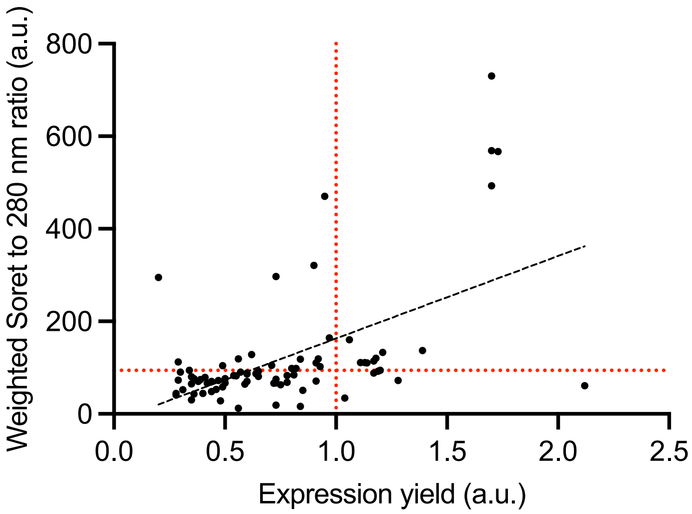


**Supplementary Figure 4:** Correlation of expression yield and chromophorylation. The expression yield is tentatively determined as the yield of the IMAC purification step in comparison with the wildtype reference of that purification batch (ReBphP-PCM wildtype = 1). This determination of expression is relatively imprecise (differences in inoculation, induction, cell cracking etc.) hence in the table (Suppl. Table 1) we only refer to good, wildtype like expression “+++” > 0.66, mediocre “++” > 0.33 < 0.66 and poor “+” < 0.33. Chromophorylation is determined as the ration between Soret and 280 nm peak (ReBphP-PCM wildtype ~ 0.4). The black dashed line indicates the linear relationship.

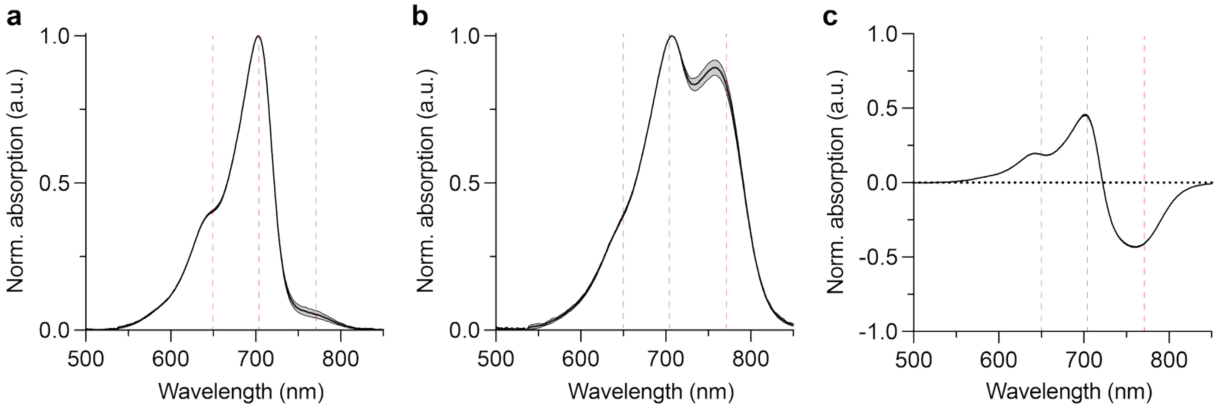


**Supplementary Figure 5:** Mean spectra of purified wildtype ReBphP-PCM in their $P_r$ (**a**) and $P_{fr}$ (**b**) states as well as the $P_r$-$P_{fr}$ difference spectra (**c**). Standard deviation is shown as shaded area of N=13 independent purifications and measurements.

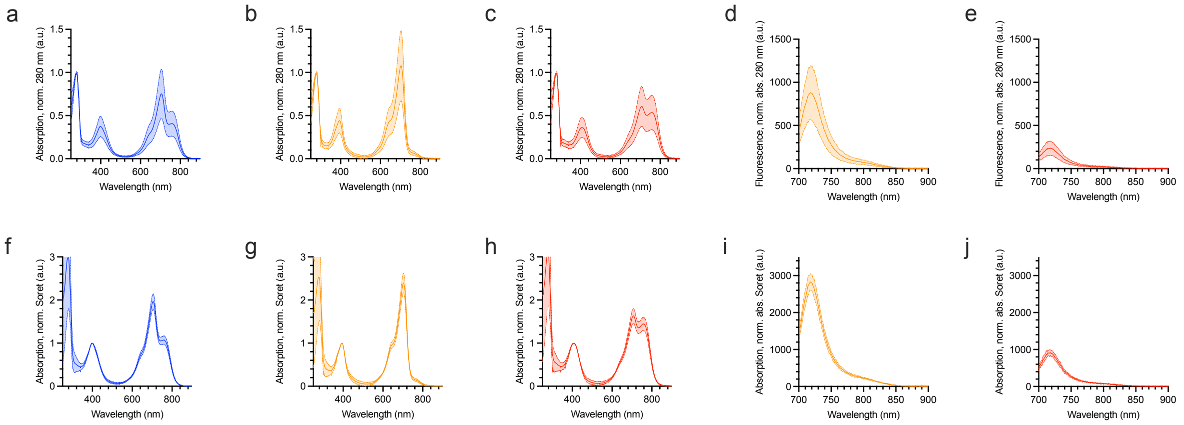


**Supplementary Figure 6:** Variance of independent preparations and measurements of ReBphP-PCM. Shown are the absorption spectra (a-c and f-h) of 12 and the fluorescence spectra (d,e,i and j) of 4 replicates as mean (solid line) and standard deviation (shaded area). The spectra are either normalized to the absorption at 280 nm (a-e) or to the absorption of the Soret band (f-j). While the former normalization includes the variance of the chromophorylation the latter solely reflects the variance of the spectral features. The variance of chromophorylation is comparably huge as visible by a comparison of Soret to 280 nm absorption for all preparations (**Suppl. Fig. 2a**).

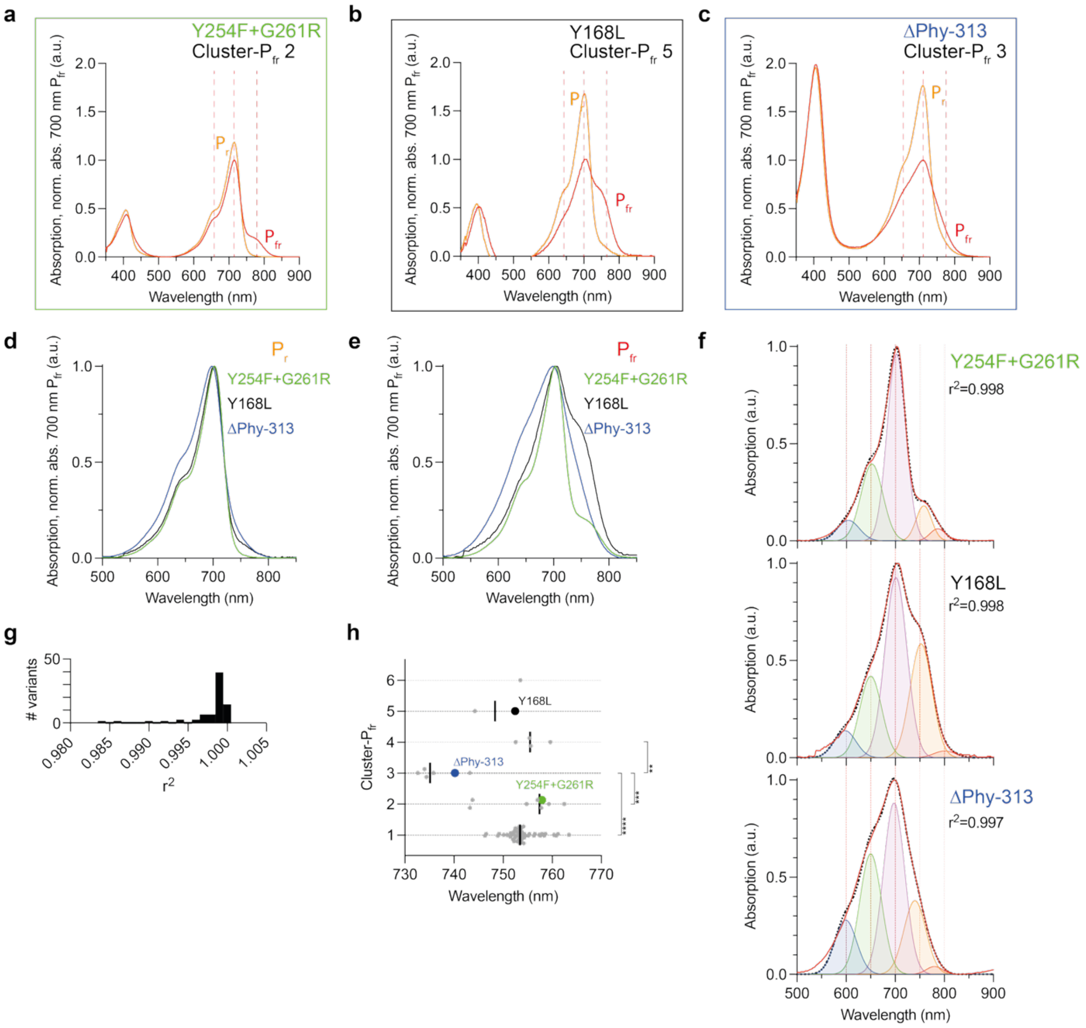

**Supplementary Figure 7:** Raw absorption spectra of $P_r$ and $P_{fr}$ of variants Y254F+R261G (**a**) Y168L (**b**) and ΔPhy-313 (**c**). For an easier comparison of the Q bands after baseline subtraction, d-e show only the Q-band in $P_r$ (**d**) and $P_{fr}$ state (**e**) of Y254F+R261G (green), Y168L (black) and ΔPhy-313 (blue). **f** Subband decomposition for the baseline corrected $P_{fr}$ spectra of Y254F+R261G, Y168L and ΔPhy-313. A model was chosen that uses only a reduced number of components likely not fully reflecting all transitions realistically. The model used the following band positions with their degree of freedom 600 nm ± 10 nm, 660 nm ± 10 nm, 700 nm ± 10 nm, 760 nm ± 40 nm, 800 nm ± 20 nm, the ~760 nm band was deliberately chosen with a larger freedom to allow to accommodate the blue and red-shifted positions. **g** All fits shows a very high quality ($r^2$). **h** Position of the ~760 nm like spectral band for the model described in g for all clusters-$P_{fr}$. Shown are the individual values together with their mean. The values for the two variants shown in this figure are shown with their color. Statistics analysis as Mann-Whitney test (p-value **** > 0.0001, *** > 0.001, ** > 0.01, analysis for clusters 5 and 6 was omitted due to lack of datapoints).

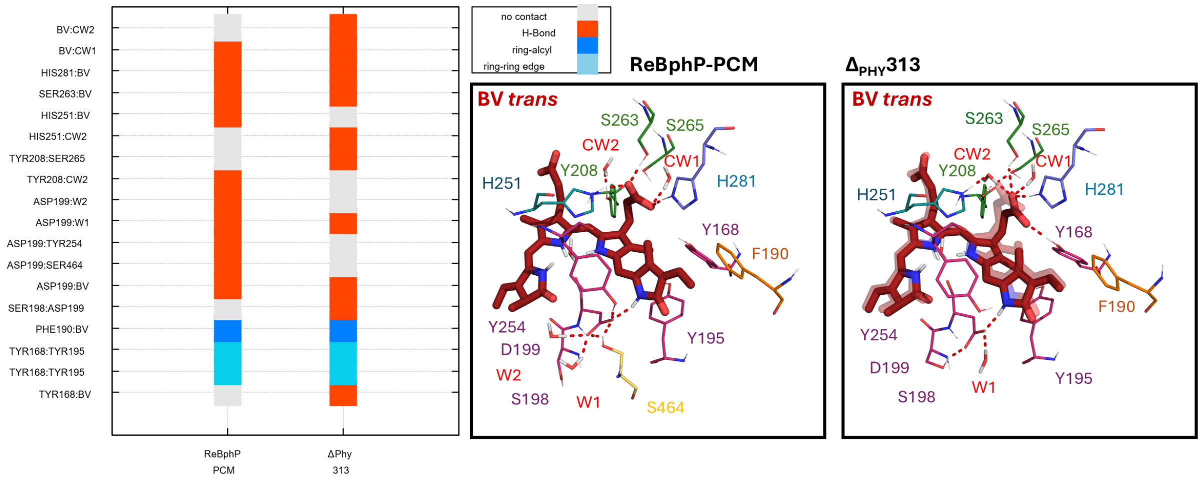


**Supplementary Figure 8:** Potential reasons for C-propionate and changed water network between *trans* states of ReBphP-PCM and $\Delta_{PHY}$313. Same labeling and representations as in **Suppl. Figure 2**. In ReBphP-PCM, D199 is anchored by an interaction with S464 of the PHY-tongue. Its position is further rigidified by a hydrogen-bonding network involving Y254 and two water molecules. In the truncated $\Delta_{PHY}$313 variant, the absence of the PHY-tongue enables the D199 side chain to reorient toward S198. While D199 maintains its interaction with the biliverdin (BV) D-ring, this reorientation exerts a 'pull' on the pyrrole, potentially altering the alignment of the C-ring and its associated C-propionate. This shift in the C-ring is coupled with a reorganization of the local contact residues. In ReBphP-PCM, Y208 interacts with the water molecule CW2 (CW: C-propionic water molecule 1 and 2 (see **Suppl. Figure 2**)). In $\Delta_{PHY}$313, Y208 reorients toward S265, causing CW2 to shift its position between the H251 side chain and the BV C-propionate. Consequently, the C-propionate adopts a distinct orientation, stabilized by a new hydrogen bond with the Y168 side chain and a coordinated switch of its interaction with H281 to the alternative carboxylate oxygen.

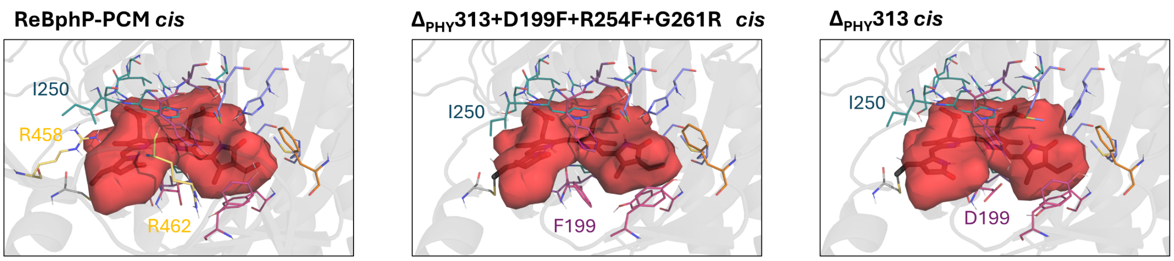


**Supplementary Figure 9:** Packing difference**.** The accessible volume of the biliverdin (BV) binding pocket was calculated for the *cis*-chromophore variants to quantify structural compaction. The ReBphP-PCM variant exhibits a pocket volume of 898 Å³, while the truncated $\Delta_{PHY}$313 shows a slight reduction to 889 Å³. Notably, the mutant $\Delta_{PHY}$313+D199F+R254F+G261R displays a significantly further decreased volume of 826 Å³. These findings correlate with the altered contact patterns presented in **Suppl. Figure 2**, indicating overall tighter π-stacking and van der Waals interactions in the mutant, particularly involving residues P201, H251 and I250. In the truncated variants, the loss of the interaction between I250 and the PHY-tongue residue R458 allows I250 to reorient toward the chromophore, thereby reducing the available pocket space. This compaction is much more pronounced in $\Delta_{PHY}$313+D199F+R254F+G261R than in the $\Delta_{PHY}$313. Further effects of the mutations are the bulkier phenylalanine side chain at position 199, which not only sterically reduces the cavity volume but also stabilizes the chromophore through ring-ring interactions.

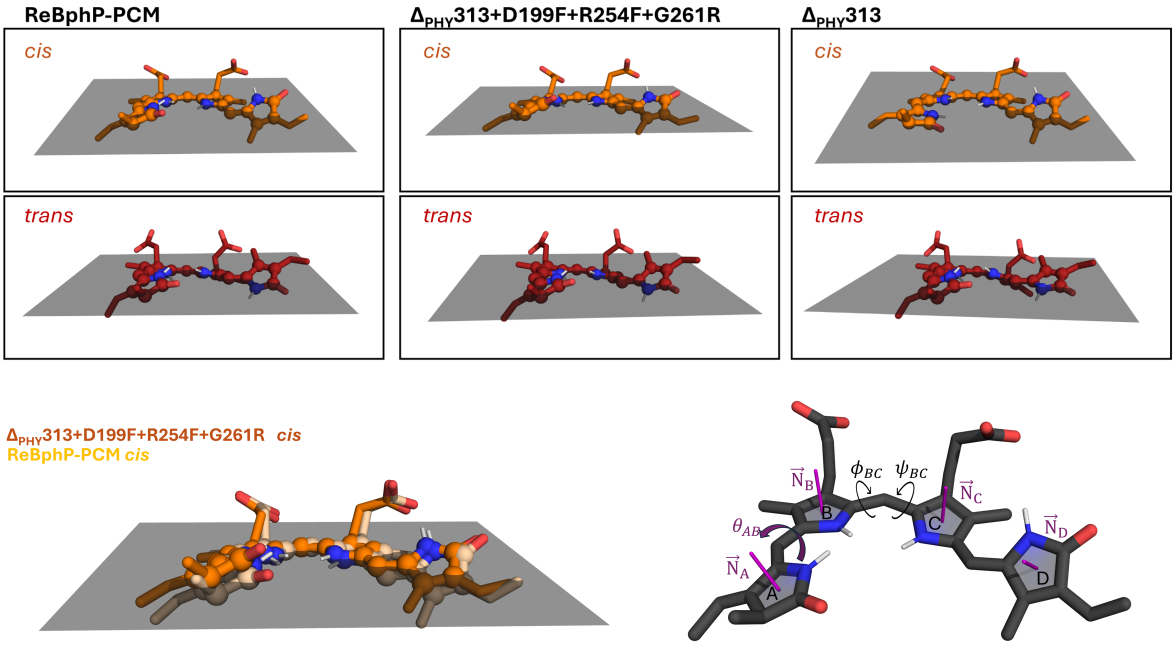


**Supplementary Figure 10:** Planarity in *trans* and *cis* BV conformations. The planarity of the biliverdin backbone was quantified by the root-mean-square deviation (RMSD) from the calculated least-squares plane. This value was derived from the 23 core atoms of the macrocyclic system, comprising the four pyrrole nitrogen atoms, the ring carbon atoms, and the linking methine bridges. All atoms are highlighted as spheres in the structural models. Substituents were excluded from the plane definition to focus the analysis on the global distortion of the conjugated π-system. All values are listed in **Suppl. Table 2**. The bottom figure illustrates the methodology for quantifying chromophore planarity through the measurement of interplanar angles (ϴ, absolute values to quantify loss of planarity) and dihedral angles (φ,ψ).

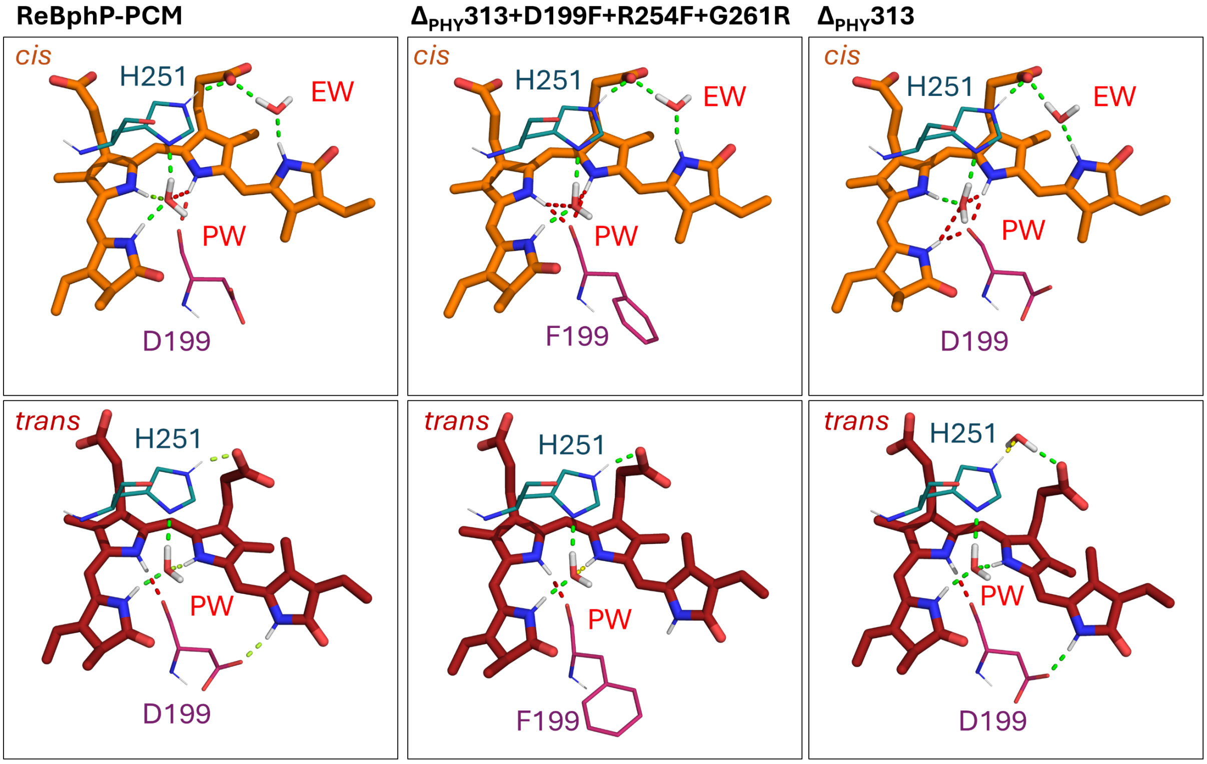


**Supplementary Figure 11:** Possible ESPT routes in *cis* and *trans* configurations. Dashed lines are color-coded in a 'traffic light' fashion to visualize the structural pre-organization for ESPT based on QM/MM ground-state energy minima; while these static models indicate the probability of transfer, actual rates may be further influenced by dynamic fluctuations in the excited state. Following established literature[15,16], interactions are categorized as:

- Green (Highly Probable): d (donor – acceptor) < 2.8 Å and α (donor - H – acceptor) > 160°
- Yellow (Moderate/Borderline): 2.8 ≤ Å d < 2.95 Å and 150° < α < 160°
- Red (Inactive/Blocked): d ≥ 2.95Å and α ≤ 150° or pathways to the peptide backbone.

Hydrogen bonds to the peptide backbone of residue 199 are consistently indicated in red; these represent structural anchors rather than functional ESPT channels due to the low proton affinity of the amide carbonyl.

In all *trans* models, the pyrrole is distinctly oriented towards this backbone oxygen. In the *trans*-$\Delta_{PHY}$313+D199F+R254F+G261R variant, the ESPT pathway via the D199 sidechain is abolished. Furthermore, the $\Delta_{PHY}$313 *trans* variant shows a perturbation of the well-established pyrrole water (PW) pathway due to the reorientation of the C-propionate group (see Suppl. Fig. 8). In this case, an additional water molecule mediates the path between H251 and the C-propionate oxygen, resulting in a slightly less favorable geometry compared to the direct path in other variants.

For the *cis* configurations, all three variants show a high probability for the ESPT route via the D-ring, 'ESPT-water,' and C-propionate. However, significant differences occur regarding the A, B, and C-ring pyrroles. While the $\Delta_{PHY}$313 and the $\Delta_{PHY}$313+D199F+R254F+G261R mutant only exhibit one favorable ESPT pathway via the pyrrole water, the ReBPHP-PCM maintains higher ESPT probabilities for both A- and B-ring pyrroles. This reduction in available ESPT channels in the truncated variants is consistent with the observed increase in fluorescence quantum yields.

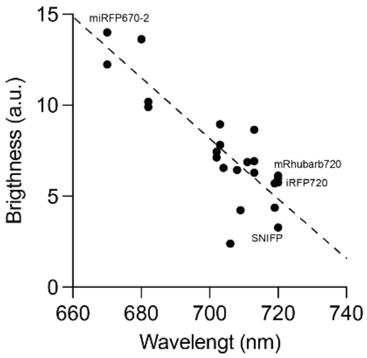


**Supplementary Figure 12:** Relation between brightness and center of the emission peak. Data taken from fpbase.org

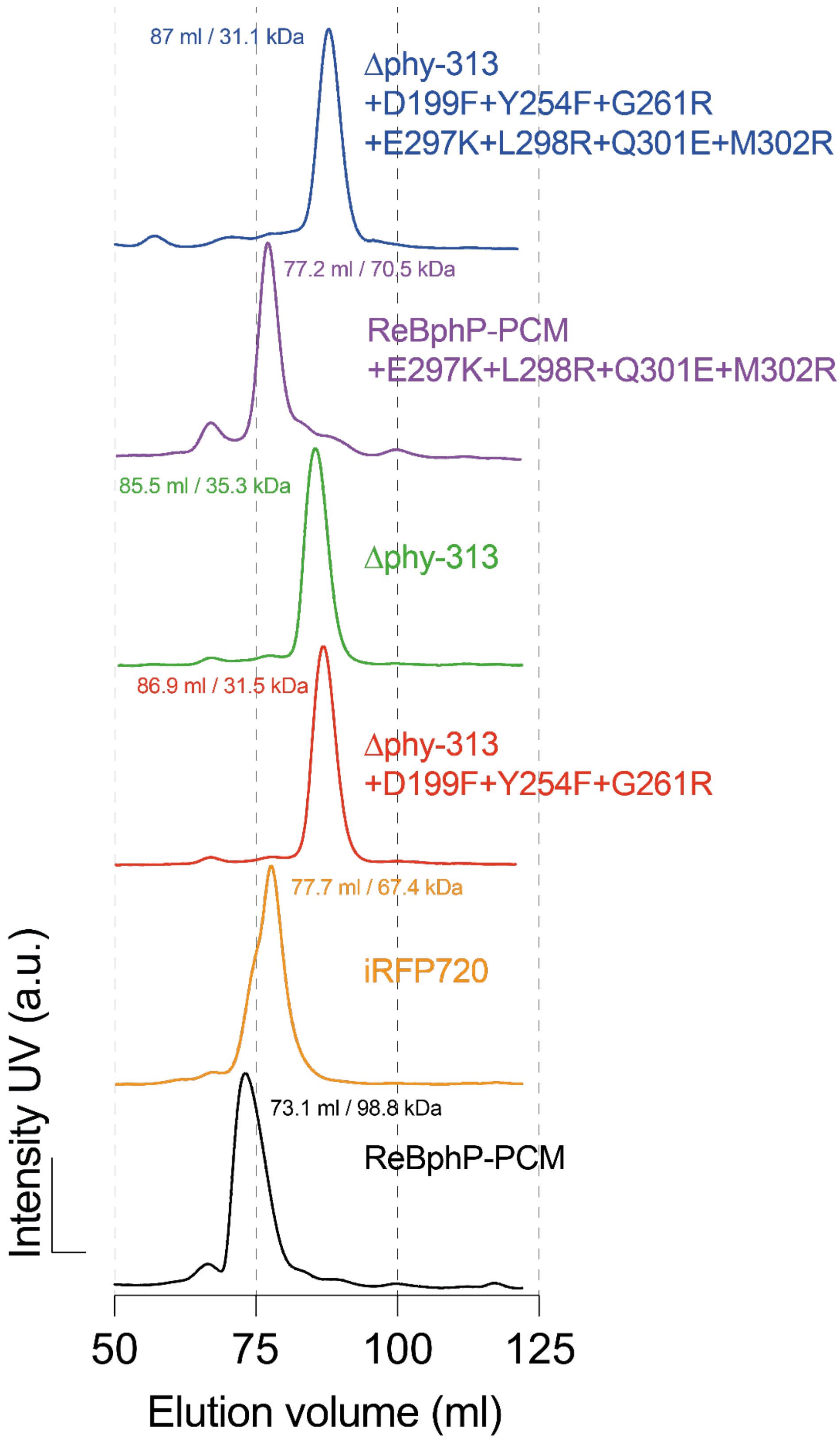


**Supplementary Figure 13:** Analytical size exclusion profiles of several variants discussed in this paper. 500 µl of 5 mg/ml mg/ml have been separated on a HiLoad 16/600 superdex 200pg column (GE Healthcare Life Sciences, Freiburg, Germany). Column elution profile was calibrated using a HMW Gel Filtration Calibration kit (Cytiva).

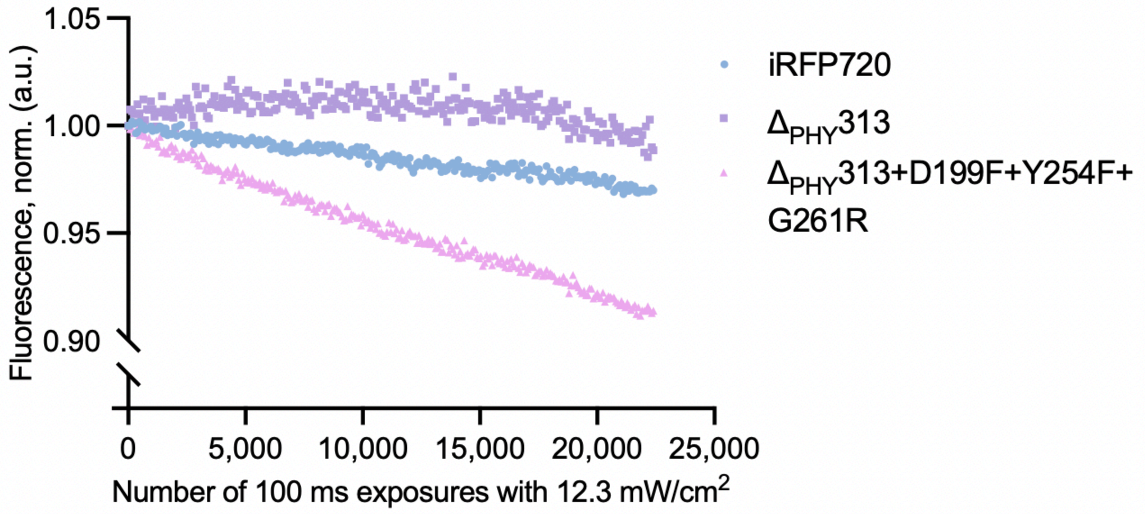


**Supplementary Figure 14:** Photofatigue measurements of variants $\Delta_{PHY}$313 and $\Delta_{PHY}$313+D199F+Y254F+G261R in comparison to IRFP720. Proteins have been illuminated with repeated exposure of 12.3 mW/cm$^2$ light. Shown is the linear part of the bleaching. All proteins show a level of first cycle drop of 2 %, 7 % and 1 %, for variants $\Delta_{PHY}$313 and $\Delta_{PHY}$313+D199F+Y254F+G261R and iRFP720, respectively.

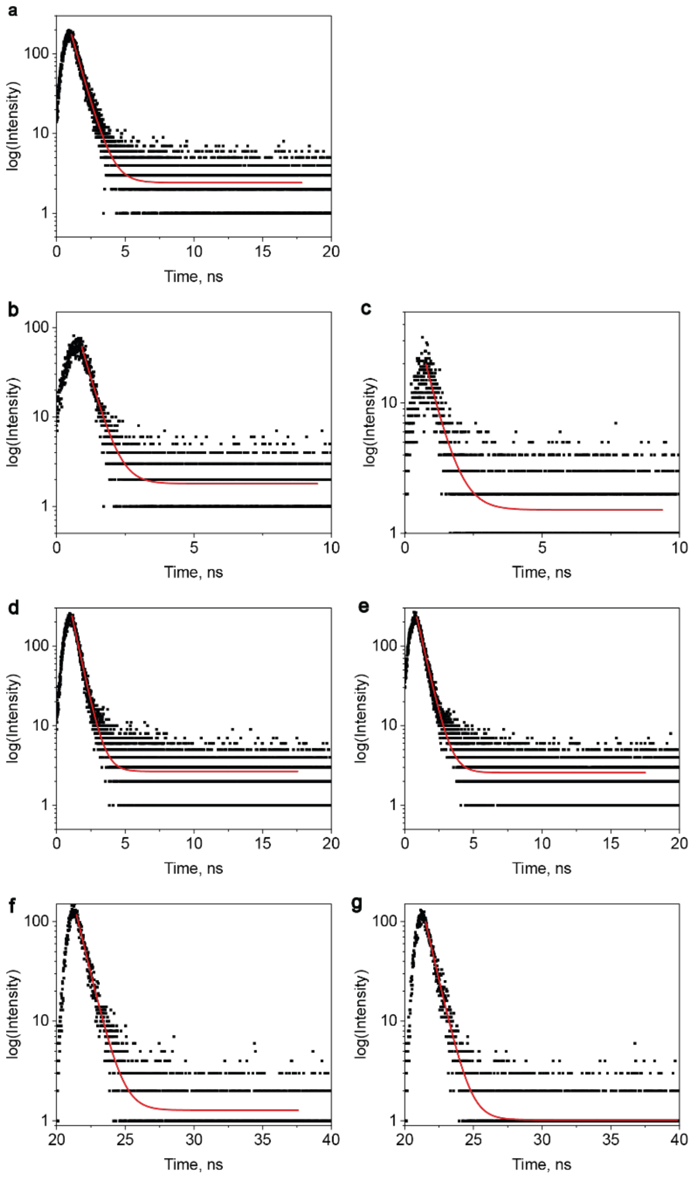


**Supplementary Figure 15:** Fluorescence lifetime measurements of iRFP720 (**a**, 710 ps, $\chi^2$_red=10.42, N=1), ReBphP $P_r$ (**b**, 390 ± 20 ps, $\chi^2$_red=4.62, N=2), ReBphP $P_{fr}$ (**c**, 490 ps, $\chi^2$_red=2.72, N=1, bad fit due to very low fluorescence)**,** Dphy-313 $P_r$ (**d**, 520 ± 20 ps, $\chi^2$_red=11.85, N=5), Dphy-313 $P_{fr}$ (**e**, 560 ± 30 ps, $\chi^2$_red=11.30, N=4), Dphy-313+D199F+Y254F+G261R $P_r$ (**f**, 750 ± 20 ps, $\chi^2$_red=7.22, N=3), Dphy-313+D199F+Y254F+G261R $P_{fr}$ (**g**, 730 ± 30 ps, $\chi^2$_red=4.11, N=3). The reported $\chi^2$_red corresponds to the curve displayed in the graph. All samples were excited at 375 nm.

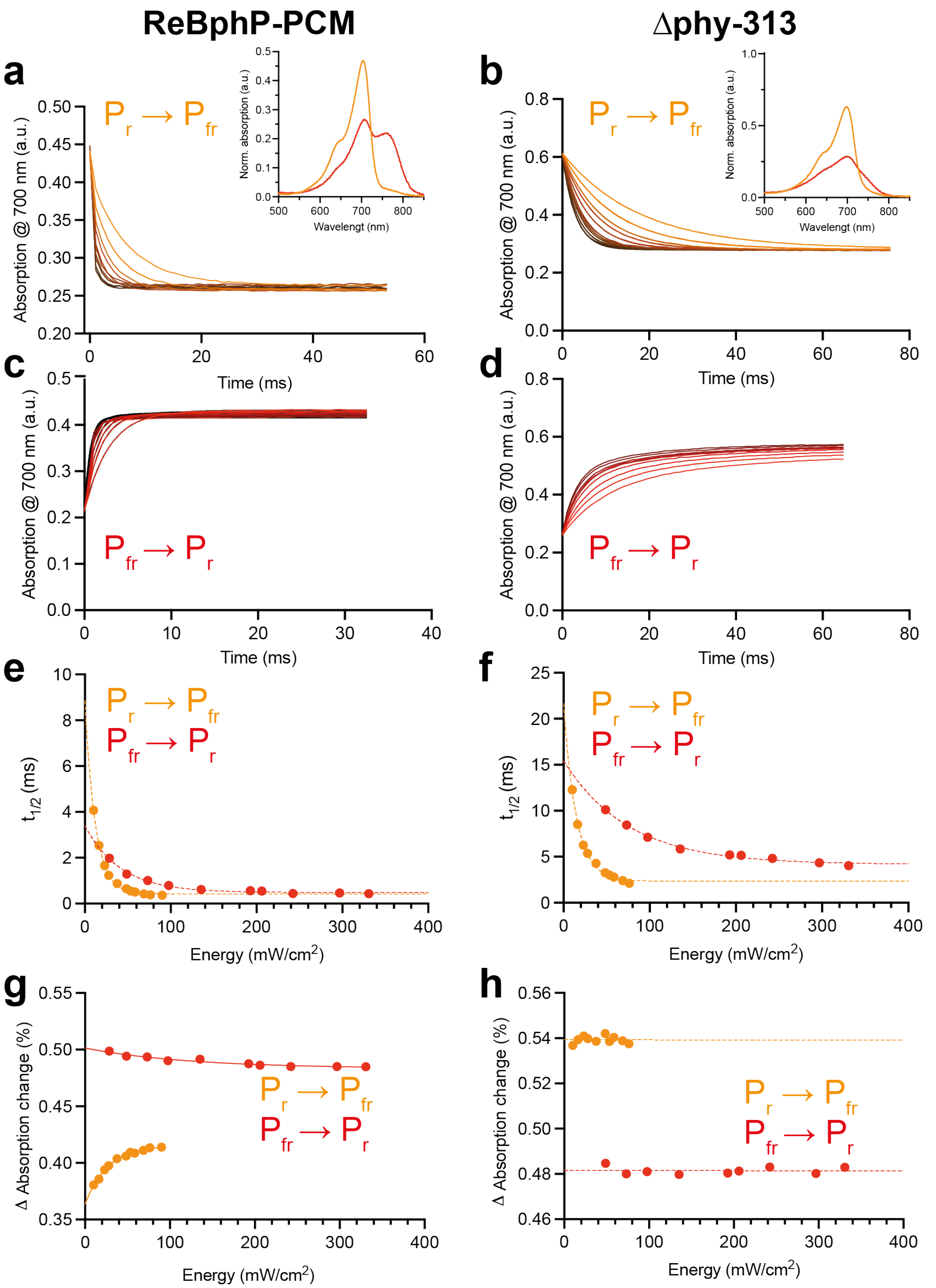


**Supplementary Figure 16:** Illumination energy dependence of photoswitching for ReBphP-PCM (**a**, **c**, **e** and **g**) and variant Δphy-313 (**b**, **d**, **f** and **h**). Shown are the $P_r$ to $P_{fr}$ (**a** and **b**) and the $P_{fr}$ to $P_r$ (**c** and **d**) transitions at different laser energies. Always the absorbance changes at 700 nm are monitored. $P_r$ and $P_{fr}$ end state spectra are shown as insets. Transitions are fit with a single exponential. The halftime (**e** and **f**) as well as the Δ of switching (**g** and **h**) in dependence of energy are shown. The dependence of halftime on energy could be fitted with an exponential in all cases while the dependence Δ of switching showed an exponential behavior only in the case of ReBphP but not the variant.

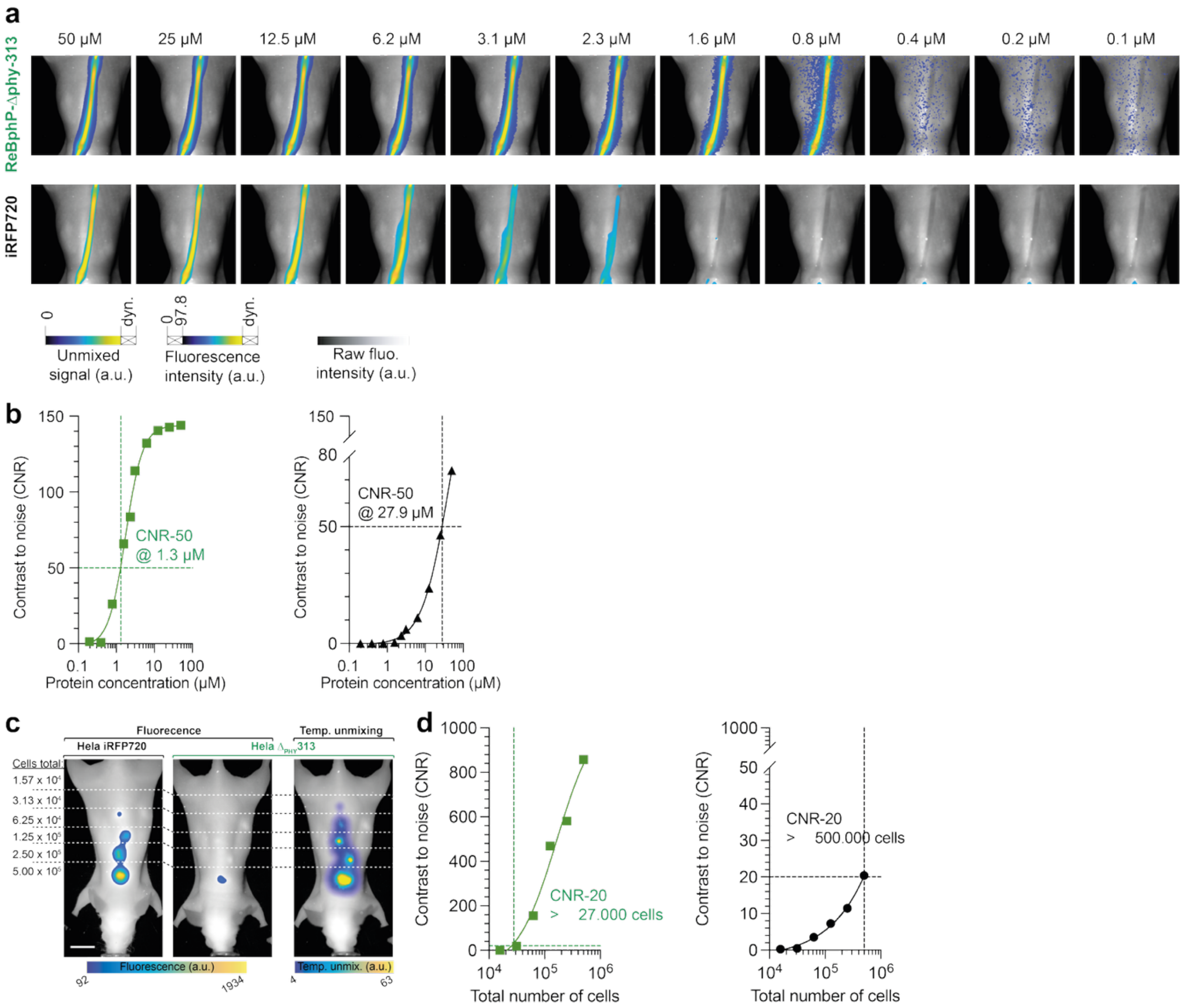


**Supplementary Figure 17:** Photoswitching driven NIR imaging of variant $\Delta_{PHY}313$ and iRFP720 using an 800 nm longpass filter for detection. Visualization as in Figure 4. In brief: **a** Ex vivo phantom with different concentrations of protein. **b** Fitting of the data shown in a. **c** In vivo imaging of Matrigel plugs with different cell numbers **d** Fitting of the data shown in c.

**Supplementary Data 1:** MSA of major engineered BphPs, their parent proteins and ReBphP-PCM. Wildtype proteins in bold, colors according to Figure 2a.

```
                                                    PAS
BrBphP          -----------MPVPLT------TP---AFGHATLANCEREQIHLAGSIQPHGILLAVKE    40
mIFP            -----------MSVPLT------TS---AFGHAFLANCEREQIHLAGSIQPHGILLAVKE    40
RpBphP1         ---------MVAGHASG------SP---AFGTADLSNCEREEIHLAGSIQPHGALLVVSE    42
miRFP           ---------MVAGHASG------SP---DFGTADPSDCEREEIHLAGSIQPHGTLLVVSE    42
miRFP2          ---------MVAGHASG------SP---DFGTASPSDCEHEEIHLAGSIQPHGTLLVVSE    42
miRFP709        ---------MVAGHASG------SP---AFGTASHSNCEHEEIHLAGSIQPHGALLVVSE    42
miRFP703        ---------MVAGHASG------SP---AFGTASHSNCEHEEIHLAGSIQPHGALLVVSE    42
miRFP670        ---------MVAGHASG------SP---AFGTASHSNCEHEEIHLAGSIQPHGALLVVSE    42
emiRFP703       ---------MAEGSV--------------ARQPDLLTCEHEEIHLAGSIQPHGALLVVSE    37
emiRFP670       ---------MAEGSV--------------ARQPDLLTCEHEEIHLAGSIQPHGALLVVSE    37
DrBphP-PCM      --------------MSRDPLPFFPPLYLGGPEITTENCEREPIHIPGSIQPHGALLTADG    46
WiPhy           MASMTGGQQMGRGSMSRDPLPFFPPLYLGGPEITTENCEREPIHIPGSIQPHGALLTADG    60
SNIFP           --------------MSRDPLPFFPPLYLGGPEITTENCEREPIHIPGSIQPHGALLTADG    46
iFP1.4          --------------MARDPLPFFPPLYLGGPEITTENCEREPIHIPGSIQPHGALLTADG    46
iFP2.0          --------------MARDPQPFFPPLYLGGPEITTENCEREPIHIPGSIQPHGALLTADG    46
ReBphP-PCM      -----------MSGTRE--------------PMDLTNCDREPIHQLGAIQPFGFLLQVSS    35
RpBphP6         --------------MPR--------------KVDLTSCDREPIHIPGSIQPCGCLLACDA    32
iRFP702         --------------MAR--------------KVDLTSCDREPIHIPGSIQPCGCLLACDA    32
iRFP670         --------------MAR--------------KVDLTSCDREPIHIPGSIQPCGCLLACDA    32
miRFP702        --------------MAR--------------KVDLTSCDREPIHIPGSIQPCGCLLACDA    32
iRFP670-2       --------------MAR--------------KVDLTSCDREPIHIPGSIQPCGCLLACDA    32
Agp1            -------------MSSH--------------TPKLDSCGAEPIHIPGAIQEHGALLVLSA    33
RpBphP2         ---------MTEGSVAR--------------QPDLSTCDDEPIHIPGAIQPHGLLLALAA    37
miRFP682        ---------MAEGSVAR--------------QPDLLTCDDEPIHIPGAIQPHGLLLALAA    37
miRFP720        ---------MAEGSVAR--------------QPDLLTCDDEPIHIPGAIQPHGLLLALAA    37
miRFP713        ---------MAEGSVAR--------------QPDLLTCDDEPIHIPGAIQPHGLLLALAA    37
miRFP680        ---------MAEGSVAR--------------QPDLLTCDDEPIHIPGAIQPHGLLLALAA    37
mRhubarb720     ---------MAEGSVAR--------------QPDLLTCDDEPIHIPGAIQPHGLLLALAA    37
mRhubarb719     ---------MAEGSVAR--------------QPDLLTCDDEPIHIPGAIQPHGLLLALAA    37
mRhubarb713     ---------MAEGSVAR--------------QPDLLTCDDEPIHIPGAIQPHGLLLALAA    37
iRFP720         ---------MAEGSVAR--------------QPDLLTCDDEPIHIPGAIQPHGLLLALAA    37
iRFP682         ---------MAEGSVAR--------------QPDLLTCDDEPIHIPGAIQPHGLLLALAA    37
iRFP713         ---------MAEGSVAR--------------QPDLLTCDDEPIHIPGAIQPHGLLLALAA    37
iRFP713/V256C   ---------MAEGSVAR--------------QPDLLTCDDEPIHIPGAIQPHGLLLALAA    37

                                                    PAS
BrBphP          PDNVVIQASINAAEFLNVSSVVGRPLRDLGGDLA----LQILPHL---NGPLHLAPMTLR    93
mIFP            PDNVVIQASINAAEFLNTNSVVGRPLRDLGGDLP----LQILPHL---NGPLHLAPMTLR    93
RpBphP1         PDHRIIQASANAAEFLNLGSVLGVPLAEIDGDLL----IKILPHL---DPTAEGMPVAVR    95
miRFP           PDHRIIQASANAAEFLNLGSVLGVPLAEIDGDLL----IKILPHL---DPTAEGMPVAVR    95
miRFP2          PDHRIIQASANAAEFLNLGSVLGIPLAEIDGDLL----IKILPHL---DLTAEGMPVAVR    95
miRFP709        HDHRVIQASANAAEFLNLGSVLGVPLAEIDGDLL----IKILPHL---DPTAEGMPVAVR    95
miRFP703        HDHRVIQASANAAEFLNLGSVLGVPLAEIDGDLL----IKILPHL---DPTAEGMPVAVR    95
miRFP670        HDHRVIQASANAAEFLNLGSVLGVPLAEIDGDLL----IKILPHL---DPTAEGMPVAVR    95
emiRFP703       HDHRVIQASANAAEFLNLGSVLGVPLAEIDGDLL----IKILPHL---DPTAEGMPVAVR    90
emiRFP670       HDHRVIQASANAAEFLNLGSVLGVPLAEIDGDLL----IKILPHL---DPTAEGMPVAVR    90
DrBphP-PCM      HSGEVLQMSLNAATFLGQEPT--VLRGQTLAALL-PEQWPALQAALP-PGCP----DALQ    98
WiPhy           HSGEVLQMSLNAATFLGQEPT--VLRGQTLAALL-PEQWPALQAALP-PGCP----DALQ    112
SNIFP           HSGEVLQMSLNAATFLGQEPT--VLRGQTLAALL-PEQWPALQAALP-PGCP----DALQ    98
iFP1.4          HSGEVLQVSLNAATFLGQEPT--VLRGQTLAALL-PEQWPALQAALP-PGCP----DALQ    98
iFP2.0          HSGEVLQVSLNAATFLGQEPT--VLRGQTLAALL-PDQWPALQTALP-PGCQ----DALQ    98
ReBphP-PCM      DWI-VTRASVNLAEFLGVTQA--DALGRPASTLIMPEALHTIRNKLTTLRGADVVERVFA    92
RpBphP6         QAVRITRISENAGAFFGRETP--R-VGELLADYFGETEAHALRNALAQSSDPKRPALIFG    89
iRFP702         QAVRITRITENAGAFFGRETP--R-VGELLADYFGETEAHALRNALAQSSDPKRPALIFG    89
iRFP670         QAVRITRITENAGAFFGRETP--R-VGELLADYFGETEAHALRNALAQSSDPKRPALIFG    89
miRFP702        QAVRITRITENAGAFFGRETP--R-VGELLADYFGETEAHALRNALAQSSDPKRPALIFG    89
iRFP670-2       QAVRITRITENAGAFFGRETP--R-VGELLADYFGETEAHALRNALAQSSDPKRPALIFG    89
Agp1            REFSVVQASDNLANYIGVDLPIGAVATEANLPFI---------------------SVLSA    72
RpBphP2         -DMTIVAGSDNLPELTGLAIG--ALIGRSAADVFDSETHNRLTIALAEPGAAVGAPIAVG    94
miRFP682        -DMTIVAGSDNLPELTGLAIG--ALIGRSAADVFDSETHNRLTIALAEPGAAVGAPITVG    94
miRFP720        -DMTIVAGSDNLPELTGLAIG--ALIGRSAADVFDSETHNRLTIALAEPGAAVGAPITVG    94
miRFP713        -DMTIVAGSDNLPELTGLAIG--ALIGRSAADVFDSETHNRLTIALAEPGAAVGAPITVG    94
miRFP680        -DMTIVAGSDNLPELTGLAIG--ALIGRSAADVFDSETHNRLTIALAEPGAAVGAPITVG    94
mRhubarb720     -DMTIVAGSDNLPELTGLAIG--ALIGRSAADVFDSETHNRLTIALAEPGAAVGAPITVG    94
mRhubarb719     -DMTIVAGSDNLPELTGLAIG--ALIGRSAADVFDSETHNRLTIALAEPGAAVGAPITVG    94
mRhubarb713     -DMTIVAGSDNLPELTGLAIG--ALIGRSAADVFDSETHNRLTIALAEPGAAVGAPITVG    94
iRFP720         -DMTIVAGSDNLPELTGLAIG--ALIGRSAADVFDSETHNRLTIALAEPGAAVGAPITVG    94
iRFP682         -DMTIVAGSDNLPELTGLAIG--ALIGRSAADVFDSETHNRLTIALAEPGAAVGAPITVG    94
iRFP713         -DMTIVAGSDNLPELTGLAIG--ALIGRSAADVFDSETHNRLTIALAEPGAAVGAPITVG    94
iRFP713/V256C   -DMTIVAGSDNLPELTGLAIG--ALIGRSAADVFDSETHNRLTIALAEPGAAVGAPITVG    94

                                                    PAS        <-> GAF
BrBphP          CTAG--------SPPRRVDCTVHRPSNGGLIVELEPATKTTN----VAPALDGAFHRITS    141
mIFP            CTVG--------SPPRRVDCTIHRPSNGGLIVELEPATKTTN----IAPALDGAFHRITS    141
RpBphP1         CRIG--------NPSTEYDGLMHRPPEGGLIIELERAGPPID----LSGTLAPALERIRT    143
miRFP           CRIG--------NPSTEYDGLMHRPPEGGLIIELERAGPPID----LSGTLAPALERIRT    143
miRFP2          CRIG--------NPSTEYDGLMHRPPEGGLIIELERAGPSID----LSGTLAPALERIRT    143
miRFP709        CRIG--------NPSTEYCGLMHRPPEGGLIIELERAGPSID----LSGTLAPALERIRT    143
miRFP703        CRIG--------NPSTEYCGLMHRPPEGGLIIELERAGPSID----LSGTLAPALERIRT    143
miRFP670        CRIG--------NPSTEYCGLMHRPPEGGLIIELERAGPSID----LSGTLAPALERIRT    143
emiRFP703       CRIG--------NPSTEYCGLMHRPPEGGLIIELERAGPSID----LSGTLAPALERIRT    138
emiRFP670       CRIG--------NPSTEYCGLMHRPPEGGLIIELERAGPSID----LSGTLAPALERIRT    138
DrBphP-PCM      YRA-----TLDWPAAGHLSLTVHR-VGELLILEFEPTEAWDSTGP---HALRNAMFALES    149
WiPhy           YRA-----TLDWPAAGHLSLTVHR-VGELLILEFEPTEAWDSTGP---HALRNAMSALES    163
SNIFP           YRA-----TLDWPAAGHLSLTVHR-VGELLILEFEPTEAWDSTGP---HALRNAMSALES    149
iFP1.4          YRA-----TLDWPAAGHLSLTVHR-VAELLILEFEPTEAWDSIGP---HALRNAMFALES    149
iFP2.0          YRA-----TLDWPAAGHLSLTVHR-VAELLILEFEPTEAWDSIGP---HALRNAMFALES    149
ReBphP-PCM      IA--------LTPDQSKFDVAVHF-NESQVIIEGERCQEDRRNA--PSLSMRSMMSRLDQ    141
RpBphP6         WR--------DGLTGRTFDISLHR-HDGTSIVEFEPAAADQADN--PLRLTRQIIARTKE    138
iRFP702         WR--------DGLTGRTFDISLHR-HDGTSIVEFEPAAAEQADN--PLRLTRQIIARTKE    138
iRFP670         WR--------DGLTGRTFDISLHR-HDGTSIIEFEPAAAEQADN--PLRLTRQIIARTKE    138
miRFP702        WR--------DGLTGRTFDISLHR-HDGTSIVEFEPAAAEQADN--PLRLTRQIIARTKE    138
iRFP670-2       WR--------DGLTGRTFDISLHR-HDGTSIIEFEPAAAEQADN--PLRLTRQIIARTKE    138
Agp1            WYSGEESNFRYAWAEKKLDVSAHR-SGTLVILEVEKAGVGES-AEKLMGELTSLAKYLNS    130
RpBphP2         FTMRKDAGFVGSW---------HR-HDQLVFLELEPPQRDVAEPQAFFRRTNSAIRRLQA    144
miRFP682        FTMRKDAGFIGSW---------HR-HDQLIFLELEPPQRDVAEPQAFFRRTNSAIRRLQA    144
miRFP720        FTMRKDAGFIGSW---------HR-HDQLIFLELEPPQRDVAEPQAFFRRTNSAIRRLQA    144
miRFP713        FTMRKDAGFIGSW---------HR-HDQLIFLELEPPQRDVAEPQAFFRRTNSAIRRLQA    144
miRFP680        FTMRKDAGFIGSW---------HR-HDQLIFLELEPPQRDVAEPQAFFRRTNSAIRRLQA    144
mRhubarb720     FTMRKDAGFIGSW---------HR-HDQLIFLELEPPQRDVAEPQAFFRHTNSAIRRLQA    144
mRhubarb719     FTMRKDAGFIGSW---------HR-HDQLIFLELEPPQRDVAEPQAFFRHTNSAIRRLQA    144
mRhubarb713     FTMRKDAGFIGSW---------HR-HDQLIFLELEPPQRDVAEPQAFFRHTNSAIRRLQA    144
iRFP720         FTMRKDAGFIGSW---------HR-HDQLIFLELEPPQRDVAEPQAFFRRTNSAIRRLQA    144
iRFP682         FTMRKDAGFIGSW---------HR-HDQLIFLELEPPQRDVAEPQAFFRRTNSAIRRLQA    144
iRFP713         FTMRKDAGFIGSW---------HR-HDQLIFLELEPPQRDVAEPQAFFRRTNSAIRRLQA    144
iRFP713/V256C   FTMRKDAGFIGSW---------HR-HDQLIFLELEPPQRDVAEPQAFFRRTNSAIRRLQA    144
```

```
                                      GAF
BrBphP            SSSLIGLCDETATIFREITGYDRVMVYRFDEEGHGEVLSERRRPDLEAFLGNRYPASDIP    201
mIFP              SSSLMGLCDETATIIREITGYDRVMVVRFDEEGNGEILSERRRADLEAFLGNRYPASTIP    201
RpBphP1           AGSLRALCDDTALLFQQCTGYDRVMVYRFDEQGHGEVFSERHVPGLESYFGNRYPSSDIP    203
miRFP             AGSLRALCDDTALLFQQCTGYDRVMVYRFDEQGHGEVYSEIHVTGLESYFGNRYPSSLVP    203
miRFP2            AGSLRALCDDTVLLFQQCTGYDRVMVYRFDERGHGEVYSEIHVTGLESYFGNRYPSSLVP    203
miRFP709          AGSLRALCDDTVLLFQQCTGYDRVMVYRFDEQGHGLVFSECHVPGLESYFGNRYPSSFIP    203
miRFP703          AGSLRALCDDTVLLFQQCTGYDRVMVYRFDEQGHGLVFSECHVPGLESYFGNRYPSSLVP    203
miRFP670          AGSLRALCDDTVLLFQQCTGYDRVMVYRFDEQGHGLVFSECHVPGLESYFGNRYPSSTVP    203
emiRFP703         AGSLRALCDDTVLLFQQCTGYDRVMVYRFDEQGHGLVFSECHVPGLESYFGNRYPSSLVP    198
emiRFP670         AGSLRALCDDTVLLFQQCTGYDRVMVYRFDEQGHGLVFSECHVPGLESYFGNRYPSSTVP    198
DrBphP-PCM        APNLRALAEVATQTVRELTGFDRVMLYKFAPDATGEVIAEARREGLHAFLGHRFPASDIP    209
WiPhy             APNLRALAEVATQTVRELTGFDRVMLYKFAPDATGEVIAEARREGLHAFLGHRFPASHIP    223
SNIFP             APNLRALAEVATQTVRELTGFDRVMLYKFAPDATGEVIAEARREGLHAFLGHRFPASLIP    209
iFP1.4            APNLRALAEVATQTVRELTGFDRVMLYKFAPDATGEMIAEARREGMQAFLGHRFPASHTP    209
iFP2.0            APNLRALAEVATQTVRELSGFDRVMLYKFAPDATGEVIAEARREGMQAYLGHRFPASTTP    209
ReBphP-PCM        AETLEAFFREGARQARALTGFDRVMVYRFDEGGSGEVVAEAARSGIGSFLGLHYPASDIP    201
RpBphP6           LKSLEEMAARVPRYLQAMLGYHRVMMYRFADDGSGKVIGEAKRSDLESFLGQHFPASDIP    198
iRFP702           LKSLEEMAARVPRYLQAMLGYHRVMLYRFADDGSGKVIGEAKRSDLESFLGQHFPASLVP    198
iRFP670           LKSLEEMAARVPRYLQAMLGYHRVMLYRFADDGSGMVIGEAKRSDLESFLGQHFPASLVP    198
miRFP702          LKSLEEMAARVPRYLQAMLGYHRVMLYRFADDGSGKVIGEAKRSDLESFLGQHFPASLVP    198
iRFP670-2         LKSLEEMAARVPRYLQAMLGYHRVMLYRFADDGSGMVIGEAKRSDLESFLGQHFPASLVP    198
Agp1              APSLEDALFRTAQLVSSISGHDRTLIYDFGLDWSGHVVAEAGSGALPSYLGLRFPAGDIP    190
RpBphP2           AETLESACAAAAQEVREITGFDRVMIYRFASDFSGEVIAEDRCAEVESYLGLHFPASDIP    204
miRFP682          AETLESACAAAAQEVRKITGFDRVMIYRFASDFSGVVIAEDRCAEVESKLGLHYPASAVP    204
miRFP720          AETLESACAAAAQEVRKITGFDRVMIYRFASDFSGEVIAEDRCAEVESKLGLHYPASFIP    204
miRFP713          AETLESACAAAAQEVRKITGFDRVMIYRFASDFSGEVIAEDRCAEVESKLGLHYPASTVP    204
miRFP680          AETLESACAAAAQEVRKITGFDRVMIYRFASDFSGEVIAEDRCAEVESKLGLHYPASTVP    204
mRhubarb720       AETLESACAAAAQEVRKITGFDRVMIYRFASDFSGEVIAEDRCAEVESKLGQHYPASDIP    204
mRhubarb719       AETLESACAAAAQEVRKITGFDRVMIYRFASDFSGEVIAEDRCAEVESKLGLHSPASYIP    204
mRhubarb713       AETLESACAAAAQEVRKITGFDRVMIYRFASDFSGEVIAEDRCAEVESKLGLHYPASTVP    204
iRFP720           AETLESACAAAAQEVRKITGFDRVMIYRFASDFSGSVIAEDRCAEVESKLGLHYPASFIP    204
iRFP682           AETLESACAAAAQEVRKITGFDRVMIYRFASDFSGVVIAEDRCAEVESKLGLHYPASAVP    204
iRFP713           AETLESACAAAAQEVRKITGFDRVMIYRFASDFSGEVIAEDRCAEVESKLGLHYPASTVP    204
iRFP713/V256C     AETLESACAAAAQEVRKITGFDRVMIYRFASDFSGEVIAEDRCAEVESKLGLHYPASTVP    204

                                      GAF
BrBphP            QIARRLYERNRVRLLVDVNYSPVPLQPRISPLNGRDLDMSLSCLRSMSPIHQKYLQNMGV    261
mIFP              QIARRLYEHNRVRLLVDVNYTPVPLQPRISPLNGRDLDMSLSCLRSMSPIHQKYMQDMGV    261
RpBphP1           QMARRLYERQRVRVLVDVSYQPVPLEPRLSPLTGRDLDMSGCFLRSMSPIHLQYLKNMGV    263
miRFP             QMARRLYERQRVRVLVDVSYQPVPLEPRLSPLTGRDLDMSGCFLRSMSPTHLQFLKNMGV    263
miRFP2            QMARRLYVRQRVRVLVDVTYQPVPLEPRLSPLTGRDLDMSGCFLRSMSPTHLQFLKNMGV    263
miRFP709          QMARQLYVRQRVRVLVDVTYQPVPLEPRLSPLTGRDLDMSGCFLRSMSPIHLQFLKDMGV    263
miRFP703          QMARQLYVRQRVRVLVDVTYQPVPLEPRLSPLTGRDLDMSGCFLRSMSPIHLQFLKDMGV    263
miRFP670          QMARQLYVRQRVRVLVDVTYQPVPLEPRLSPLTGRDLDMSGCFLRSMSPCHLQFLKDMGV    263
emiRFP703         QMARQLYVRQRVRVLVDVTYQPVPLEPRLSPLTGRDLDMSGCFLRSMSPIHLQFLKDMGV    258
emiRFP670         QMARQLYVRQRVRVLVDVTYQPVPLEPRLSPLTGRDLDMSGCFLRSMSPCHLQFLKDMGV    258
DrBphP-PCM        AQARALYTRHLLRLTADTRAAAVPLDPVLNPQTNAPTPLGGAVLRATSPMHMQYLRNMGV    269
WiPhy             AQARALYTRHLLRLTADTRAAAVPLDPVLNPQTNAPTPLGGAVLRATSPMHMQFLRNMGV    283
SNIFP             AQARALYTRHLLRLTADTRAAAVPLDPVLNPQTNAPTPLGGAVLRATSPMHMQFLRNMGV    269
iFP1.4            AQARALYTRHLLRLTADTRAAAVPLDPVLNPQTNAPTPLGGAVLRATSPMHMQYLRNMGV    269
iFP2.0            AQARALYTRHLLRLTADTRAAAVPLDPVLNPQTNAPTPLGGAVLRATSPMHMQYLRNMGV    269
ReBphP-PCM        VQARALYLRNLFRIIADVDAVPVPILPERD-EHGQPLDLSMSVLRSVSPIHIEYLKNMGV    260
RpBphP6           QQARLLYLKNAIRVISDSRGISSRIVPERD-ASGAALDLSFAHLRSVSPIHLEYLRNMGV    257
iRFP702           QQARLLYLKNAIRVVSDSRGISSRIVPEHD-ASGAALDLSFAHLRSISPIHLEFLRNMGV    257
iRFP670           QQARLLYLKNAIRVVSDSRGISSRIVPEHD-ASGAALDLSFAHLRSISPCHLEFLRNMGV    257
miRFP702          QQARLLYLKNAIRVVSDSRGISSRIVPEHD-ASGAALDLSFAHLRSISPIHLEFLRNMGV    257
iRFP670-2         QQARLLYLKNAIRVVSDSRGISSRIVPEHD-ASGAALDLSFAHLRSISPCHLEFLRNMGV    257
Agp1              PQARQLYTINRLRMIPDVDYKPVPIRPEVNAETGAVLDMSFSQLRSVSPVHLEYMRNMGT    250
RpBphP2           AQARRLYTINPVRIIPDINYRPVPVTPDLNPVTGRPIDLSFAILRSVSPVHLEYMRNIGM    264
miRFP682          AQARRLYTINPVRIIPDINYRPVPVTPDLNPVTGRPIDLSFAILRSVSPCHLEFMRNIGM    264
miRFP720          AQARRLYTINPVRIIPDINYRPVPVTPDLNPVTGRPIDLSFAILRSVSPVHLEFMRNIGM    264
miRFP713          AQARRLYTINPVRIIPDINYRPVPVTPDLNPVTGRPIDLSFAILRSVSPVHLEFMRNIGM    264
miRFP680          AQARRLYTINPVRIIPDINYRPVPVTPDLNPVTGRPIDLSFAILRSVSPCHLEFMRNIGM    264
mRhubarb720       AQARRLYTINPVRIIPDINYRPVPVTPDLNPVTGRPIDLSFAILRSVSPVHLEFMRNIGM    264
mRhubarb719       AQARRLYTINPVRIIPDINYRPVPVTPDLNPVTGRPIDLSFAILRSVSPVHLEFMRNIGM    264
mRhubarb713       AQARRLYTINPVRIIPDINYRPVPVTPDLNPVTGRPIDLSFAILRSVSPVHLEFMRNIGM    264
iRFP720           AQARRLYTINPVRIIPDINYRPVPVTPDLNPVTGRPIDLSFAILRSVSPNHLEFMRNIGM    264
iRFP682           AQARRLYTINPVRIIPDINYRPVPVTPDLNPVTGRPIDLSFAILRSVSPCHLEFMRNIGM    264
iRFP713           AQARRLYTINPVRIIPDINYRPVPVTPDLNPVTGRPIDLSFAILRSVSPVHLEFMRNIGM    264
iRFP713/V256C     AQARRLYTINPVRIIPDINYRPVPVTPDLNPVTGRPIDLSFAILRSVSPCHLEFMRNIGM    264

                                      GAF                          ←→ PHY
BrBphP            GATLVCSLMVSGRLWGLIACHHYEPRFVPFDIRAAGEALAETCAIRIAALESFAQSQSEL    321
mIFP              GATLVCSLMVSGRLWGLIACHHYEPRFVPFHIRAAGEALAETCAIRIATLESFAQSQSK-    320
RpBphP1           RATLVVSLVVGGKLWGLVACHHYLPRFIHFELRAICELLAEAIATRITALES--------    315
miRFP             RATLVVSLVVGGKLWGLVICHHYLPRFIHFELRAICELLAEAIATRITAL----------    313
miRFP2            RATLVVSLVVGGKLWGLVICHHYLPRFIHFELRAICVLLAEAIATRITAL----------    313
miRFP709          RATLAVSLVVGGKLWGLVVCHHYLPRFIRFELRAICKRLAERIATRITALES--------    315
miRFP703          RATLAVSLVVGGKLWGLVVCHHYLPRFIRFELRAICKRLAERIATRITALES--------    315
miRFP670          RATLAVSLVVGGKLWGLVVCHHYLPRFIRFELRAICKRLAERIATRITALES--------    315
emiRFP703         RATLAVSLVVGGKLWGLVVCHHYLPRFIRFELRAICKRLAERIATRITALES--------    310
emiRFP670         RATLAVSLVVGGKLWGLVVCHHYLPRFIRFELRAICKRLAERIATRITALES--------    310
DrBphP-PCM        GSSLSVSVVVGGQLWGLIACHHQTPYVLPPDLRTTLEYLGRLLSLQVQVKEAADVAAFRQ    329
WiPhy             GSSLSVSVVVGGQLWGLIACHHQTPYVLPPDLRTTLEYLGRELSEQVQVKEA--------    335
SNIFP             RSSLSVSVVVGGQLWGLIACHHQTPYVLPPDLRTTLEYLGRELSEQVQVKE---------    320
iFP1.4            GSSLSVSVVVGGQLWGLIVCHHQTPYVLPPDLRTTLEELGRKLSGQVQRKEA--------    321
iFP2.0            GSSLSVSVVVGGQLWGLIVCHHQTPYVLPPDLRTTLEYLGRLLSLQVQRKEA--------    321
ReBphP-PCM        GASLSISIVVDGKLWGLFACHHYEPRLPSAQSRSTAELFGQMFASRLESRERRLALDYET    320
RpBphP6           SASMSLSIIIDGTLWGLIACHHYEPRAVPMAQRVAAEMFADFFSLHFTAAHHQR------    311
iRFP702           SASMSLSIIIDGTLWGLIICHHYEPRAVPMAQRVAAEMFADFLSLHFTAAHHQR------    311
iRFP670           SASMSLSIIIDGTLWGLIICHHYEPRAVPMAQRVAAEMFADFLSLHFTAAHHQR------    311
miRFP702          SASMSLSIIIDGTLWGLIICHHYEPRAVPMAQRVAAKRFAERLSTHFTAAHHQR------    311
iRFP670-2         SASMSLSIIIDGTLWGLIICHHYEPRAVPMAQRVAAKRFAERLSTHFTAAHHQR------    311
Agp1              AASMSVSIVVNGALWGLIACHHATPHSVSLAVREACDFAAQLLSMRIAMEQSSQDASRRV    310
RpBphP2           HGTMSISILRGERLWGLIACHHRKPNYVDLDGRQACELVAQVLAWQIGVMEE--------    316
miRFP682          HGTMSISILRGERLWGLIVCHHRTPYYVDLDGRQACKRVAERLATQIGVMEE--------    316
miRFP720          HGTMSISILRGERLWGLIVCHHRTPYYVDLDGRQACKRVAERLATQIGVMEE--------    316
miRFP713          HGTMSISILRGERLWGLIVCHHRTPYYVDLDGRQACKRVAERLATQIGVMEE--------    316
miRFP680          HGTMSISILRGERLWGLIVCHHRTPYYVDLDGRQACKRVAERLATQIGVMEE--------    316
mRhubarb720       HGTMSISILRGERLWGLIVCHHRTPYYVDLDGRQACELVAQVLARAIGVMEE--------    316
mRhubarb719       HGTMSISILRGERLWGLIVCHHRTPYYVDLDGRQACELVAQVLARAIGVMEE--------    316
mRhubarb713       HGTMSISILRGERLWGLIVCHHRTPYYVDLDGRQACELVAQVLARAIGVMEE--------    316
iRFP720           HGTMSISILRGERLWGLIVCHHRTPYYVDLDGRQACELVAQVLAWQIGVMEE--------    316
iRFP682           HGTMSISILRGERLWGLIVCHHRTPYYVDLDGRQACELVAQVLAWQIGVMEE--------    316
iRFP713           HGTMSISILRGERLWGLIVCHHRTPYYVDLDGRQACELVAQVLAWQIGVMEE--------    316
iRFP713/V256C     HGTMSISILRGERLWGLIVCHHRTPYYVDLDGRQACELVAQVLAWQIGVMEE--------    316
```

```
                                       PHY
BrBphP             VVRRLEQRLVEAVSRDGEWQAALFDGS---QSLLQPLGASGAALLFEGQVMTAGDVPSTQ    378
mIFP               ------------------------------------------------------------    320
RpBphP1            ------------------------------------------------------------    315
miRFP              ------------------------------------------------------------    313
miRFP2             ------------------------------------------------------------    313
miRFP709           ------------------------------------------------------------    315
miRFP703           ------------------------------------------------------------    315
miRFP670           ------------------------------------------------------------    315
emiRFP703          ------------------------------------------------------------    310
emiRFP670          ------------------------------------------------------------    310
DrBphP-PCM         SLREHHARVALAAAHSLSPHDTL---SDPALDLLGLMRAGGLILRFEGRWQTLGEVPPAP    386
WiPhy              ------------------------------------------------------------    335
SNIFP              ------------------------------------------------------------    320
iFP1.4             ------------------------------------------------------------    321
iFP2.0             ------------------------------------------------------------    321
ReBphP-PCM         KARRIADRLLTSVADNASLLDDP---AWLIEALADAIPADGIGVWINGRLALAGIGPDKK    377
RpBphP6            ------------------------------------------------------------    311
iRFP702            ------------------------------------------------------------    311
iRFP670            ------------------------------------------------------------    311
miRFP702           ------------------------------------------------------------    311
iRFP670-2          ------------------------------------------------------------    311
Agp1               ELGHIQARLLKGMAAAEKWVDGLLGGEGEREDLLKQVGADGAALVLGDDYELVGNTPSRE    370
RpBphP2            ------------------------------------------------------------    316
miRFP682           ------------------------------------------------------------    316
miRFP720           ------------------------------------------------------------    316
miRFP713           ------------------------------------------------------------    316
miRFP680           ------------------------------------------------------------    316
mRhubarb720        ------------------------------------------------------------    316
mRhubarb719        ------------------------------------------------------------    316
mRhubarb713        ------------------------------------------------------------    316
iRFP720            ------------------------------------------------------------    316
iRFP682            ------------------------------------------------------------    316
iRFP713            ------------------------------------------------------------    316
iRFP713/V256C      ------------------------------------------------------------    316

                                       PHY
BrBphP             RIRDLASWLDRRRKAPDAPAQGLTVTASLTLDEPSFADIRSIASGLIAAPLSASEGEYLL    438
mIFP               ------------------------------------------------------------    320
RpBphP1            ------------------------------------------------------------    315
miRFP              ------------------------------------------------------------    313
miRFP2             ------------------------------------------------------------    313
miRFP709           ------------------------------------------------------------    315
miRFP703           ------------------------------------------------------------    315
miRFP670           ------------------------------------------------------------    315
emiRFP703          ------------------------------------------------------------    310
emiRFP670          ------------------------------------------------------------    310
DrBphP-PCM         AVDALLAWLETQPG-------ALVQTDALGQLWPAGADLAPSAAGLLAISVGEGWSECLV    439
WiPhy              ------------------------------------------------------------    335
SNIFP              ------------------------------------------------------------    320
iFP1.4             ------------------------------------------------------------    321
iFP2.0             ------------------------------------------------------------    321
ReBphP-PCM         NFASLVRHLNRNAGG------RIYAVDRLSQTYPDLEI-DAVVAGMLAIPISRSPRDYVV    430
RpBphP6            ------------------------------------------------------------    311
iRFP702            ------------------------------------------------------------    311
iRFP670            ------------------------------------------------------------    311
miRFP702           ------------------------------------------------------------    311
iRFP670-2          ------------------------------------------------------------    311
Agp1               QVEELILWLGEREIA------DVFATDNLAGNYPTAAAYASVASGIIAMRVSELHGSWLI    424
RpBphP2            ------------------------------------------------------------    316
miRFP682           ------------------------------------------------------------    316
miRFP720           ------------------------------------------------------------    316
miRFP713           ------------------------------------------------------------    316
miRFP680           ------------------------------------------------------------    316
mRhubarb720        ------------------------------------------------------------    316
mRhubarb719        ------------------------------------------------------------    316
mRhubarb713        ------------------------------------------------------------    316
iRFP720            ------------------------------------------------------------    316
iRFP682            ------------------------------------------------------------    316
iRFP713            ------------------------------------------------------------    316
iRFP713/V256C      ------------------------------------------------------------    316

                                       PHY
BrBphP             WFRPEQVRTVTWGGDPLKAVIIGDDPSDLSPRRSFAQWHQLVEGKSEPWSPAELASARLV    498
mIFP               ------------------------------------------------------------    320
RpBphP1            ------------------------------------------------------------    315
miRFP              ------------------------------------------------------------    313
miRFP2             ------------------------------------------------------------    313
miRFP709           ------------------------------------------------------------    315
miRFP703           ------------------------------------------------------------    315
miRFP670           ------------------------------------------------------------    315
emiRFP703          ------------------------------------------------------------    310
emiRFP670          ------------------------------------------------------------    310
DrBphP-PCM         WLRPELRLEVAWGGATPDQA-----KDDLGPRHSFDTYLEEKRGYAEPWHPGEIEEAQDL    494
WiPhy              ------------------------------------------------------------    335
SNIFP              ------------------------------------------------------------    320
iFP1.4             ------------------------------------------------------------    321
iFP2.0             ------------------------------------------------------------    321
ReBphP-PCM         LFRQELVRTVRWGGDPHKPVEYGPNGPRLTPRKSFEEWSELVRGRSLPFTEAEQRVAETI    490
RpBphP6            ------------------------------------------------------------    311
iRFP702            ------------------------------------------------------------    311
iRFP670            ------------------------------------------------------------    311
miRFP702           ------------------------------------------------------------    311
iRFP670-2          ------------------------------------------------------------    311
Agp1               WFRPEVIKTVRWGGDPHKTVQE---SGRIHPRKSFEIWKEQLRNTSFPWSEPELAAAREL    481
RpBphP2            ------------------------------------------------------------    316
miRFP682           ------------------------------------------------------------    316
miRFP720           ------------------------------------------------------------    316
miRFP713           ------------------------------------------------------------    316
miRFP680           ------------------------------------------------------------    316
mRhubarb720        ------------------------------------------------------------    316
mRhubarb719        ------------------------------------------------------------    316
mRhubarb713        ------------------------------------------------------------    316
iRFP720            ------------------------------------------------------------    316
iRFP682            ------------------------------------------------------------    316
iRFP713            ------------------------------------------------------------    316
iRFP713/V256C      ------------------------------------------------------------    316
```

```
                                     PHY
BrBphP           SETVADVALQLRSVRMVIAQDQLATISAQVLRSEQPVIIADVEGRILLMNEAFEQQLRAS    558
mIFP             ------------------------------------------------------------    320
RpBphP1          ------------------------------------------------------------    315
miRFP            ------------------------------------------------------------    313
miRFP2           ------------------------------------------------------------    313
miRFP709         ------------------------------------------------------------    315
miRFP703         ------------------------------------------------------------    315
miRFP670         ------------------------------------------------------------    315
emiRFP703        ------------------------------------------------------------    310
emiRFP670        ------------------------------------------------------------    310
DrBphP-PCM       RDTLTGAL----------------------------------------------------    502
WiPhy            ------------------------------------------------------------    335
SNIFP            ------------------------------------------------------------    320
iFP1.4           ------------------------------------------------------------    321
iFP2.0           ------------------------------------------------------------    321
ReBphP-PCM       RVTLIEVVLRLTDEASM---------ARQMANERQELLIA---------------ELNLE    526
RpBphP6          ------------------------------------------------------------    311
iRFP702          ------------------------------------------------------------    311
iRFP670          ------------------------------------------------------------    311
miRFP702         ------------------------------------------------------------    311
iRFP670-2        ------------------------------------------------------------    311
Agp1             RGAIIGIVLRKTEEMAD--------LTRELQRTNKELEAF---------------SYSVS    518
RpBphP2          ------------------------------------------------------------    316
miRFP682         ------------------------------------------------------------    316
miRFP720         ------------------------------------------------------------    316
miRFP713         ------------------------------------------------------------    316
miRFP680         ------------------------------------------------------------    316
mRhubarb720      ------------------------------------------------------------    316
mRhubarb719      ------------------------------------------------------------    316
mRhubarb713      ------------------------------------------------------------    316
iRFP720          ------------------------------------------------------------    316
iRFP682          ------------------------------------------------------------    316
iRFP713          ------------------------------------------------------------    316
iRFP713/V256C    ------------------------------------------------------------    316
```


1 Boitreaud, J. *et al.* (2024). https://doi.org:10.1101/2024.10.10.615955

2 Passaro, S. *et al.* Boltz-2: Towards Accurate and Efficient Binding Affinity Prediction. *bioRxiv* (2025). https://doi.org:10.1101/2025.06.14.659707

3 Schmidt, A. *et al.* Structural snapshot of a bacterial phytochrome in its functional intermediate state. *Nat Commun* **9**, 4912 (2018). https://doi.org:10.1038/s41467-018-07392-7

4 Weiner, S. J. *et al.* A new force field for molecular mechanical simulation of nucleic acids and proteins. *Journal of the American Chemical Society* **106**, 765-784 (2002). https://doi.org:10.1021/ja00315a051

5 Modi, V., Donnini, S., Groenhof, G. & Morozov, D. Protonation of the Biliverdin IXalpha Chromophore in the Red and Far-Red Photoactive States of a Bacteriophytochrome. *J Phys Chem B* **123**, 2325-2334 (2019). https://doi.org:10.1021/acs.jpcb.9b01117

6 Nielsen, J. E. & Vriend, G. Optimizing the hydrogen-bond network in Poisson-Boltzmann equation-based pK(a) calculations. *Proteins* **43**, 403-412 (2001). https://doi.org:10.1002/prot.1053

7 Vedani, A. & Huhta, D. W. Algorithm for the systematic solvation of proteins based on the directionality of hydrogen bonds. *Journal of the American Chemical Society* **113**, 5860-5862 (2002). https://doi.org:10.1021/ja00015a049

8 Neese, F. Software update: The ORCA program system—Version 5.0. *WIREs Computational Molecular Science* **12** (2022). https://doi.org:10.1002/wcms.1606

9 Weigend, F. & Ahlrichs, R. Balanced basis sets of split valence, triple zeta valence and quadruple zeta valence quality for H to Rn: Design and assessment of accuracy. *Phys Chem Chem Phys* **7**, 3297-3305 (2005). https://doi.org:10.1039/b508541a

10 Perdew, J. P., Burke, K. & Ernzerhof, M. Generalized Gradient Approximation Made Simple. *Phys Rev Lett* **77**, 3865-3868 (1996). https://doi.org:10.1103/PhysRevLett.77.3865

11 Caldeweyher, E. *et al.* A generally applicable atomic-charge dependent London dispersion correction. *J Chem Phys* **150**, 154122 (2019). https://doi.org:10.1063/1.5090222
12 Stoychev, G. L., Auer, A. A. & Neese, F. Automatic Generation of Auxiliary Basis Sets. *J Chem Theory Comput* **13**, 554-562 (2017). https://doi.org:10.1021/acs.jctc.6b01041
13 Scheurer, M. *et al.* PyContact: Rapid, Customizable, and Visual Analysis of Noncovalent Interactions in MD Simulations. *Biophys J* **114**, 577-583 (2018). https://doi.org:10.1016/j.bpj.2017.12.003
14 Grigorenko, B. L., Polyakov, I. V. & Nemukhin, A. V. Modeling photophysical properties of the bacteriophytochrome-based fluorescent protein IFP1.4. *J Chem Phys* **154**, 065101 (2021). https://doi.org:10.1063/5.0026475
15 Steiner, T. The hydrogen bond in the solid state. *Angew Chem Int Ed Engl* **41**, 49-76 (2002). https://doi.org:10.1002/1521-3773(20020104)41:1<48::aid-anie48>3.0.co;2-u
16 . https://doi.org:10.1021/ja9756331
